# Molecular de-extinction of antibiotics enabled by deep learning

**DOI:** 10.1101/2023.10.01.560353

**Authors:** Fangping Wan, Marcelo D. T. Torres, Jacqueline Peng, Cesar de la Fuente-Nunez

**Author notes:** These authors contributed equally to this work (Fangping Wan, Marcelo D. T. Torres).

## Abstract

Molecular de-extinction is an emerging field that aims to resurrect molecules to solve present-day problems such as antibiotic resistance. Here, we introduce a deep learning approach called Antibiotic Peptide de-Extinction (APEX) to mine the proteomes of all available extinct organisms (the “extinctome”) searching for encrypted peptide (EP) antibiotics. APEX mined a total of 10,311,899 EPs and identified 37,176 sequences predicted to have broad-spectrum antimicrobial activity, 11,035 of which were not found in extant organisms. Chemical synthesis and experimental validation yielded archaic EPs (AEPs) with activity against dangerous bacterial pathogens. Most peptides killed bacteria by depolarizing their cytoplasmic membrane, contrary to known antimicrobial peptides, which target the outer membrane. Notably, lead peptides, including those derived from the woolly mammoth, ancient sea cow, giant sloth, and extinct giant elk, exhibited anti-infective activity in preclinical mouse models. We propose molecular de-extinction, accelerated by deep learning, as a framework for discovering therapeutic molecules.

## Introduction

Biological molecules serve as invaluable records of evolutionary history^1^ and may be able to provide blueprints for therapeutic design. We recently introduced the term molecular de-extinction^2^, referring to the resurrection of extinct molecules of life to tackle contemporary challenges. By uncovering a new sequence space of previously unexplored molecules, molecular de-extinction offers a promising approach to expand our vision of life’s molecular diversity while helping unveil molecules that may play a role in host immunity throughout evolution. Specifically, this study proposes molecular de-extinction as a framework for drug discovery, aiming to address the urgent global health issue of antimicrobial-resistant (AMR) infections.

With AMR infections causing approximately 1.27 million deaths annually worldwide, and projections indicating a potential 10 million annual fatalities by 2050^3^ in the absence of effective new drugs, urgent measures are required to combat antibiotic resistance. Furthermore, according to the World Health Organization, by 2030, around 24 million individuals could face extreme poverty due to the high cost of treating these infections^3^.

Computational approaches have been developed for the design and discovery of peptide antibiotics^4,5^. For example, machine learning (ML) models have been used to generate peptide sequences^6,7^ and to predict antimicrobial activity^8,9^, hemolysis^10^, and AMR^11,12^. Recently, computational methods have been developed to discover new antibiotics through proteome mining^2,13^. We previously performed a proteome-wide exploration of the human body and identified encrypted peptides (EPs), fragments within a protein sequence that possess antimicrobial properties^13^. We hypothesized that EPs exist not only in modern humans but also throughout evolution. Thus, subsequently, through paleoproteome mining, we extended this approach to identify similar molecules in ancient humans^2^.

In the present work, we introduce **A**ntibiotic **P**eptide de-**Ex**tinction (APEX, **Fig. 1**), a new multitask deep learning (DL) approach. Using APEX, we systematically mined all available proteomes of extinct organisms (the “extinctome”) to discover potential antimicrobial peptides. This effort led to the identification of 37,176 EPs with predicted antibiotic activity (**Data S1**). Of these peptides, 11,035 were classified as archaic EPs (AEPs), meaning they were absent in extant organisms, while the rest, referred to as modern EPs (MEPs), were present in both extant and extinct organisms. These peptides (AEPs and MEPs) were computationally predicted to exhibit antimicrobial activity based on our broad-spectrum classification threshold (median MIC ≤80 μmol L^-1^).

**Figure 1.**
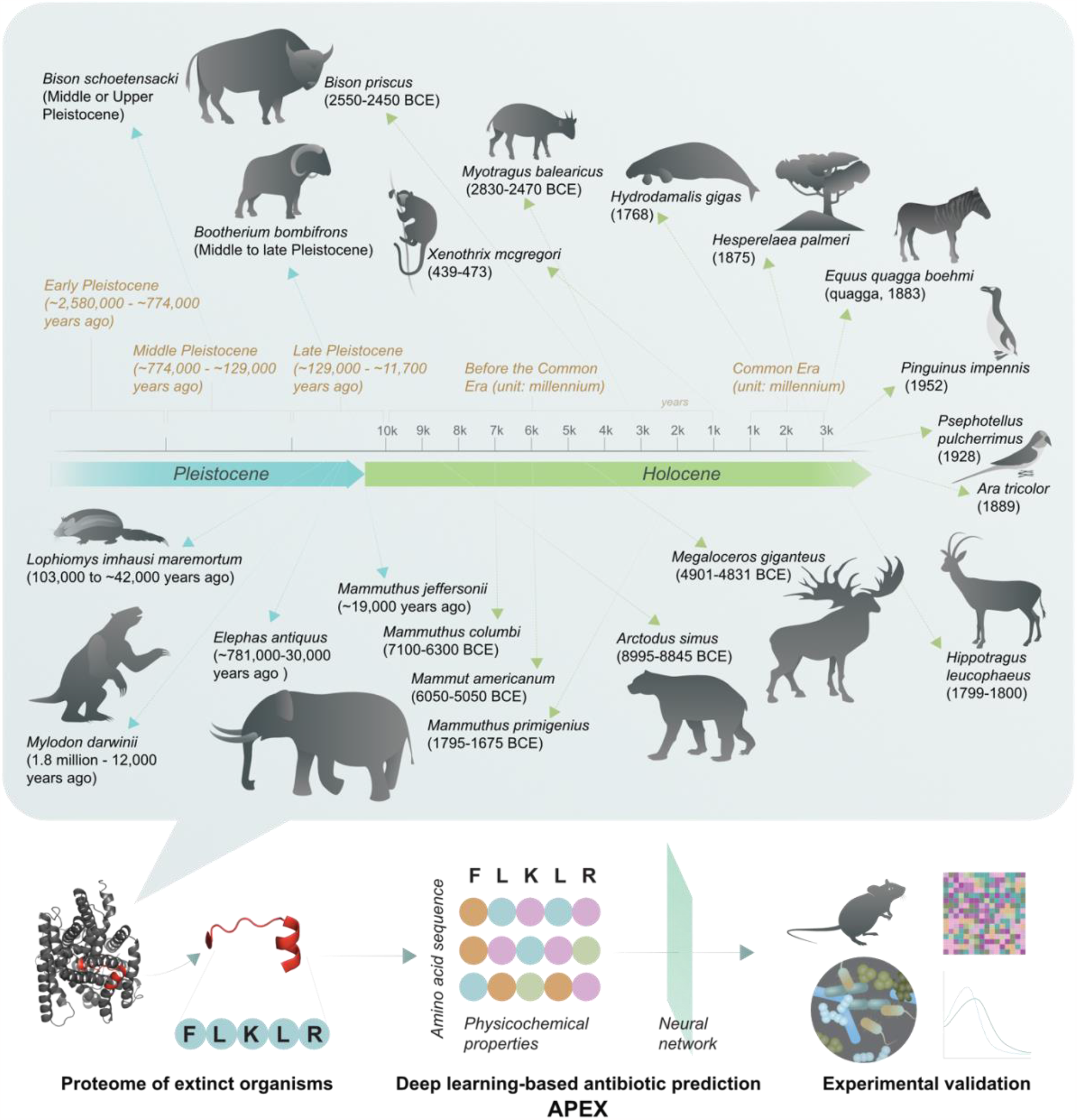
Molecular de-extinction of antibiotics from ancient proteomes using deep learning. All available proteomes of extinct organisms were mined by APEX, our deep learning algorithm. Amino acid sequences ranging from 8 to 50 amino acid residues within proteins from extinct organisms were inputted into a multitask deep learning model that trained on both public and in-house peptide data to evaluate the potential antimicrobial activity. The highest ranked peptides based on predicted antimicrobial activities were then selected and thoroughly characterized against clinically relevant pathogens both *in vitro* and in animal models. The mechanism of action, physicochemical features, and synergistic interactions of these peptides were also assayed. The dates reported the extinction date or period for the organisms studied. The protein and peptide structures shown in the figure were created with PyMOL Molecular Graphics System, Version 2.1 Schrödinger, LLC.

To validate their functionality, we synthesized a diverse set of 69 EPs (**Data S2**), comprising 20 AEPs and 49 MEPs, covering a wide range of peptide sequences to ensure diversity. These peptides were subjected to thorough *in vitro* characterization, including assessment of antimicrobial activity, mechanism of action, secondary structure, and synergy. Our synthesis strategy ensured that no more than three peptides from the same extinct organism were included, both for AEPs and MEPs, to broaden organismal coverage. The assays yielded 13 AEPs and 28 MEPs that displayed antimicrobial activity *in vitro*.

To expand our analysis to *in vivo* settings, we evaluated three AEPs and four MEPs in two preclinical mouse models infected with the major Gram-negative pathogen *Acinetobacter baumannii*. Notably, two AEPs and one MEP exhibited anti-infective activity comparable to polymyxin B, a widely used antibiotic, under physiologically relevant conditions^14^. These findings demonstrate the potential of the identified peptides as effective antimicrobial agents.

This study illustrates the power of deep learning in propelling the field of molecular de-extinction and its potential as a drug discovery framework. The findings also highlight the valuable insights that can be gained from molecules of the past, which hold promise for addressing present-day challenges and benefiting us in the present.

## Results and Discussion

### Antimicrobial activity prediction from sequence using deep learning

Recent advances have enabled the exploration of proteomes for antibiotic discovery^2,13^. To enhance these efforts, we have developed APEX, a deep learning (DL) model that employs a multitask learning architecture to predict whether EPs have antimicrobial activity (**Fig. S1**). APEX was trained on peptide data from both our in-house dataset and from the Database of Antimicrobial Activity and Structure of Peptides (DBAASP), a publicly available database^15^, to extract hidden features of peptide sequences. APEX utilizes an encoder neural network, combining recurrent and attention neural networks (**Fig. S1**), to extract hidden features from peptide sequences. The encoder neural network was then coupled with multiple downstream neural networks to predict antimicrobial activity according to the source of the peptide (*i*.*e*., from in-house or public datasets). In our platform, the extracted hidden features were fed into two separate, fully connected neural networks (FCNNs): one neural network was trained on our in-house peptide dataset and used to predict antimicrobial activity against specific bacterial strains (*i*.*e*., a regression problem); the other neural network was trained on publicly available antimicrobial peptides (AMPs) and inactive peptides (referred here as non-AMPs) derived from the DBAASP dataset^15^ to perform a binary classification (*i*.*e*., a classification problem). We defined as non-AMPs those peptides that were not active at the range of concentrations selected as threshold, *i*.*e*., MIC >30 μmol L^-1^ (***Publicly available AMP sequences*** in ***Methods***). Any publicly available peptide sequences that overlapped with our in-house dataset were removed from the model training to prevent label information leakage. Since the encoder neural network was trained on both the in-house and public datasets, the incorporation of the latter FCNN served as a data augmentation strategy to improve prediction performance.

To train our APEX model, we utilized a combination of 988 in-house peptides and 5,093 and 5,500 publicly available AMPs and non-AMPs, respectively, obtained from the DBAASP^15^. Our in-house dataset included 14,738 antimicrobial activity data values obtained from 34 bacterial strains. To assess the antimicrobial prediction performance of APEX, we randomly split our in-house dataset into a cross validation (CV) set and an independent set, consisting of 790 and 198 peptides, respectively. Five-fold CV was first used to tune the hyperparameters on the CV set, while the independent set was used to evaluate the final prediction performance of ML models trained on the CV set with determined hyperparameters.

To compare the performance of our DL approach with simple ML predictors, we implemented several baseline ML models, including elastic net, linear support vector regression, extra-trees regressor, random forest, and gradient boosting decision tree, and trained and evaluated them on the same datasets. The hyperparameter ranges searched for each ML model are provided in **Tables S1-S4**. On the CV set, our APEX model with the best hyperparameter combination outperformed all baseline ML models in terms of predicted activity for most bacteria, focusing particularly on 11 bacterial pathogens known as the ESKAPEE pathogens. These pathogens are classified by the World Health Organization as the most dangerous threats to our society, and include *Enterococcus faecium, Staphylococcus aureus, Klebsiella pneumoniae, A. baumannii, P. aeruginosa, Enterobacter* spp., and *Escherichia coli*)^16^.

Specifically, APEX outperformed all baseline ML models on most pathogen-specific MIC predictions in terms of R-squared Pearson, and Spearman correlations (*i*.*e*., single APEX in **Figs. S2-S7** and **Tables S5-S7)**. The average R-squared, Pearson, and Spearman correlations of the baseline ML methods were, at best, 0.378, 0.584, and 0.523, respectively. Compared to the baseline, the single best obtained similar R-squared = 0.369, and better Pearson correlation = 0.621, and Spearman correlation = 0.556 (**Tables S5-S7**). To improve the prediction performance, we adopted an ensemble learning approach by selecting the top eight APEX models (with different neural network architectures and training strategies; for details, see subtopic “***Hyperparameter tuning, model evaluation and ensemble learning***” in the **Methods** section, **Figs. S8-S10**) ranked by the average R-squared on the CV set and obtained the final predictions by averaging the predictions from these models. The ensemble learning approach (*i*.*e*., ensemble APEX v1) increased the prediction performance to 0.473, 0.669, and 0.594 in terms of R-squared, Pearson correlation, and Spearman correlation, respectively (**Figs. S2-S7** and **Tables S5-S7**). APEX involved the following multitask training steps: (1) using a single FCNN to simultaneously predict the antimicrobial activities of the peptides for the 34 strains from the in-house collection; (2) augmenting the training data by incorporating another FCNN to predict whether peptides from public databases (either AMPs or non-AMPs) are antimicrobials; and (3) we imposed a multitask training constraint on the learnable weights of the last layer in the species-specific antimicrobial prediction FCNN. Briefly, this last constraint encouraged the model to give similar prediction results for similar bacteria (defined by having shorter phylogenetic distance from each other). To evaluate the effectiveness of adding publicly available AMPs/non-AMPs data into our training as well as that of using the multitask training constraint, we conducted an ablation study by dropping these two parts during training and evaluated the corresponding prediction performance on the CV set. We observed that dropping the publicly available AMPs/non-AMPs data from the training set significantly decreased prediction performance (**Figs. S11-S13 and Tables S8-S10**). The addition of the multitask training constraint led to either increased or decreased prediction performance depending on the target bacterial strain (**Figs. S11-S13 and Tables S8-S10**). Thus, we treated the presence or absence of the multitask training constraint as an additional hyperparameter, and we allowed subsequent hyperparameter tuning to decide whether to use the constraint or not. Of note, among the top eight selected APEX models, six used the constraint during model training (**Table S11**).

Because the model selection was based on the CV set, evaluating the prediction performance on the CV set alone may overestimate the generalization ability of APEX and the baseline models. Therefore, we trained the ML models with the determined hyperparameters on the whole CV set and evaluated their prediction performance on the independent set. Similar to the results obtained on the CV set, ensemble APEX v1 achieved an R-squared value of 0.520, a Pearson correlation of 0.706, and a Spearman correlation of 0.582 on average (**Fig. 2a, Figs. S14-S18** and **Tables S12-S14**), outperforming all baseline ML methods. In practice, to make prediction results more robust, a single ML model may be trained with different random seeds. We averaged the prediction results from all model copies, to counter the potential stochastic behavior caused by the choice of random seeds. For each APEX model we selected, we trained five copies with different random seeds and created a second ensemble learning version (ensemble APEX v2) with 40 APEX models (i.e., eight APEX models × five copies). This ensemble learning approach increased the prediction performance to 0.546, 0.728 and 0.607 in terms of R-squared, Pearson correlation, and Spearman correlation, respectively (**Fig. 2a, Figs. S14-S21** and **Tables S12-S14**).

**Figure 2.**
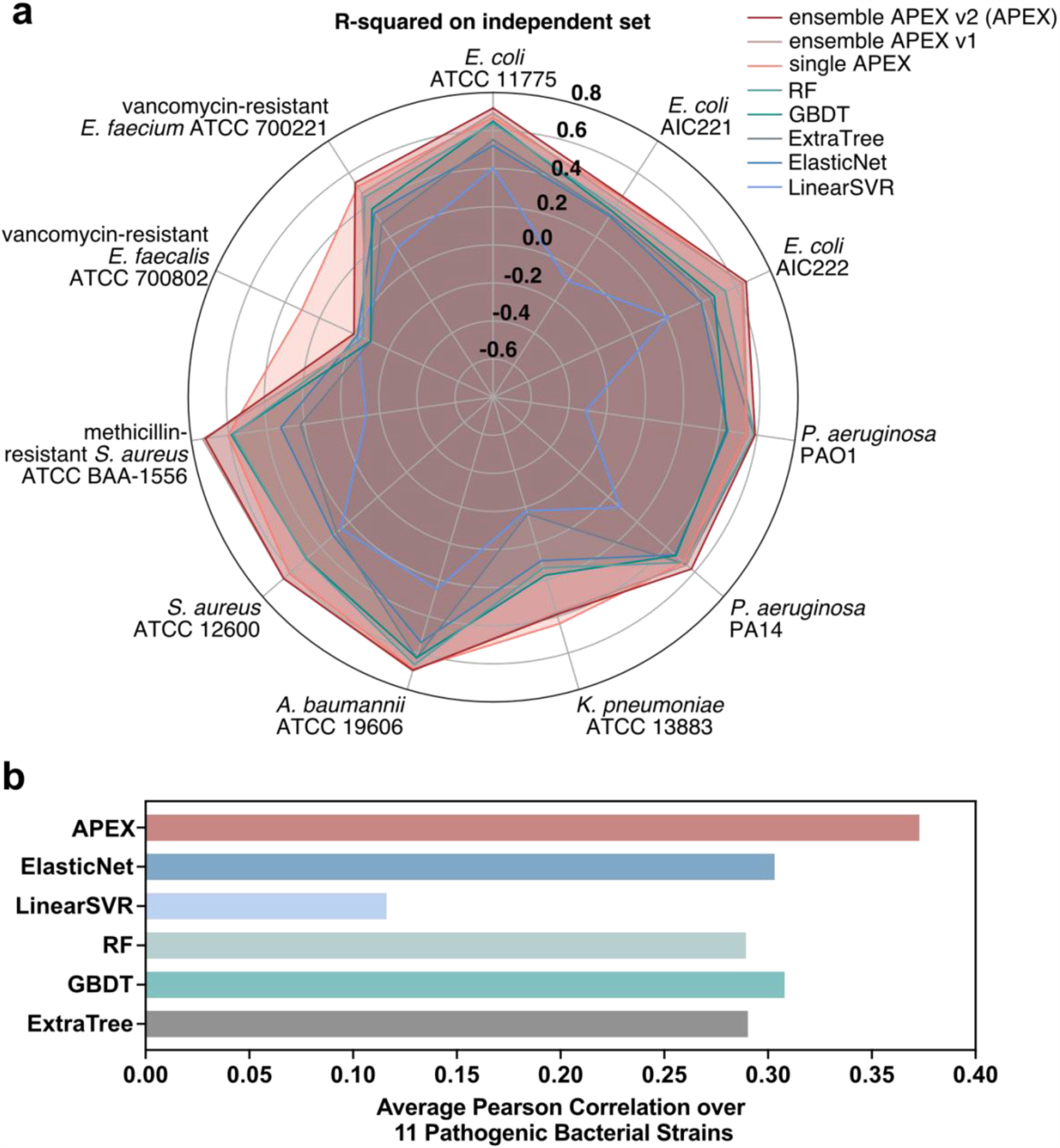
APEX prediction performance and comparison with other models. **(a)** Radar chart showing R-squared correlation in terms of species-specific antimicrobial activity prediction on an independent dataset (a held-out subset from our in-house peptide dataset) for various machine learning (ML) models. The radius reflects the R-squared value for each of the models. APEX variants outperformed the baseline ML methods with most of the pathogens analyzed. RF: random forest; GBDT: gradient boosting decision tree; ExtraTree: extra-tree regressor; ElasticNet: elastic net; LinearSVR: linear support vector regression. **(b)** Mean of species-wise Pearson correlation of log_2_-transformed MICs between values obtained experimentally and predicted by various ML models, evaluated dataset: 69 peptides we synthesized and tested.

We then tested the predictive power of APEX compared to that of a scoring function used previously to discover EPs in the human proteome^13^. Given a peptide sequence, the scoring function^17^ used hydrophobicity and net charge to compute a predictive score of antimicrobial potential. Since the 56 human EPs validated experimentally for antimicrobial activity in our previous work^13^ are part of our in-house dataset, we used them here as the test set and used the rest of our in-house dataset (932 peptides) for model training and selection. For the dataset consisting of the 56 validated human EPs, the ensemble APEX v2 achieved highest values for Pearson and Spearman correlations in most cases (**Figs. S22-S25** and **Tables S15-S16**). Lastly, during the subsequent prediction of the antimicrobial activities for the 69 synthesized EPs using ML models trained on the entire in-house dataset, APEX outperformed all the baseline ML methods. Notably, APEX achieved the highest Pearson correlation for the MIC prediction (**Fig. 2b**, more details on ***In vitro antimicrobial activity of encrypted peptides from extinct organisms***). Collectively, these results substantiate our computational validation of APEX as the most accurate model for antimicrobial activity prediction in comparison to all the other models tested by us (**Fig. 2a, b and Figs. S2-25**). Based on these results, we decided to use the ensemble APEX v2 model (hereafter referred to as APEX for simplicity) to mine extinct proteomes for EPs.

### Differences between modern and archaic encrypted peptides

To investigate the differences between modern EPs (MEPs) and archaic EPs (AEPs) and determine if these sequences represent potential novel classes of antimicrobial peptides, we gathered 12,860 protein sequences from 208 extinct species obtained from NCBI. After removing redundant sequences, we were left with 5,190 proteins. EPs were selected based on specific criteria, including the length of substrings ranging from 8-50 amino acid residues within these protein sequences, which align with the lengths of most active antimicrobial peptides reported previously^13^. Briefly, using our APEX model, we first predicted the antimicrobial activity of the resulting 10,311,899 peptide sequences derived from the 5,190 proteins of the explored extinct species (**Fig. 3a**). As APEX is built to predict species-specific antimicrobial activities, there are multiple ways to select EPs for downstream validation. Here, we ranked the 10,311,899 EPs by median antimicrobial activities (i.e., broad spectrum EPs), or selectivity against Gram-positive or Gram-negative pathogens. For each ranked list, we used the following criteria to filter out EPs: (i) length not ranging from 8 to 30 amino acid residues, (ii) sequences that are present in our in-house dataset, (iii) with high sequence similarity to AMPs from public databases, and (iv) EPs that are present in the modern human proteome. For each resulting list, we grouped the EPs by their source organisms and selected the top ranked EPs that were not too hydrophobic to be chemically synthesized by solid-phase peptide synthesis. To ensure sequence novelty, we selected EP sequences from each list that were not too similar to each other. Detailed selection and filtering criteria can be found in **Methods** subtopic ***Encrypted peptide screening and selection from extinct proteomes***.

**Figure 3.**
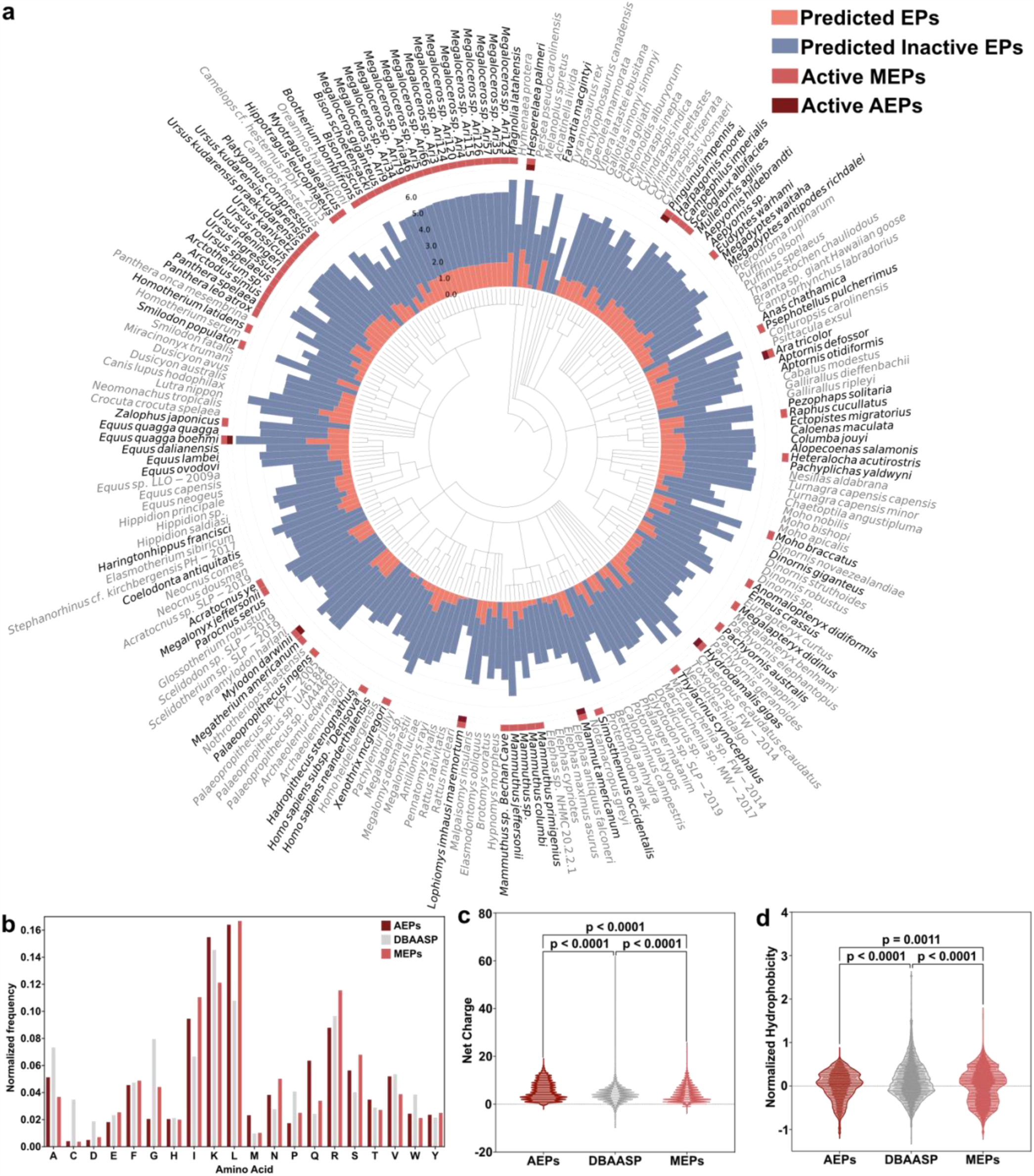
Antimicrobials identified by APEX in extinct organisms and their composition and physicochemical properties. **(a)** Phylogenetic tree showing the extinct organisms scanned by APEX. Circular bars denote the log_10_-transformed average active (red) and inactive (blue) encrypted peptides discovered by APEX. The values were normalized by the number of proteins per organism scanned. The organisms whose encrypted peptides were selected for validation are highlighted in bold type. Extinct organisms that presented active encrypted peptides (EPs) validated experimentally are indicated by a light red square and, within that group, those organisms encoding extinct sequences absent in extant organisms are highlighted with a dark red square. **(b)** Amino acid frequency in AEPs and MEPs compared with known AMPs from the DBAASP database. AEPs present a higher frequency of the basic residue K, the aliphatic residue V, and uncharged polar residues (M, Q, and T) than MEPs. **(c-d)** The distribution of two physicochemical properties of peptides with antimicrobial activity (AEPs, MEPs and AMPs from the DBAASP): net charge **(c)**; and normalized hydrophobicity **(d)**. Net charge directly influences the initial electrostatic interactions between the peptide and negatively charged bacterial membranes, and hydrophobicity directly influences the interactions of the peptide with lipids in the membrane bilayers. Encrypted peptides from extinct organisms are slightly less hydrophobic than encrypted peptides or peptides from the DBAASP and have a net positive charge.

We then compared the differences between peptide sequences identified with APEX and those identified with our earlier scoring function^13^. First, for a direct comparison, we used APEX to find peptide sequences within proteins from the modern human proteome, extracted these sequences, and determined their amino acid compositions (**Fig. 3b and Fig. S26**). In contrast to the scoring function, which takes into consideration net positive charge to identify sequences^13^, APEX selected sequences in which negatively charged residues (i.e., aspartic acid and glutamic acid), as well as glycine and polar uncharged residues (i.e., asparagine, glutamine, and serine), were overrepresented. These are amino acids residues with net charge and hydrophobicity features that are not relevant to the scoring function, which mostly considers net positive charge and amphipathic peptide sequences. The EPs identified by the scoring function also showed a higher content of cysteine, methionine, phenylalanine, and arginine. Interestingly, lysine, which is preferentially scored by the scoring function, was slightly overrepresented in APEX-identified sequences (**Fig. S26**).

Furthermore, we compared the amino acid composition of EPs identified by APEX to conventional AMPs from the DBAASP (**Fig. 3b**). Generally, EPs identified by APEX presented lower cysteine, aspartic acid, and glycine content compared to AMPs from the DBAASP (**Fig. S27**). EPs derived from proteins of extinct organisms also had a lower asparagine and higher methionine and glutamine content compared to AMPs from the DBAASP (**Fig. S28**). The MEPs identified by APEX had a much lower alanine, proline, and tryptophan content but a much higher isoleucine, leucine, asparagine, and serine content than peptides in the DBAASP (**Fig. S29**). Comparative analysis between AEP and MEP sequences identified by APEX revealed an overrepresentation of methionine and glutamine in the AEPs and of glycine in the MEPs (**Fig. S30**).

When the physicochemical features that contribute to antimicrobial properties^15^ were compared (**Fig. S31**), AEPs showed a lower amphiphilicity (amphiphilicity index <2; **Fig. S31a**) than MEPs or AMPs (**Fig. S31b**) but a slightly higher propensity to be disordered (disordered conformation propensity score from -0.5 to 1). These results indicate that the interactions between the AEPs and the bacterial membrane are likely to differ from those of standard AMPs, which are more amphiphilic and tend to assume a defined structure upon contact with the lipid from the membrane bilayer^15,18^ (**Fig. S31a-b and Fig. S32**). Additionally, to determine the potential toxicity and amphiphilicity of the peptides^18^, we assessed their theoretical tendency toward aggregation (**Fig. S31c)** and the angle of the hydrophobic residues upon adopting a secondary structure (**Fig. S31d)**. These physicochemical parameters are predictive of how the peptides interact with membrane lipids to exert antimicrobial activity^13^. Interestingly, when comparing AEPs with either MEPs or AMPs from the DBAASP, we found that AEPs were less prone to aggregate (*in vitro* aggregation propensity score <500) and presented a smaller predicted hydrophobic face (<100°) (**Fig. S31c-d**). These results are a direct consequence of the higher frequency of uncharged polar residues in AEPs. To investigate peptide structure further, we obtained the predicted normalized hydrophobic moment (**Fig. S31e)** and isoelectric point of the AEPs (**Fig. S31f**), which presented a low range of normalized hydrophobic moment (0-0.6) and which clustered within a short isoelectric point range (9.5 to 13). These values, found for sequences in extinct organisms (AEPs), overlapped with those found for sequences in extinct and extant organisms (MEPs) as well as in AMPs from the DBAASP (**Fig. S31e-f**). The values aligned with the lower abundance of acidic residues compared to basic ones, particularly lysine, in the AEPs (**Fig. 3c-d**).

Collectively, the AEPs identified by APEX represent a distinct family of peptides with a higher abundance of uncharged polar residues and an increased aliphatic content (particularly isoleucine and leucine) with respect to other classes of peptide antibiotics, such as other EPs^13,19,20^ or AMPs^21^. There are a few AMP families, such as he-1, brevenins, pleurains, and frog defensins^21^, having a lower net charge than most standard AMPs, which have more uncharged polar residues or a balance of positively charged and acidic residues, and whose antimicrobial activity depends on how their electronic density is distributed^16^. Like the AMPs in these families but unlike previously described EPs^13^ (**Fig. S26**) or conventional AMPs (**Fig. S27**), AEPs have a high abundance of uncharged polar residues^18^. Leucine and isoleucine, in particular, are structurally important: the stiffness of these bulky branched residues limits the internal flexibility of the peptide, whereas other aliphatic residues favor specific foldamers during the folding process^22^. The difference between the amino acid composition of known AMPs and that of AEPs and MEPs results in significantly different physicochemical features (**Fig. 3c-d** and **Fig. S31**), reaffirming that the EPs identified by APEX are different from known antimicrobial peptides.

### In vitro antimicrobial activity of encrypted peptides from extinct organisms

To further validate the predictive accuracy of APEX in identifying active EPs from extinct organisms, we synthesized and tested two non-overlapping sets of peptides: (i) 49 EPs predicted by our scoring function (**Fig. S33, Data S3**) and (ii) 69 EPs predicted by APEX (**Fig. 3a-b, Data S2**). While the 49 EPs predicted by the scoring function occurred in both extinct and extant organisms (*i*.*e*., all were MEPs), the APEX-predicted EPs included many that were unique to extinct organisms (20 AEPs and 49 MEPs). These peptides were encrypted in proteins found in 98 extinct species. The selection of peptides for synthesis was based on their predicted antimicrobial activities, the source organism, and sequence diversity, ensuring a comprehensive representation (refer to the ***Encrypted peptide screening and selection from extinct proteome*** section in the **Methods** for more information).

Among the EPs identified by APEX, 21 (5 AEPs and 16 MEPs) were derived from secreted proteins, while 48 (15 AEPs and 33 MEPs) were from non-secreted proteins. We included EPs from non-secreted proteins due to the limited annotations of extinct proteins. Filtering out unannotated secreted proteins would have restricted the sequence space explored, so we considered EPs from both secreted and non-secreted proteins. Out of the 21 peptides from secreted proteins identified by APEX, 4 were predicted to target Gram-positive bacteria selectively, 10 to target Gram-negative bacteria selectively, and 7 to exhibit broad-spectrum activity. Among the 48 peptides selected from non-secreted proteins by APEX, 19 were predicted to selectively target Gram-positive bacteria, 10 to selectively target Gram-negative bacteria, and 19 to display broad-spectrum activity.

Next, we synthesized the 21 AEPs and 48 MEPs identified by APEX from extinct organisms and experimentally determined their MICs for 11 clinically relevant bacterial pathogens (seven Gram-negatives and four Gram-positives), ten of which are on the ESKAPEE pathogen list^16^ (**Fig. 4a-b, Data S2**). All experimentally determined MICs (log_2_ transformed) were compared to predictions generated by APEX, yielding Pearson and Spearman correlation values of 0.448 and 0.404, respectively (**Fig. 5a**), underscoring APEX’s significant predictive power. In terms of species-specific antimicrobial activity prediction, APEX showed a high predicted versus experimentally validated activity correlation (Pearson correlation >0.3) for *A. baumannii* ATCC 19606, *Escherichia coli* strains AIC221, AIC222 (a colistin-resistant strain), and ATCC 11775, *P. aeruginosa* strains PAO1 and PA14, and *E. faecium* ATCC 700221 (a vancomycin-resistant strain). All correlation results obtained for the 11 experimentally validated strains, shown in **Figs. S34-S44**. Furthermore, when the average species-specific Pearson correlations of the log_2_-transformed MIC predictions for the 11 pathogens of various baseline ML models were compared with that given by APEX, APEX gave the most accurate predictions (**Fig. 2b**). Of the 69 synthesized peptides, 41 showed significant antimicrobial activity (i.e., MIC ≤128 μmol L^-1^) against at least one bacterial strain, demonstrating a 59% hit rate for identifying peptides with antimicrobial activity (**Fig. 5b**). This hit rate is much higher than the one found for the scoring function (24%) when it was used for extracting EPs from the same extinct sources (**Fig. 5b**). In addition, 13 of the 41 active EPs found by APEX were AEPs, meaning that they were present in extinct organisms but not in any extant organism.

**Figure 4.**
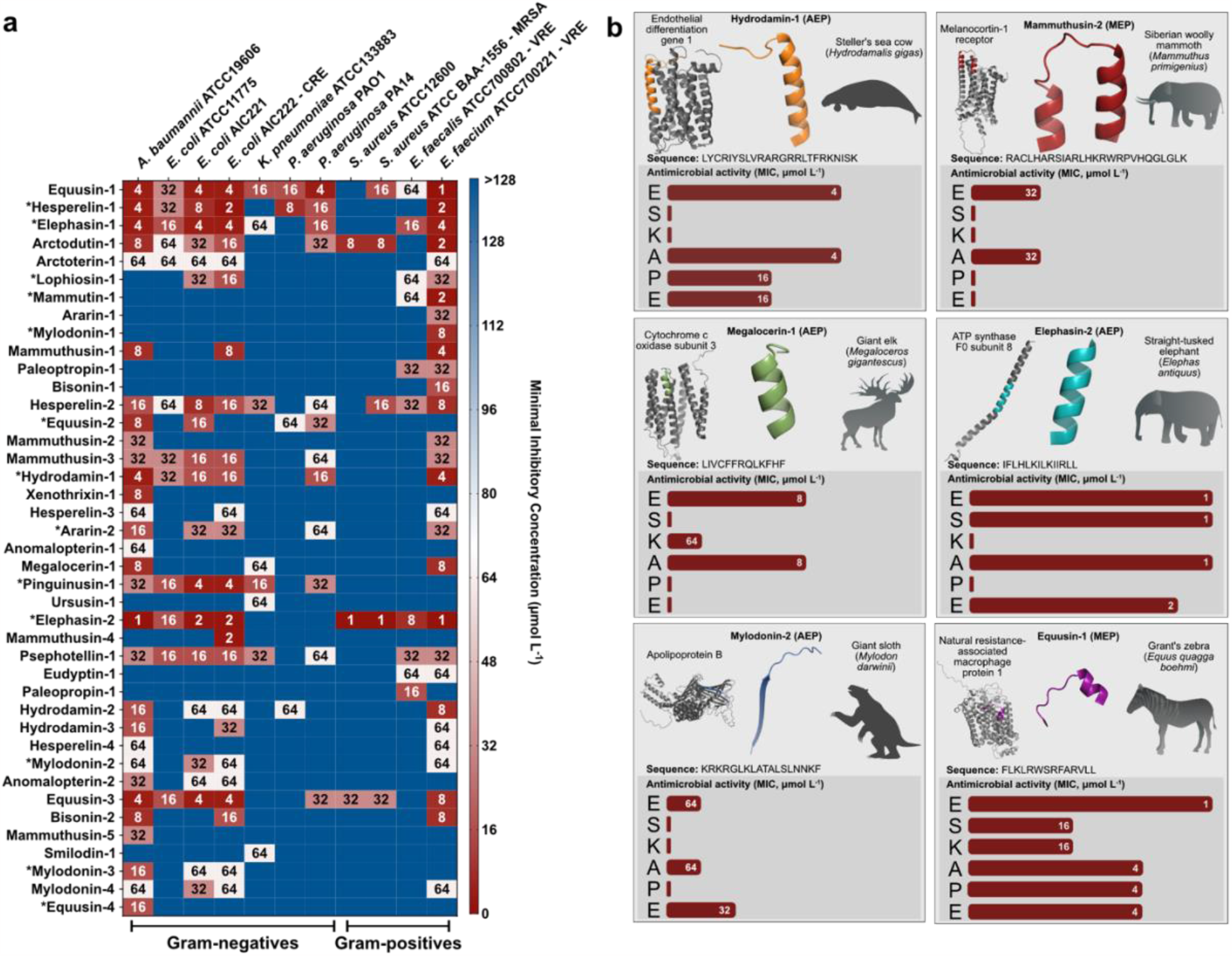
Antimicrobial activity profiles of sequences from the proteomes of extinct organisms. **(a)** Heat map of the antimicrobial activities (μmol L^−1^) of the active encrypted peptides from extinct organisms against 11 clinically relevant pathogens, including four strains resistant to conventional antibiotics. Briefly, 10^6^ bacterial cells and serially diluted encrypted peptides (0-128 μmol L^−1^) were incubated at 37 °C. One day post-treatment, the optical density at 600 nm was measured in a microplate reader to evaluate bacterial growth in the presence of the encrypted peptides from extinct organisms. MIC values in the heat map are the arithmetic mean of the replicates in each condition. **(b)** Examples of active archaic encrypted peptides (AEPs) and modern encrypted peptides (MEPs) from various extinct organisms, their parent protein, and their activity profile against ESKAPE pathogens (*Enterococcus spp*., *S. aureus, K. pneumoniae, A. baumannii, P. aeruginosa, E. coli*). Antimicrobial activity is expressed as the MIC (μmol L^−1^) and activity bars are presented as -log_2_ MIC. ^13^The data for the assays in **a** are the mean and the experiments were performed in three independent replicates. AEPs in **a** are indicated by an asterisk (*). The protein and peptide structures shown in the figure were created with PyMOL Molecular Graphics System, Version 2.1 Schrödinger, LLC.

**Figure 5.**
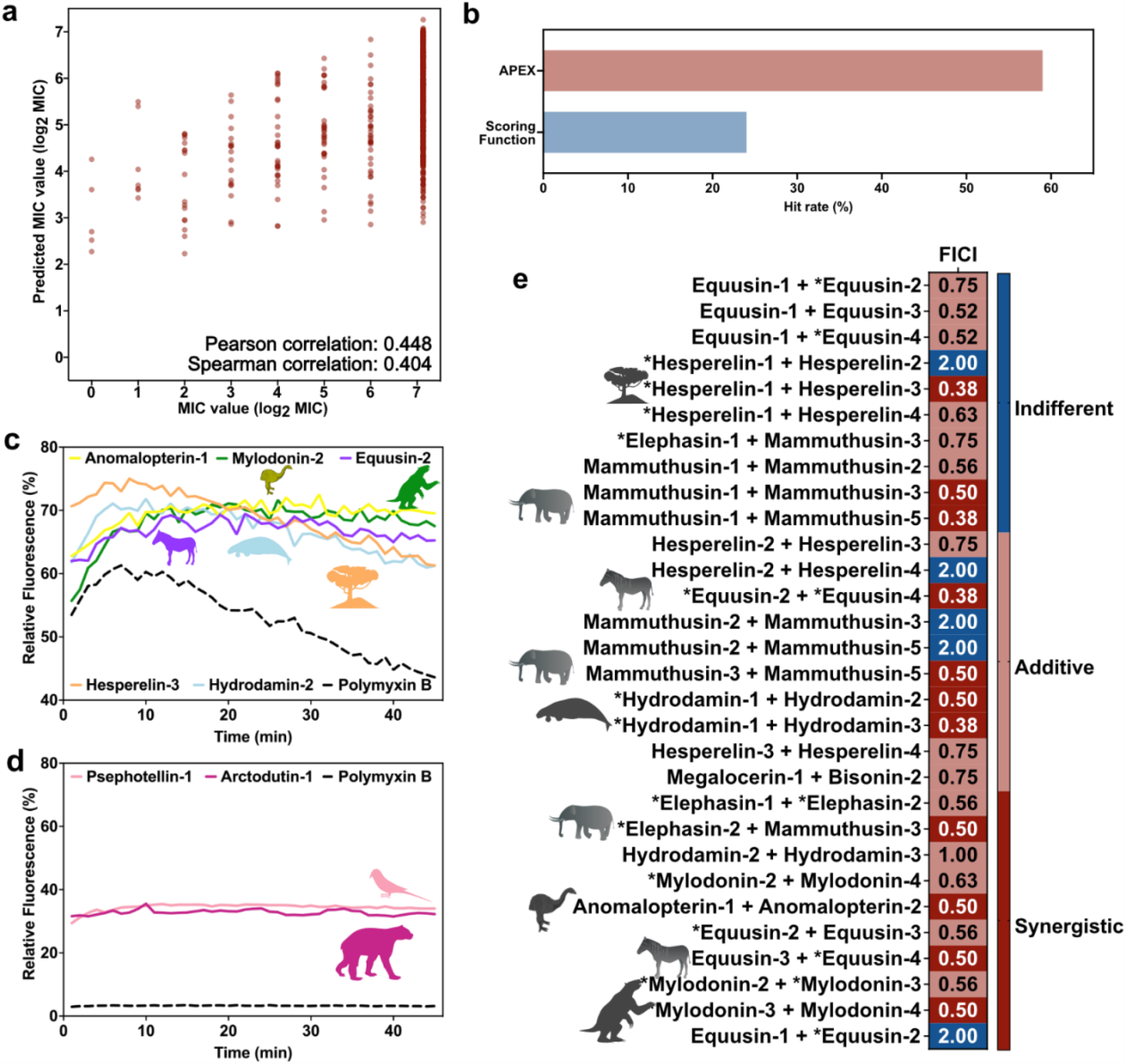
Antimicrobial activity, mechanism of action, and synergy of antimicrobials from the proteomes of extinct organisms. **(a)** Pan-bacterial Pearson and Spearman correlations of log_2_-transformed MICs between experimentally validated values and values predicted by APEX. **(b)** Comparison between the hit rates of APEX and the scoring function, described by Torres *et al*.^13^, previously used to detect encrypted peptides in the human proteome. **(c)** Cytoplasmic membrane depolarization by five encrypted peptides from extinct organisms. The *A. baumannii* membrane was more strongly depolarized by the encrypted peptides than by the antibiotic polymyxin B. **(d)** NPN permeabilization assays showing the effect of two encrypted peptides from extinct organisms on the outer membrane of *A. baumannii*. Higher permeability was observed with the encrypted peptides than with the antibiotic polymyxin B. **(e)** Heat map showing interactions between EPs found by APEX in the same source organisms or multiple source organisms with at least one source in common, expressed as the fractional inhibitory concentration index (FICI). Most of the tested pairs of encrypted peptides from extinct organisms either synergized or had an additive effect on *A. baumannii* and *P. aeruginosa* cells; the latter was only tested against the peptide pair composed of equusin-1 and equusin-2 shown in the last row of the heatmap. The data for the assays in **c, d**, and **e** are the mean and the experiments were performed in three independent replicates. AEPs in **e** are indicated by an asterisk (*).

Next, we used the selectivity score (see ***Selectivity score*** from the **Methods** section for details) to quantify peptide selectivity. Specifically, among the 69 peptides synthesized, 20 were computationally determined to selectively target Gram-negative pathogens and 23 to selectively target Gram-positive pathogens. For peptides predicted to be selective for Gram-negative pathogens, the Pearson correlation of selectivity scores calculated by experimentally validated and predicted MICs was 0.295. In addition, the mean Gram-negative selectivity score derived from experimental MICs for peptides selective for Gram-negatives was 0.783, and 1.33 for the rest of peptides tested. A p-value of 0.013 for the Mann–Whitney U test on the selectivity scores from these two lists suggested a statistically significant difference. This result demonstrated the ability of APEX to discover peptide sequences that selectively target Gram-negative pathogens. On the other hand, we observed a weak Pearson correlation (0.11) between selectivity scores calculated by predicted and validated MIC values of peptides that were selected to target Gram-positive bacteria. The mean selectivity score of these peptides for Gram-positive bacteria was 1.02, indicating that the peptides did not selectively kill Gram-positive pathogens. However, we observed that the remaining peptides had a significantly higher mean selectivity score of 2.13 (p-value = 0.08, Mann–Whitney U test), suggesting that APEX can discover peptides that were relatively not specific against Gram-negative pathogens.

To compare the predictive performance of APEX with our previously described scoring function^13^, we applied the scoring function to identify EPs from secreted proteins of the 208 extinct organisms. As mentioned above, the scoring function ranked and selected 49 MEPs as the top sequences, but it did not identify any AEPs. Subsequently, we synthesized these peptides and experimentally validated their MICs *in vitro* (**Fig. S33, Data S3**). Among the 49 peptides synthesized and tested, 12 displayed antimicrobial activity (*i*.*e*., MIC ≤128 μmol L^-1^) against at least one bacterial strain, demonstrating a 24% EP antibiotic discovery rate by the scoring function (**Fig. 5b**). These results highlight that APEX outperformed our previously reported scoring function, with APEX exhibiting a 2.458-times increase in accuracy in predicting experimentally validated EPs (**Fig. 5b**).

### Secondary structure of encrypted peptides from extinct organisms

Given that the peptides identified by APEX were, on average, different from other EPs predicted by the scoring function and other AMPs in terms of physicochemical descriptors and amino acid residue composition, we decided to determine their secondary structure. When AMPs come into contact with bacterial membranes, they typically adopt an α-helical conformation due to their amphipathicity, net charge, hydrophobicity, and length, all of which directly influence their secondary structure.

To determine the secondary structure of the active EPs obtained from extinct organisms displaying antimicrobial activity, we exposed them to a helix-inducing medium^23^ (trifluoroethanol in water, 3:2, v:v). Interestingly, most of the EPs synthesized and tested that were identified through the scoring function were not α-helical, but instead had a relatively high content of anti-parallel β-structure and were largely unstructured (**Fig. S32a-b**). In contrast, AEPs and MEPs derived from APEX demonstrated predominantly helical structures under the analyzed conditions, despite their unusual abundance of uncharged polar residues and low amphiphilicity (**Fig. S32c-d**). These results highlight the significantly higher success rate achieved by APEX compared to the scoring function. This improvement may be attributed to the greater prevalence of α-helical peptides in nature, as their amphipathic structure enhances peptide-membrane interactions, resulting in more effective membrane damage ^18,24^(**Fig. 5b**).

### Mechanism of action of encrypted peptides

The bacterial membrane is a common target for most known AMPs, where they engage in non-specific interactions with the lipid bilayer^18^. The antimicrobial activity of AMPs is influenced by their amino acid composition, distribution, and various physicochemical characteristics such as amphiphilicity and hydrophobicity. To investigate the underlying mechanisms by which the peptides identified by APEX kill bacteria, we tested whether differences in the composition of the AEPs and MEPs in extinct organisms would affect their mechanism of action. The differences in composition we investigated include the content of uncharged polar and aliphatic residues (**Fig. 3b** and **Figs. S27-S30**) and physicochemical features ranges (**Fig. 3c-d** and **Fig. S31**).

First, we tested whether AEPs and MEPs from extinct organisms depolarized the cytoplasmic membrane of *A. baumannii*. We used the potentiometric fluorophore 3,3′-dipropylthiadicarbocyanine iodide (DiSC_3_-5) whose fluorescence is suppressed by its accumulation and aggregation within the cytoplasmic membrane. In reaction to disturbances in the transmembrane potential of the cytoplasmic membrane, this fluorophore migrates to the outer environment and emits fluorescence. Polymyxin B was used as a positive control in these experiments as it is a depolarizer that also permeabilizes and damages bacterial membranes. Notably, the AEPs and MEPs depolarized the cytoplasmic membrane more effectively than polymyxin B (**Fig. S45**).

The AEPs and MEPs found by APEX in extinct protein sequences depolarized the cytoplasmic membrane more effectively than the peptides found in modern proteins (**Fig. S45**). The most potent depolarization effects were found for the following five peptides: anomalopterin-1, mylodonin-4, equusin-2, hesperelin-3, and hydrodamin-2 (**Fig. 5c**). We hypothesize that this increased depolarization is a consequence of their different amino acid composition, particularly the higher content of long aliphatic residues (*e*.*g*., leucine and isoleucine) in the AEPs and MEPs obtained from APEX compared to known AMPs.

Anomalopterin-1, a peptide originating from the extinct moa species *Anomalopteryx didiformis*, is a fragment of the dynein axonemal heavy chain 3, which forms part of the microtubule-associated motor protein complex. Mylodonin-4, originating from the extinct South American giant sloth *Mylodon darwinii*, corresponds to a fragment of apolipoprotein B, a lipoprotein that functions as a ligand for the low-density lipoprotein (LDL) receptor. Interestingly, EPs derived from the modern human apolipoprotein B have also been described as depolarizers of the membranes of Gram-negative bacterial pathogens^25^. Equusin-2, an AEP originating from the extinct Grant’s zebra *Equus quagga boehmi*, is derived from the abnormal spindle-like microcephaly-associated protein. This protein is responsible for calmodulin-binding activity and plays a role in regulating the meiotic cell cycle, gamete generation, centrosome location maintenance, and nervous system development.

Hesperelin-3 is produced by both an extinct magnolia species (*Magnolia latahensis*) and an extinct palm tree species (*Hesperelaea palmeri*). It is a component of the protein ribulose bisphosphate carboxylase large chain (RuBisCO). RuBisCO catalyzes two reactions: the carboxylation of D-ribulose 1,5-bisphosphate, which is the primary step in carbon dioxide fixation, and the oxidative fragmentation of the pentose substrate in photorespiration. The extinct manatee *Hydrodamalis gigas* yielded a fragment from the von Willebrand factor (hydrodamin-2), a protein involved in hemostasis. EPs derived from the von Willebrand factor have also been previously found in modern humans^13^.

To determine whether the EPs permeabilized the bacterial outer membrane, we performed 1-(N-phenylamino)naphthalene (NPN) assays. NPN, a lipophilic dye, fluoresces faintly in aqueous solutions but fluoresces significantly more when it encounters lipidic environments such as bacterial membranes. NPN can penetrate the bacterial outer membrane only if it is disrupted or compromised. In contrast to cells treated with the positive control polymyxin B or cells left untreated (untreated control group), bacteria exposed to the most active EPs at their MIC were, in general, not effectively permeabilized (**Fig. 5d and Fig. S46**). The EPs that permeabilized the outer membrane, causing a higher increase in the uptake of the fluorescent probe, were the MEP psephotellin-1, derived from the ATP synthase unit of the extinct parrot *Psephotellus pulcherrimus*, and arctodutin-1, a MEP that is part of the enzyme NADH-ubiquinone oxidoreductase chain 5 from the extinct bear *Arctodus simus*. Both enzymes would have played a crucial role in the metabolism of these extinct organisms. These results show that the two MEPs, psephotellin-1 and arctodutin-1, are more effective permeabilizers than most known AMPs and EPs derived from human proteins reported previously^13^. Outer membrane permeabilization is the most common mechanism of action described for AMPs and a key mechanistic driver for EPs derived from modern human proteins, such as natriuretic peptide, SCUB1-SKE25, and SCUB3-MLP22^13^. However, the permeabilization effect exhibited by AEPs and MEPs from extinct organisms was not as potent as that shown by EPs from modern human proteins.

### Synergistic interactions of encrypted peptides

To investigate whether EPs from the same extinct organism can synergize and thus potentiate each other’s activity against pathogens, we performed checkerboard assays^13^ at peptide concentrations ranging from twice the MIC to concentrations up to 64-times lower in the same conditions as used for the antimicrobial assays. First, we selected the peptides according to their MIC values (**Fig. 4a and Fig. S47**) for two pathogenic strains, *A. baumannii* ATCC 19606 and *P. aeruginosa* PAO1. The former is an opportunistic nosocomial pathogen with increasing antibiotic resistance that has resulted in significant mortality worldwide^26^. The latter is an intrinsically resistant bacterium associated with infections of the urinary tract, gastrointestinal tissue, skin, and soft tissues and a cause of bacterial pneumonia, as well as a common opportunistic pathogen in cystic fibrosis patients^27^. Most of the combinations tested resulted in synergistic or additive interactions, calculated by using the fractional inhibitory concentration index^28^ (FICI, **Fig. 5e**). The MICs of combined EPs (**Fig. S47**) were mostly 2- to 3-fold lower than those of the individual peptides, but in some cases, e.g., equusin-1 and equusin-3, the MICs decreased 50 times (from 4 μmol L^-1^ to 78 nmol L^-1^), reaching sub-micromolar concentrations that are comparable to the MICs of some of the most potent antibiotics^29^.

Several pairs of EPs demonstrated particularly strong synergistic interactions, with FICI values as low as 0.38 for *A. baumannii*. These pairs included hesperelin-1 (AEP) and hesperelin-3 (MEP) from *Hesperelaea palmeri*, mammuthusin-1 (MEP) and mammuthusin-3 (MEP) from *Mammuthus primigenius*, equusin-2 (AEP) and equusin-4 (AEP) from *Equus quagga boehmi*, as well as hydrodamin-1 (AEP) and hydrodamin-3 (MEP) from *Hydrodamalis gigas* (**Fig. 5e**).

### Cytotoxicity assays

All 41 active AEPs and MEPs identified in the antimicrobial assays (**Fig. 4a**) were tested for cytotoxic activity against human embryonic kidney (HEK293T) cells (**Table S17**), an extensively characterized cell line. This assay is widely used to assess the toxicity of antimicrobials because the results are highly reproducible^30–32^. Of the peptides tested, 39 displayed no significant cytotoxicity at the concentration range tested (8-128 μmol L^-1^). Cytotoxicity was detected for the AEP lophisin-1 from the ancient crested rat (*Lophiomys imhausi maremortum*) and for the MEP xenothrixin-1 from the extinct Jamaican monkey (*Xenothrix mcgregori*). The peptide dose that led to 50% cytotoxicity (CC_50_), estimated by non-linear regression, was 68.02 and 70.77 μmol L^-1^, for lophisin-1 and xenothrixin-1, respectively. Despite their slight toxicity, the concentration of these two peptides needed to exert antimicrobial activity was, respectively, 8- and 8.84-times lower than their CC_50_ values, underscoring their value as antimicrobials. The CC_50_ values for the other 39 peptides were higher than the maximum concentration analyzed, reinforcing the overall excellent safety profiles of this class of peptides.

### Resistance to proteolytic degradation assays

The AEPs hydrodamin-1, elephasin-2, and mylodonin-2 and the MEPs mammuthusin-2 and megalocerin-1 were selected for further stability and animal studies because of their potent antimicrobial activity (**Fig. 4a**) and good safety (**Table S17**) profiles. To assess their stability in the presence of human proteases, we exposed the EPs to human serum and aliquots were collected and analyzed for 6 h at 37 °C. The AEP elephasin-2 and the MEP mammuthusin-2, from organisms that belong to the same taxonomic order, *Proboscidea*, were the peptides with higher resistance to proteolytic degradation with ∼40% peptide remaining after 6 h of exposure (**Fig. S48**). Mammuthusin-2 presented slower degradation kinetics, whereas elephasin-2 was present at ∼40% within the first 30 min of the experiment. All the other AEPs and MEP analyzed quickly degraded in the first 30 min to 1 h of the experiment (**Fig. S48**).

### Anti-infective efficacy of encrypted peptides in animal models

To assess whether the active AEPs and MEPs had anti-infective efficacy *in vivo*, we used preclinical mouse models of skin abscess^32^ and intramuscular thigh infection^13,33^. Seven EPs were tested with a single dose at their MIC concentration after the infection was established. We selected five EPs having a wide range of MIC values (1-64 μmol L^-1^) when tested *in vitro* against *A. baumannii*: the MEP elephasin-2 (MIC = 1 μmol L^-1^) from *Elephas antiquus*, the AEP hydrodamin-1 (MIC = 4 μmol L^-1^) from *Hydrodamalis gigas*, the MEP megalocerin-1 (MIC = 8 μmol L^-1^) from *Megaloceros sp*., the MEP mammuthusin-2 (MIC = 32 μmol L^-1^) from *Mammuthus primigenius*, and the AEP mylodonin-2 (MIC = 64 μmol L^-1^) from *Mylodon darwinii*, as well as two EPs that we had found to have activity against *A. baumannii*: the AEP hesperelin-1 (MIC = 8 μmol L^-1^) from *Hesperelaea palmeri*, and the MEP equusin-1 (MIC = 16 μmol L^-1^) from *Equus quagga boehmi*.

In the skin abscess infection model, mice were infected with bacterial loads of 10^6^ cells of the pathogen *A. baumannii* (**Fig. 6a**). Each EP was administered as a single dose over the infected area. After two days, bacterial counts showed that all EPs tested, except hydrodamin-1, markedly reduced the bacterial load by 2-3 orders of magnitude. These results highlight the potential anti-infective activity of these EPs (**Fig. 6b**) for *A. baumannii* infections. After four days, we observed that the EPs had either cleared the infection or reduced it by 3-5 orders of magnitude. The peptide hydrodamin-1, which was not active during the first two days post-infection, demonstrated activity by day 4 (**Fig. 6b**). The results obtained for the more active EPs tested (elephasin-2 and mylodonin-2) indicated antibacterial activity that was comparable to that of the widely used antibiotic polymyxin B, which was used as an antimicrobial control (**Fig. 6b**). Changes in weight, a surrogate measure of toxicity, were monitored from the time of the bacterial injection. No variations in weight, damage to the skin tissue, or other harmful consequences induced by the EPs were seen in the mice throughout our experiments (**Fig. S49**).

**Figure 6.**
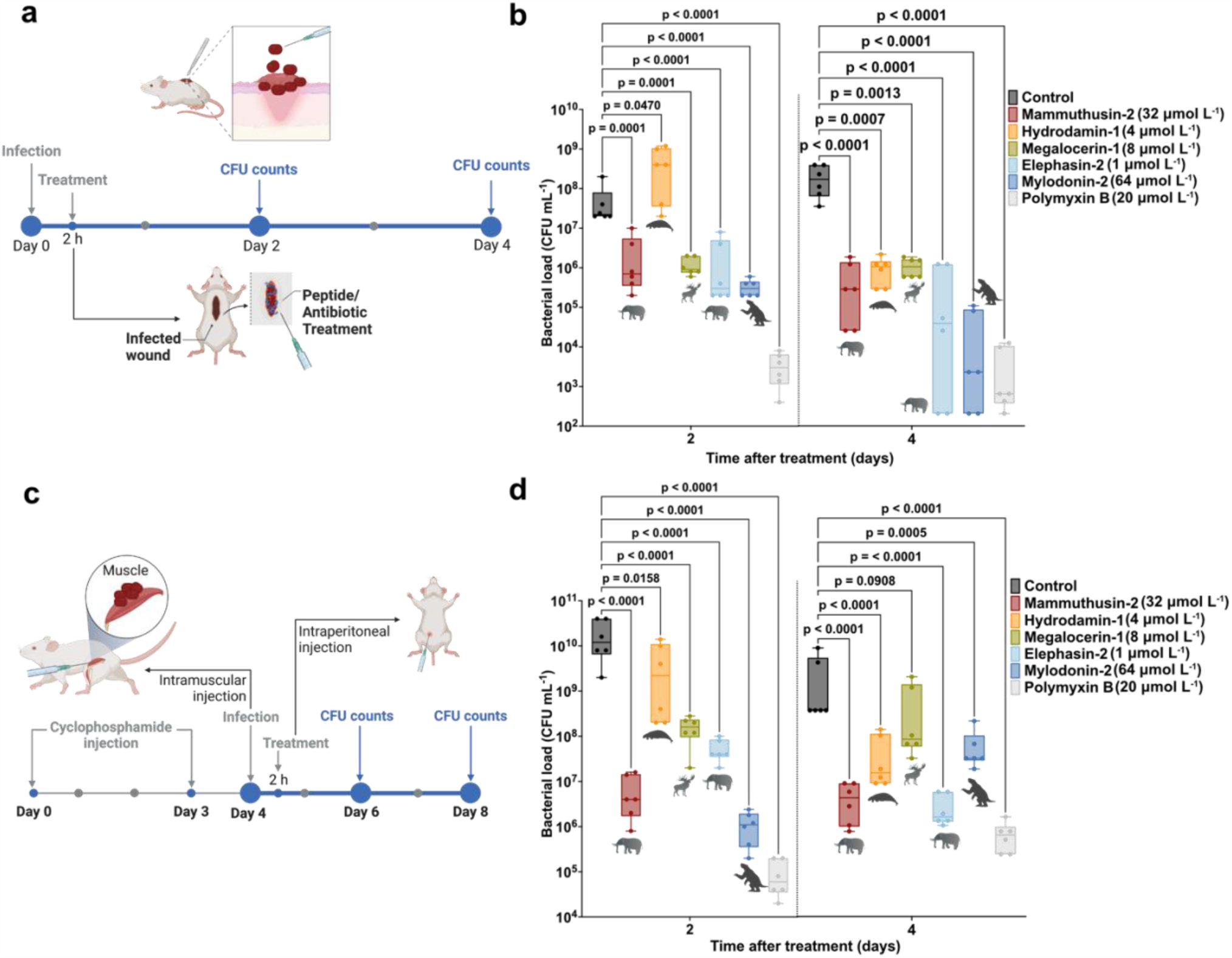
Anti-infective activity of de-extinct peptides in animal models. **(a)** Schematic of the skin abscess mouse model used to assess selected encrypted peptides from extinct organisms (n = 6) for activity against *A. baumannii* ATCC 19606. **(b)** The encrypted peptides mammuthusin-2 (*Mammuthus primigenius*), hydrodamin-1 (*Hydrodamalis gigas*), megalocerin-1 (*Megalocerus sp*.), elephasin-2 (*Elephas antiquus*), and mylodonin-2 (*Mylodon darwinii*), administered at their MIC in a single dose, inhibited the proliferation of the infection for up to four days after treatment compared to the untreated control group. Elephasin-2 and mylodonin-2 cleared the infection in some of the mice, with activity comparable to that of the antibiotic used as control, polymyxin B. **(c)** Schematic of the neutropenic thigh infection mouse model in which encrypted peptides from extinct organisms were injected intraperitoneally. Anti-infective activity against *A. baumannii* ATCC 19606 was assessed 2 and 4 days after systemic administration (n = 6). **(d)** Two days after intraperitoneal injection, mylodonin-2 at its MIC reduced infections caused by *A. baumannii* ATCC19606 as much as polymyxin B, compared to the untreated control group. Four days post-treatment, mammuthusin-2 and elephasin-2 showed the same level of activity as polymyxin B. Statistical significance in **b** and **d** was determined using one-way ANOVA followed by Dunnett’s test; p values are shown in the graph. The solid line inside each box represents the mean value obtained for each group. Figure panels **a** and **c** were created with BioRender.com.

Using a murine deep thigh infection model, we assessed the efficacy of the EPs elephasin-2, hydrodamin-1, megalocerin-1, mammuthusin-2, and mylodonin-2, which were administered after the establishment of the intramuscular thigh infection (**Fig. 6c**). The protocol used is an established preclinical model particularly suited to assess the translatability of potential antibiotics. Briefly, mice were rendered neutropenic by cyclophosphamide treatment before intramuscular injection of 10^6^ *A. baumannii* cells (**Fig. 6c**). Next, a single dose of each peptide at its MIC was injected intraperitoneally. Two- and four-days post-treatment, all peptides tested, except for hydrodamin-2, had reduced the bacterial load by 2-4 orders of magnitude compared to the untreated control group (**Fig. 6d**). Two days post-treatment, peptide mylodonin-2 from *Mylodon darwinii* presented the most potent activity, which was comparable to that of the positive control antibiotic, polymyxin B (4-5 orders of magnitude reduction in bacterial counts). Four days post-treatment, peptides elephasin-2 from *Elephas antiquus* and mammuthusin-2 from *Mammuthus primigenius* had decreased the bacterial loads in the mice by 3-4 orders of magnitude, resulting in levels as low as those in the mice treated with polymyxin B (**Fig. 6d**). None of the peptides tested were harmful to the mice based on weight monitoring during the experimental period (**Fig. S50**).

These substantial *in vivo* results with two preclinical mouse models demonstrate that two AEPs (elephasin-2 and mylodonin-2) and one MEP (mammuthusin-2) displayed anti-infective efficacy comparable to that of a widely used antibiotic under physiologically relevant conditions, underscoring the potential of molecular de-extinction as a sound approach for antibiotic discovery. Additionally, peptide cocktails led to synergistic interactions *in vitro*. For example, combining the MEPs mammuthusins-1 and 5, both derived from the woolly mammoth (*Mammuthus primigenitus*), increased antimicrobial activity against *A. baumannii* cells *in vitro* by 2- and 3-fold compared to each peptide alone.

This systematic analysis of all extinct organisms as a source of previously unrecognized antimicrobials demonstrates the concept of molecular de-extinction. Additionally, our deep learning model outperforms previous work in this emerging area^2^ and constitutes an important platform for discovering new antibiotics by mining proteomes. Paleoproteome mining enables the exploration of new sequence space, expanding our vision of molecular diversity and potentially unlocking new biology. We hypothesize that EPs play a role in immunity throughout evolution and future work will be needed to test this notion. Finally, our approach yielded preclinical candidates with activity comparable to the standard of care (e.g., polymyxin B), highlighting its broad potential applications in biotechnology and medicine. In sum, we have mined the proteomes of all available extinct organisms and have identified antibiotics effective against some of the bacterial pathogens most dangerous to our society.

### Limitations

We have leveraged DL to establish molecular de-extinction as a framework for antibiotic discovery. As a nascent area of research, however, this study has some limitations. For example, our DL model is purely sequence-based and does not contain structural information. Structural and three-dimensional descriptors may be incorporated in the future to increase the accuracy of the model to predict antimicrobial activity. Our model is also limited by the number of sequences present in our in-house dataset. Future work will focus on expanding this dataset to characterize larger spaces of peptide sequence and antimicrobial activity. Another limitation of our platform is the dearth of information available on extinct proteins, as most available proteomes so far have yielded only a few dozen proteins with reliable sequencing information. Our AEP classification was similarly limited by currently available extant proteomes, which may change in the future as new proteomes are revealed. In this work, we present a proof-of-concept demonstration of the de-extinction of antimicrobial molecules from extinct organisms by combining deep learning with wet-lab validation both *in vitro* and in animals. Our approach of mining proteomes from extinct organisms unveils a previously untapped source of potential antibiotics. Furthermore, molecular de-extinction holds the potential to provide a source for other medicinal discoveries in the future.

## Acknowledgments

Cesar de la Fuente-Nunez holds a Presidential Professorship at the University of Pennsylvania and acknowledges funding from the Procter & Gamble Company, United Therapeutics, a BBRF Young Investigator Grant, the Nemirovsky Prize, Penn Health-Tech Accelerator Award, and the Dean’s Innovation Fund from the Perelman School of Medicine at the University of Pennsylvania. Research reported in this publication was supported by the Langer Prize (AIChE Foundation), the National Institute of General Medical Sciences of the National Institutes of Health under award number R35GM138201, and the Defense Threat Reduction Agency (DTRA; HDTRA11810041, HDTRA1-21-1-0014, and HDTRA1-23-1-0001). We thank Dr. Mark Goulian for kindly donating the following strains: *Escherichia coli* AIC221 [*Escherichia coli* MG1655 phnE_2::FRT (control strain for AIC 222)] and *Escherichia coli* AIC222 [*Escherichia coli* MG1655 pmrA53 phnE_2::FRT (polymyxin resistant)]. We thank Dr. Andy Goodman for kindly donating wild type *Bacteroides thetaiotaomicron* (Background: VPI 5482), which was only used as part of the training set for the computational work presented here. The following organisms, which were only used as part of the training set for the computational work presented here, were obtained through BEI Resources, NIAID, NIH: *Salmonella enterica* subsp. *enterica*, Strain ATCC®9150™, NR-515, *Salmonella enterica* subsp. *enterica*, NR-170, *Salmonella enterica* subsp. *enterica*, Strain LT2, NR-174, *Listeria monocytogenes*, Strain Li 20, NR-106. We thank Dr. Karen Pepper for editing the manuscript and de la Fuente Lab members for insightful discussions. We thank Dr. Myra Laird for revising the content of Figure 1. Figures created with BioRender.com are attributed as such. Molecules were rendered using the PyMOL Molecular Graphics System, Version 2.1 Schrödinger, LLC.

## Author contributions

Conceptualization: FW, MDTT, CFN

Methodology: FW, MDTT

Prepared phylogenetic distance matrix: JP

Investigation: FW, MDTT

Visualization: FW, MDTT

Funding acquisition: CFN

Supervision: CFN

Software: FW

Formal analysis: FW, MDTT

Writing – original draft: FW, MDTT, CFN

Writing – review & editing: FW, MDTT, CFN

Writing – review & editing final version of the paper: FW, MDTT, JP, CFN

## Methods

### Datasets

#### Proteomes of extinct organisms

We collected extinct organisms from the NCBI taxonomy browser (https://www.ncbi.nlm.nih.gov/Taxonomy/taxonomyhome.html/index.cgi?chapter=extinct, <u>access time: December 2021</u>). For each species, we checked the corresponding Entrez records and downloaded the available protein sequences. In total, we retrieved 208 extinct species and a total of 12,860 protein sequences (5,190 non-redundant protein sequences) from them.

#### Modern human proteome

To construct the human proteome, we downloaded 20,388 reviewed *Homo sapiens* proteins (20,307 unique ones) from UniProt (https://www.uniprot.org/).

#### In-house peptide dataset

We utilized our high-quality in-house peptide dataset to train and evaluate APEX. In total, the dataset contained 14,738 antimicrobial activity measurements obtained by determining the minimum inhibitory concentration (MIC) of 988 peptides and 34 bacterial strains, including the following, which were used to train our model: *Escherichia coli* ATCC 11775, *Pseudomonas aeruginosa* PAO1, *Pseudomonas aeruginosa* PA14, *Staphylococcus aureus* ATCC 12600, *Escherichia coli* AIC221, *Escherichia coli* AIC222, *Klebsiella pneumoniae* ATCC 13883, *Acinetobacter baumannii* ATCC 19606, *Akkermansia muciniphila* ATCC BAA-835, *Bacteroides fragilis* ATCC 25285, *Bacteroides vulgatus* (*Phocaeicola vulgatus*) ATCC 8482, *Collinsella aerofaciens* ATCC 25986, *Clostridium scindens* ATCC 35704, *Bacteroides thetaiotaomicron* ATCC 29148, *Bacteroides thetaiotaomicron* Δ*tdk* Δ*lpxF* (Background: VPI 5482)^34^, *Bacteroides uniformis* ATCC 8492, *Bacteroides eggerthi* ATCC 27754, *Clostridium spiroforme* ATCC 29900, *Parabacteroides distasonis* ATCC 8503, *Prevotella copri* DSMZ 18205, *Bacteroides ovatus* ATCC 8483, *Eubacterium rectale* ATCC 33656, *Clostridium symbiosum* ATCC 14940, *Ruminococcus obeum* ATCC 29174, *Ruminococcus torques* ATCC 27756, methicillin-resistant *Staphylococcus aureus* ATCC BAA-1556, *vancomycin-resistant Enterococcus faecalis* ATCC 700802, *vancomycin-resistant Enterococcus faecium* ATCC 700221, *Escherichia coli* Nissle 1917, *Salmonella enterica* ATCC 9150 (BEIRES NR-515), *Salmonella enterica* (BEIRES NR-170), *Salmonella enterica* ATCC 9150 (BEIRES NR-174), *Listeria monocytogenes* ATCC 19111 (BEIRES NR-106).

Inactive data points, *i*.*e*., MIC values higher than 128 μmol L^-1^, were labeled as 140 μmol L^-1^. All antimicrobial activities were transformed by 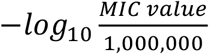 and were treated as labels to be predicted in the machine learning (ML) setting. To perform hyperparameter tuning and prediction performance evaluation, our in-house dataset was randomly split into a cross validation (CV) set and an independent set, which consisted of 790 and 198 peptides (*i*.*e*., an 80%:20% split), respectively. Here, the CV set was used to determine the optimal hyperparameters for ML models. ML models trained with determined hyperparameters on the CV set were evaluated on the independent set to measure their generalizability.

### Publicly available AMP sequences

We augmented the peptide training data by incorporating publicly available AMPs and non-AMPs into our DL model training. Public AMP data was retrieved from the DBAASP^15^. Peptide sequences that consisted of only the 20 canonical or unknown amino acid residues were selected. The unknown amino acids were denoted by X, which corresponds to any possible canonical amino acid residues that usually occurred for the proteins having isoforms proteins whose composition was undetermined because there were issues with metagenomic studies. Any peptide sequences that overlapped with our in-house peptide database were removed. As a peptide may have multiple MIC values for different bacterial species, we used the median MIC value to binarize the data. By using a stringent cutoff (i.e., AMPs with MIC ≤30 μmol L^-1^), we created a balanced binary classification dataset (5,093 AMPs and 5,500 non-AMPs) for data augmentation and model training. To compare physicochemical properties, 14,114 peptides consisting of 20 canonical amino acids and having sequence length ≥4 amino acid residues were retrieved. We labeled this group of peptides as the DBAASP dataset.

### Physicochemical properties of peptides

To analyze the physicochemical properties of the all peptide datasets (DBAASP, EPs generated by the scoring function^13^, and EPs generated by APEX), we used the DBAASP server to calculate the following twelve physicochemical properties that are usually considered in the design and study of peptide antibiotics^15^: normalized hydrophobic moment, normalized hydrophobicity, net charge, isoelectric point, penetration depth, tilt angle, disordered conformation propensity, linear moment, propensity to aggregation *in vitro*, angle subtended by the hydrophobic residues, amphiphilicity index, and propensity to PPII coil. We used the Eisenberg and Weiss scale (the consensus scale) as the hydrophobicity scale^35^.

### Secreted protein labeling

As proteins from extinct organisms are not as well annotated as those from extant organisms, we resorted to Orthologous Matrix (OMA)^36^ and DeepGOWeb^37^ to predict gene ontology (GO) terms from protein sequences. Given a protein sequence, if any of its GO terms predicted by OMA or DeepGOWeb corresponds to an extracellular region (GO:0005576) or a child of extracellular region in a GO-directed acyclic graph, then we considered this sequence to be secreted. Among the proteins from the extinct organisms we collected, 157 sequences are labeled as secreted.

### Peptide sequence encoding

We treated a peptide as a sequence of amino acids. We further added two special characters as start (*i*.*e*., ‘1’) and terminal symbols (*i*.*e*., ‘2’) to the beginning and the end of this sequence, respectively. For each amino acid in the sequence, we used the AAindex^38^, which is a 566-dimensional vector storing various physicochemical and biochemical properties of each amino acid to represent it. Non-amino acid symbols and unknown amino acids were represented by 566-dimensional zero vectors. In this work, we only considered peptides shorter than 50 residues to ensure that they could be synthesized by solid phase peptide synthesis. We created a fixed size input by zero-padding each sequence to maximum length, so that each peptide sequence can be represented by a matrix ***x*** ∈ ℝ^52×566^ (52 = maximum peptide length + two special characters).

### Bacterial distance

Taxonomy tree (bac120.tree) was downloaded from the Genome Taxonomy Database (GTDB)^39^. The phylogenetic distance matrix ***D*** ∈ ℝ^*g*×*g*^, which stores distances between bacterial species, was calculated via the python package DendroPy^40^. Here, *g* denotes the number of bacterial species. We converted the distance matrix ***D*** to a bacterial similarity matrix ***P*** ∈ ℝ^*g*×*g*^ using the following function:

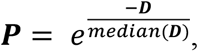

where *median*(***D***) stands for the median value of matrix ***D***. We further binarized the similarity matrix ***P*** using the K nearest neighbor algorithm, in which K was set to a heuristic number *Ceiling* 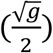. Here, *Ceiling*(•) stands for the ceiling function.

### Construction of APEX

#### The encoder architecture

The encoder started from a recurrent neural network (RNN) to process peptide sequence input ***x*** and extract its hidden features, ***h***_***rnn***_ ∈ ℝ^52×*n*^:

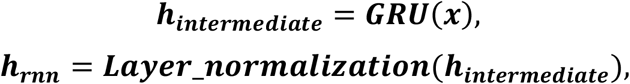

where *n* and ***GRU***(•) denotes the hidden feature dimension and gated recurrent unit^41^, respectively. In addition, we added a layer normalization^42^ to stabilize the model training. On top of the RNN, we designed a two-layer attention neural network to better model feature (i.e., amino acids) interactions globally and compress the hidden features to a lower-dimensional representation, respectively. Specifically, the first attention layer has the following form:

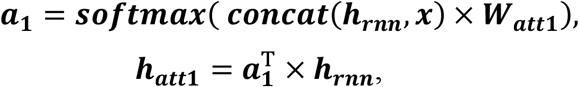

where ***concat***(***h***_***rnn***_, ***x***) ∈ ℝ^52×(*n*+566)^ stands for concatenation operation along feature dimension and can be considered as a residual connection^43^, ***W***_***att*1**_ ∈ ℝ^(*n*+566)×52^ is the learnable weights in this attention layer, ***softmax***(•) stands for the softmax function, ***a***_**1**_ ∈ ℝ^52×52^ denotes the attention weights, and ***h***_***att*1**_ ∈ ℝ^52×*n*^ is the output of this attention layer. For the second attention layer, it can be written as:

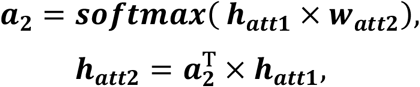

where ***w***_***att*2**_ ∈ ℝ^*n*×1^ is the learnable weights in this attention layer, ***a***_**2**_ ∈ ℝ^52×1^ denotes the attention weights, and ***h***_***att*2**_ ∈ ℝ^1×*n*^ is the output of the second attention layer. In addition, we used a learnable linear transformation with weight matrix ***W***_***fc***_ ∈ ℝ^*n*×*m*^ and bias term ***b***_***fc***_ ∈ ℝ^1×*m*^ to create the final hidden representation ***h*** ∈ ℝ^1×*m*^ for a peptide:

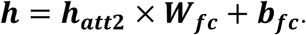

#### FCNNs on the prediction of sequences with antimicrobial activity

The hidden representation ***h*** of a peptide generated by the encoder above could be fed into two separate FCNNs that predict species-specific antimicrobial activity or a binary AMP/non-AMP label, respectively. For convenience of hyperparameter tuning, both FCNNs were implemented as a 4-layer pyramid-like architecture:

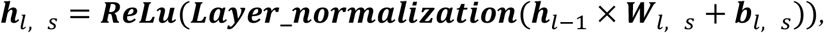

where *s* ∈ {*in* − *house, public*} denotes the training dataset for the FCNN, *l* ∈ {1,2,3,4} denotes the layer index (note that ***h***_0_ *=* ***h***), ***W***_*l, s*_ and ***b***_*l, s*_ are weight matrix and bias term of *l*th layer, respectively. At *l*th layer, a linear transformation was first performed and followed by a layer normalization, a nonlinear transformation using rectified linear unit^44^ (ReLU). In addition, if the *l*th layer is not an output layer, a dropout layer^45^ that randomly set the input value to zero with probability *p* was added to its output side (we empirically set *p =* 0.1 ). The output dimensions of hidden layers (i.e., ***h***_1, *s*_, ***h***_2, *s*_, and ***h***_3,*s*_) were set as 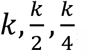, respectively. The FCNN that was trained on our in-house data adopted a multitask learning strategy to predict species-specific antimicrobial activity. Suppose there are *g* bacterial species (*i*.*e*., 34 in our context), the corresponding output ***h***_4, *in*−*house*_ is a *g*-dimensional vector, in which each element is a predicted antimicrobial activity against a certain bacterial strain. The FCNN that was trained on public AMP data only outputted a scaler value ∈[0,1] indicating the probability of the input peptide to be antimicrobial.

#### Loss function

The loss function for training the FCNN that performed binary classification was binary cross-entropy *l*_*BCE*_. For the other FCNN, the loss function for predicting species-specific antimicrobial activity was the mean squared error:

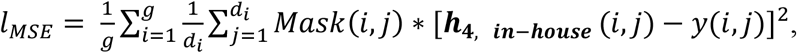

where *d*_*i*_ denotes the number of training data points for *i*th bacterial strain, ***h***_**4, *in***−***house***_ (*i, j*), *y*(*i, j*) and *Mask*(*i, j*) are the predicted antimicrobial activity, the experimentally validated antimicrobial activity, and the binary mask (1 for having antimicrobial activity, and 0 for not tested) between *i*th bacterial strain and *j*th peptide. In addition to these two loss functions on AMP prediction, we further imposed a constraint loss on the weights of output layer in the species-specific AMP prediction FCNN. Given a bacterial distance matrix ***P*** ∈ ℝ^*g*×*g*^, and the weights ***W***_4, *in*−*house*_ that we want to regularize, the constraint loss can be written as:

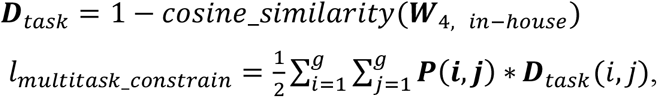

where *cosine*_*similarity*(•) calculates the pairwise cosine similarity between two rows of the given input matrix, matrix ***D***_*task*_ ∈ ℝ^*g*×*g*^ stores the pairwise cosine distance between two learnable weights of two predictors. Intuitively, if two tasks are similar, their predictors should also be similar (*i*.*e*., learnable weights have shorter distances). Adding *l*_*multitask*_*constrain*_ to the loss function encourages similar bacterial strains to have similar predictors and outputs. Taken together, the final loss function has the following form:

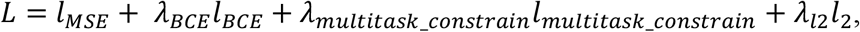

where *l*_2_ is the L2 regularization for constraining DL model complexity, *λ*_*BCE*_, λ_*multitask*_*constrain*_, and *λ*_*l*2_ are the weight parameters that balances different types of losses. To train the DL models, we used mini-batch training with an Adam optimizer^46^. Specifically, at each iteration, we selected *B* peptides from the in-house dataset and the same number of peptides from public AMP data we curated to perform feed-forwarding pass and back propagation. The training terminated when the procedure iterated the whole in-house dataset 5,000 times. The learning rate of the Adam optimizer was empirically set to 0.0001 and was scheduled to decay ten times every thousand training epochs. Batch size *B* was empirically set to 128.

### Baseline methods

We compared the prediction performance of APEX to that of several baseline ML models, including elastic net, linear support vector regression, extra-trees regressor, random forest, and gradient boosting decision tree. For the baseline models, we represented each peptide sequence by the following features: (i) k-mer (*i*.*e*., frequency of k-residue substrings, where k=1, 2 and 3) and (ii) ten peptide properties calculated by modlAMP^47^, including sequence length, molecular weight, sequence charge, charge density, isoelectric point, instability index, aromaticity, aliphatic index, Boman index, and hydrophobic ratio. Note that for some of the bacterial strains, the trained Elastic Net outputted a constant prediction regardless of the peptide inputs. In this case, the Pearson and Spearman correlations could not be calculated, and we used 0 as pseudo correlation.

### Hyperparameter tuning, model evaluation and ensemble learning

We conducted a five-fold cross validation on the CV set to select the hyperparameters of DL and baseline models. Specifically, the five-fold cross validation split the whole dataset evenly into five groups. At each time, one group was selected as the test dataset, while the rest was used for ML model training. We used averaged R-squared, Pearson correlation coefficient, and Spearman’s rank correlation coefficient under five-fold cross validation to evaluate the prediction performance on the test set. We used grid search to find the best hyperparameters (see **Tables S1-S4** for hyperparameter range we searched). Hyperparameters were ranked by the averaged R-squared under cross validation. For baseline methods, we determined the best hyperparameters to be the ones resulting in the highest R-squared and trained the ML models with the selected hyperparameters. The trained models were then evaluated on the independent dataset. For APEX, we adopted an ensemble learning strategy. Specifically, we averaged the prediction results from the top eight APEX models. After plotting the prediction performances versus the number of DL models involved in the ensemble learning (**Figs. S8-S10**), we observed that the elbow region (i.e., the area where the curve becomes smaller) was around 5-9 APEX models. After this step, improvement on prediction performance gradually became negligible. This observation led to the conclusion that we should average prediction results from no more than nine APEX models. We decided to select eight APEX models for ensemble learning, given our computational resources (*i*.*e*., eight GPUs were available). To counter the potential stochastic behavior during mini-batch training and in order to make prediction results more robust, we trained five copies of an APEX model with the same hyperparameters under different random seeds. In total, we trained 40 APEX models (*i*.*e*., eight different hyperparameters × five different random seeds) and used the averaged predictions on the independent dataset for prediction performance evaluation. After the performance evaluation, we retrained the 40 APEX models on the entire in-house dataset and used the averaged antimicrobial activity prediction values from the trained models to discover encrypted peptide sequences from extinct organisms.

### Selectivity score

Since APEX was designed to predict species-specific antimicrobial activity, we defined the following two selectivity scores that quantify peptides’ ability to specifically target Gram-positive or Gram-negative bacteria:

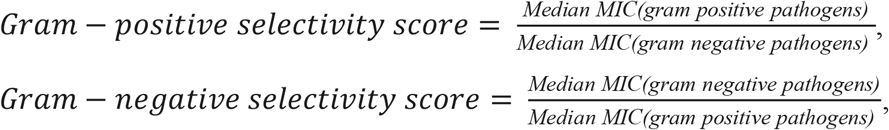

where *Median MIC*(•) calculates the median value from a given input list. The input list consisted of the Gram-positive pathogens *S. aureus* ATCC 12600, methicillin-resistant *S. aureus* ATCC BAA-1556, vancomycin-resistant *E. faecalis* ATCC 700802, and vancomycin-resistant *E. faecium* ATCC 700221 and the Gram-negative pathogens *P. aeruginosa* PAO1, *P. aeruginosa* PA14, *E. coli* ATCC11775, *E. coli* AIC221, *E. coli* AIC222, *K. pneumoniae* ATCC 13883, and *A. baumannii* ATCC 19606. A selectivity score < 1.0 means that the median MIC of the target bacteria (numerator term) is smaller than that of the off-target bacteria (denominator term), yielding a selective peptide sequence. Thus, the closer to zero the better is the selective activity towards the specific bacterial target.

### Sequence similarity score

Given two peptide sequences *i* and *j*, we used the Smith-Waterman algorithm^48^ to calculate their sequence alignment score *SW*(*i, j*). The sequence similarity score between these two peptides was defined as the normalized alignment score: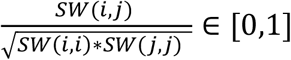. A higher score reflects higher sequence similarity between two peptides than a lower score.

### Encrypted peptide screening and selection from extinct proteomes

Given the proteome of extinct organisms, we considered as encrypted peptide (EP) sequences substrings ranging from 8 to 50 amino acid residues. In total, paleoproteome mining yielded 10,311,899 unique EPs from extinct proteomes of which 771,431 peptide sequences came from secreted proteins. EPs from secreted proteins would have had a higher likelihood of encountering bacterial cells, which are mostly found outside host cells, than EPs from non-secreted proteins Nevertheless, most EPs came from non-secreted proteins and consequently represented a more abundant peptide source. We hypothesized that these non-secreted proteins might also contain antibiotic-like substrings. Therefore, in this work, we selected, synthesized, and validated EPs originating from both secreted and non-secreted proteins.

Since we used 40 APEX models for the activity prediction (*i*.*e*., ensemble APEX v2 or APEX in the main text), we averaged the prediction results from these 40 models to provide a final predictive output for each peptide. Based on these APEX predictions, we used multiple criteria to select EPs from our predictions for downstream validation. First, we mainly focused on EPs that could target a subset of the eleven pathogens that were listed in the ***Selectivity score*** section (i.e., *E. coli* ATCC 11775, *P. aeruginosa* PAO1, *P. aeruginosa* PA14, *S. aureus* ATCC 12600, *E. coli* AIC221, *E. coli* AIC222, *K. pneumoniae* ATCC 13883, *A. baumannii* ATCC 19606, methicillin-resistant *S. aureus* ATCC BAA-1556, vancomycin-resistant *E. faecalis* ATCC 700802, and vancomycin-resistant *E. faecium* ATCC 700221). Given the entire encrypted peptide list, we first generated two encrypted peptide lists; one with sequences predicted to selectively target Gram-positive and another with sequences predicted to target Gram-negative pathogens. To this end, the peptides in these lists were ranked increasingly by the *Gram* − *positive selectivity score* and the *Gram* − *ne*g*ative selectivity sc*o*re*, respectively. We then generated another peptide list by ranking the peptides based on their median MIC predictions for the eleven pathogens of interest, indicating the level of broad-spectrum antimicrobial activity of the peptides.

As a result, three lists were generated for EPs derived from non-secreted proteins and another three lists were generated for EPs derived from secreted proteins, corresponding in each case to: (1) EPs predicted to selectively target Gram-positive bacteria; (2) EPs predicted to selectively target Gram-negative bacteria; and (3) EPs predicted to have a broad-spectrum of activity. The six lists were filtered by: (1) including only peptides ranging from 8 to 30 residues, as they are easier to synthesize and as they comprise the range of sequence length in which most peptides with antimicrobial activity are active; (2) excluding sequences that appeared in our previous modern human proteome exploration^13^, (3) excluding peptides that appeared in our in-house dataset, which is composed of natural and synthetic sequences, as well as computationally designed peptides; (4) excluding peptides that showed sequence similarity >0.75 to any peptide from the DBAASP dataset, which ensured that we would not explore peptides derived from modern natural versions of reported AMPs (i.e., we only kept EPs that were not similar to modern AMPs); (5) if two EPs in a list had a sequence similarity >0.75, keeping only the one that ranked higher (i.e., the more active or more selective peptide) in the list, which ensured that we explored the largest sequence space possible; and (6) for each of the lists, selecting a maximum of 1,000 peptides based on the ranking of predicted antimicrobial activity; (7) for broad-spectrum peptide lists, peptides whose median MIC predictions were ≥80 μmol L^-1^ were excluded as they were deemed inactive. For selectivity peptide lists, peptides whose selectivity scores were ≥0.75 were excluded as they were deemed as not selective enough. Finally, we grouped the selected peptides by their source organisms. We selected the top (*i*.*e*., more active or selective) EPs while also taking into account species and sequence diversity (**Data S2**). Twenty-one EPs from secreted extinct proteins were selected for synthesis and subsequent experimental validation: 4 that were predicted to selectively target Gram-positive bacteria, 10 that were predicted to selectively target Gram-negative bacteria, and 7 that were predicted to have broad-spectrum activity. Forty-eight EPs from non-secreted extinct proteins were also selected for downstream experimental validation: 19 that were predicted to selectively target Gram-positive bacteria, 10 that were predicted to selectively target Gram-negative bacteria, and 19 that were predicted to display broad-spectrum activity.

### Classification guidelines to identify archaic encrypted peptides (AEPs)

Since de-extinct sequences have not been previously identified, we developed our own classification guidelines to determine whether a particular peptide sequence qualified as truly de-extinct, *i*.*e*., not present in the available proteomic data we had from living organisms. Since EP sequences from the proteomes of extinct organisms may also be found in modern organisms, we used the following procedure to classify an EP sequence as an AEP: First, we accessed the NCBI taxonomy browser (https://www.ncbi.nlm.nih.gov/Taxonomy/taxonomyhome.html/index.cgi?chapter=extinct, <u>access time: December, 2022</u>) to get the most up-to-date taxonomy IDs of extinct organisms. Next, we created a protein sequence set with all possible organism sources by downloading protein sequences and their corresponding taxonomy IDs from Reviewed (Swiss-Prot), Unreviewed (TrEMBL), and Isoform sequences at UniProt^49^ (https://www.uniprot.org/help/downloads). We excluded from the protein sequence set protein sequences whose taxonomy IDs belonged to extinct organisms. We labeled the resulting set as “extant protein set” as it contained protein sequences from extant organisms. If a given EP sequence was not present in any protein sequence from the modern protein set available, we defined it as an AEP.

### Peptide synthesis

All peptides were synthesized by solid-phase peptide synthesis using the Fmoc strategy and purchased from AAPPTec.

### Bacterial strains and growth conditions used in the experiments

The following Gram-negative bacteria were used in our study: *Acinetobacter baumannii* ATCC 19606, *Escherichia coli* ATCC 11775, *Escherichia coli* AIC221 (*Escherichia coli* MG1655 phnE_2::FRT), *Escherichia coli* AIC222 [*Escherichia coli* MG1655 pmrA53 phnE_2::FRT (colistin-resistant)], *Klebsiella pneumoniae* ATCC 13883, *Pseudomonas aeruginosa* PAO1, and *Pseudomonas aeruginosa* PA14. The following Gram-positive bacteria were also used in our study: *Staphylococcus aureus* ATCC 12600, *Staphylococcus aureus* ATCC BAA-1556 (methicillin-resistant strain), *Enterococcus faecalis* ATCC 700802 (vancomycin-resistant strain), and *Enterococcus faecium* ATCC 700221 (vancomycin-resistant strain). Bacteria were grown from frozen stocks and plated on Luria-Bertani (LB) or *Pseudomonas* Isolation (*Pseudomonas aeruginosa* strains) agar plates and incubated overnight at 37 °C. After the incubation period, a single colony was transferred to 5 mL of LB medium, and cultures were incubated overnight (16 h) at 37 °C. The following day, an inoculum was prepared by diluting the overnight cultures 1:100 in 5 mL of the respective media and incubating them at 37 °C until bacteria reached logarithmic phase (OD_600_ = 0.3-0.5).

### Antibacterial assays

The *in vitro* antimicrobial activity of the peptides was assessed by subjecting them to the broth microdilution assay^32^. Minimum inhibitory concentration (MIC) values of the peptides were determined with an initial inoculum of 2×10^6^ cells mL^-1^ in LB in microtiter 96-well flat bottom transparent plates. Aqueous solutions of the peptides were added to the plate at concentrations ranging from 1 to 64 μmol L^-1^. The lowest concentration of peptide that inhibited 100% of the visible growth of bacteria was established as the MIC value in an experiment of 20 h of exposure at 37 °C. The optical density of the plates was measured at 600 nm using a spectrophotometer. All assays were done as three biological replicates.

### Outer membrane permeabilization assays

The membrane permeability of the peptides was determined by using the 1-(N-phenylamino)naphthalene (NPN) uptake assay^13^. NPN is a hydrophobic fluorescent dye that does not readily permeate the bacterial outer membrane. However, when the membrane integrity is compromised, NPN can enter the cell and bind to the bacterial membrane lipids. This causes the dye to exhibit a strong fluorescence. *A. baumannii* ATCC19606 and *P. aeruginosa* PAO1 were grown (OD_600_ = 0.4), centrifuged (10,000 rpm at 4 ºC for 10 min), washed, and resuspended in buffer (5 mmol L^-1^ HEPES, 5 mmol L^-1^ glucose, pH 7.4). NPN solution (4 μL at the working concentration of 10 mmol L^-1^ after dilution) was added to 100 μL of the bacterial solution in a white 96-well plate. The fluorescence was recorded at λ_ex_ = 350 nm and λ_em_ = 420 nm. Aqueous solutions of the peptides (100 μL final volume at their MIC against the strain of interest) were added to a white 96-well plate, and fluorescence was recorded for 20 min after no further increase in fluorescence was observed. All assays were done as three biological replicates. The relative fluorescence values were calculated for the entire course of the experiment using non-linear fitting and the untreated control (buffer + bacteria + fluorescent dye) as baseline. The following equation was applied to show % difference between the fluorescence of the untreated control (baseline) and the sample:

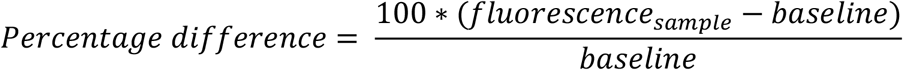

### Cytoplasmic membrane depolarization assays

The depolarization of the bacterial cytoplasmic membrane was determined by fluorescence measurements of the membrane potential-sensitive dye, 3,3’-dipropylthiadicarbocyanine iodide DiSC_3_-5^13^. Briefly, *A. baumannii* ATCC 19606 and *P. aeruginosa* PAO1 were grown at 37 ºC until mid-log phase (OD_600_ = 0.5). The cells were then centrifuged using the same conditions described for the NPN uptake assays, washed twice with washing buffer containing 20 mmol L^-1^ glucose and 5 mmol L^-1^ HEPES (pH 7.2). The cells were diluted 1:10 (OD_600_ = 0.05) in a buffer containing 0.1 mol L^-1^ KCl, 20 mmol L^-1^ glucose and 5 mmol L^-1^ HEPES (pH 7.2). One hundred μL of bacterial solution were then incubated for 15 min with 20 nmol L^-1^ of DiSC_3_-5 until the fluorescence reached a plateau, i.e., the dye was fully internalized into the bacterial membrane. Transmembrane potential changes were monitored by observing the difference in the fluorescence emission intensity of DiSC_3_-5 (λ_ex_ = 622 nm, λ_em_ = 670 nm), after the addition of 100 μL of peptide aqueous solution at its MIC. All assays were performed in three biological replicates. The relative fluorescence values were calculated for the course of the experiment using non-linear fitting and the untreated control (buffer + bacteria + fluorescent dye) served as baseline. The following equation was applied to show % difference between the fluorescence of the untreated control (baseline) and the sample:

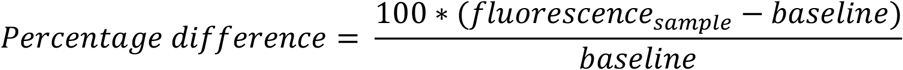

### Synergy between encrypted peptides from extinct organisms

*P. aeruginosa* PAO1 and *A. baumannii* ATCC 19606 were used to assess the synergistic interactions of peptides derived from the same organisms because of their resistance to antimicrobials. The most active de-extinct EPs against *P. aeruginosa* PAO1 and *A. baumannii* ATCC 19606 were orthogonally diluted, using the microdilution technique, to concentrations ranging from 4-times MIC to 0.0625-times MIC in checkerboard assays. Plates were incubated for 20 h at 37 °C. All assays were done in three biological replicates.

### Eukaryotic cell culture conditions

Human embryonic kidney (HEK293T) cells were obtained from the American Type Culture Collection (ATCC; CRL-3216™). The cells were cultured in high-glucose Dulbecco’s modified Eagle’s medium (DMEM) supplemented with 1% penicillin and streptomycin (antibiotics) and 10% fetal bovine serum (FBS) and grown at 37 °C in a humidified atmosphere containing 5% CO_2_.

### Cytotoxicity assays

One day before the experiment, an aliquot of 100 μL of the cells at 50,000 cells mL^-1^ was seeded into each well of the cell-treated 96-well plates used in the experiment (*i*.*e*., 5,000 cells per well). The attached HEK293T cells were then exposed to increasing concentrations of the peptides (8-128 μmol L^-1^) for one day. After the incubation period, we performed the (3-(4,5-dimethylthiazol-2-yl)-2,5-diphenyltetrazolium bromide) tetrazolium reduction assay (MTT assay)^50^. The MTT reagent was dissolved at 0.5 mg mL^-1^ in medium without phenol red and was used to replace cell culture supernatants containing the peptides (100 μL per well), and the samples were incubated for 4 h at 37 °C in a humidified atmosphere containing 5% CO_2_ yielding the insoluble formazan salt. The resulting salts were then resuspended in hydrochloric acid (0.04 mol L^-1^) in anhydrous isopropanol and quantified by spectrophotometric measurements of absorbance at 570 nm. All assays were done as three biological replicates.

### Resistance to proteolytic degradation assays

To assess the resistance of EPs to proteolysis, we incubated them in human serum^51^. The following five lead EPs: mammuthusin-2, hydrodamin-1, megalocerin-1, elephasin-2, and mylodonin-2 (at 3 mg mL^-1^) were exposed to 25% human serum in water for 6 h at 37 °C. One hundred μL aliquots were collected after 0, 0.5, 1, 3, and 6 h, and 10 μL of 100% trifluoroacetic acid (TFA) was added to each sample to induce protein precipitation and incubated for 10 min on ice (at ∼4 °C). Samples were then processed in a Waters Acquity UPLCMS equipped with a photodiode array detector (190-400 nm data collection) and a Waters TQD triple quadrupole MSMS, with 5 μL injections. The column used was a Waters Acquity UPLC HSS C_18_, 1.8 μm (2.1 mm x 50 mm). The mobile phases used were A (100% water with 0.1%, v/v, formic acid) and B (100% acetonitrile with 0.1%, v/v, formic acid), Fisher optima grades. Measurements were made by ionization ESI +/- simultaneous over m/z 100-2,000 Da. The percentage of remaining peptide was calculated by integrating the area under the curve related to the peptide at time point zero. Experiments were performed in three independent replicates.

| Time (min) | A (%) | B (%) | Flow rate (mL min<sup>-1</sup>) |
| --- | --- | --- | --- |
| 0 | 95 | 5 | 0.5 |
| 0.5 | 95 | 5 | 0.5 |
| 2.5 | 5 | 95 | 0.5 (linear gradient) |
| 3 | 5 | 95 | 0.5 |
| 3.25 | 5 | 95 | 0.5 |

### Circular dichroism assays

Circular dichroism assays were performed at the University of Pennsylvania’s Biological Chemistry Resource Center (BCRC) using a J1500 circular dichroism spectropolarimeter (Jasco). The circular dichroism spectra represent an average of three accumulations at 25 °C obtained using a quartz cuvette with an optical path length of 1.0 mm. The experiments covered a wavelength range from 260 to 190 nm at a rate of 50 nm min^-1^ and a bandwidth of 0.5 nm. The concentration of all peptides tested was kept at 50 μmol L^-1^, and the measurements were performed in a mixture of water and trifluoroethanol (TFE) in a 3:2 ratio. A Fourier transform filter was applied to minimize background effects. Secondary structure fraction values were calculated using the single spectra analysis tool on the server BeStSel^52^.

### Skin abscess infection mouse model

*A. baumannii* ATCC 19606 were grown in LB medium to an OD_600_ = 0.5. Cells were washed twice with sterile PBS (pH 7.4, 13,000 rpm for 2 min) and resuspended to a final concentration of 2×10^5^ (*A. baumannii* cells) colony-forming units (CFU) mL^-1^. Six-week-old female CD-1 mice from Charles River (stock number 18679700-022) were anesthetized with isoflurane and their backs were sterilized and shaved. A superficial linear skin abrasion was made with a needle to damage the stratum corneum and upper layer of the epidermis. An aliquot of 20 μL containing the bacterial load resuspended in PBS was inoculated over the scratched area. One hour after the infection, peptides diluted in water at their MIC value were administered to the infected area. Mice were euthanized and the area of scarified skin was excised two- and four-days post-infection, homogenized using a bead beater for 20 min (25 Hz), and 10-fold serially diluted for CFU quantification in MacConkey agar plates. The experiments were performed with 6 mice per group. All experiments were performed blindly, and no animal subjects were excluded from the analysis. The skin abscess infection mouse model was approved by the University Laboratory Animal Resources (ULAR) from the University of Pennsylvania (Protocol 806763). Statistical significance was determined using one-way ANOVA followed by Dunnett’s test in a log_10_-transformed data to mitigate the effect of outliers; p values are presented for each group, with all groups being compared to the untreated control group.

### Thigh infection mouse model

Six-week-old female CD-1 mice from Charles River (stock number 18679700-022) were rendered neutropenic by two doses of cyclophosphamide (150 mg Kg^-1^) applied intraperitoneally with an interval of 72 h. One day after the last dose of cyclophosphamide, the mice were injected intramuscularly in their right thigh with a bacterial load of 10^6^ CFU mL^-1^ of *A. baumannii* ATCC 19606 cells. The bacteria had been grown in LB broth, washed twice with PBS (pH 7.4), and resuspended to the desired concentration. Two hours after bacterial injection, peptides resuspended in water were administered intraperitoneally. Prior to each injection, mice were anesthetized with isoflurane and monitored for respiratory rate and pedal reflexes. Next, we monitored the establishment of the infection and euthanized the mice. The infected area was excised two days and four days post-infection, homogenized using a bead beater for 20 min (25 Hz), and 10-fold serially diluted for CFU quantification in MacConkey agar plates. The experiments were performed with 6 mice per group. All experiments were performed blindly, and no animal subjects were excluded from the analysis. The thigh infection mouse model was approved by the University Laboratory Animal Resources (ULAR) from the University of Pennsylvania (Protocol 807055). Statistical significance was determined using one-way ANOVA followed by Dunnett’s test in a log_10_-transformed data to mitigate the effect of outliers; p values are presented for each group, with all groups being compared to the untreated control group.

### Statistical analysis

Unless otherwise stated, all assays were performed in three independent biological replicates as indicated in each figure legend and Methods sections. The cytotoxic activities values were estimated using non-linear regression based on the range of concentrations screened and were shown as the values that cause lysis of 50% of the cells in the experiment. Two technical replicates were performed in the cytotoxicity assays within each of the three biological replicates.

In the mouse experiments, the statistical significance was determined using one-way ANOVA followed by Dunnett’s test. All the p values are shown for each of the groups, all groups were compared to the untreated control group. The solid line inside each box represents the mean value obtained for each group.

All calculation and statistical analyses of the experimental data were conducted using GraphPad Prism v.10.0.2 and computational data were performed in Python. Statistical significance between different groups was calculated using the tests indicated in each figure legend. No statistical methods were used to predetermine sample size.

### Data availability

- Proteomes from which EPs were obtained are publicly available from the NCBI and UniProt: https://www.ncbi.nlm.nih.gov/Taxonomy/taxonomyhome.html/index.cgi?chapter=extinct#4D616D6D616C73 and https://www.uniprot.org/.
- All data pertaining to the experimental validation of generated peptides are available in the Supplementary Data and upon request.
- Pretrained APEX models are available on GitLab (https://gitlab.com/machine-biology-group-public/apex).
- Any additional information required to reanalyze the data reported in this paper is available from the lead contact upon request.

