## Supplementary Information File for "Molecular de-extinction of antibiotics enabled by deep learning"

**This PDF file includes:**

Abbreviations
Figs. S1 to S50
Tables S1 to S17
Caption for Data S1 to S3

**Other Supplementary Materials for this manuscript include the following:**

Data S1 to S3

**Supplementary section**

**Abbreviations:**

EP: encrypted peptides; Encrypted peptides are defined as peptide sequences embedded in the proteome (*i.e.*, the sequence of proteins);
MEPs: modern EPs, EPs that are present in extant organisms;
AEPs: archaic EPs, EPs that are only present in extinct organisms;
AMPs: antimicrobial peptides from available databases;
DL: deep learning;
APEX: Antibiotic Peptide de-EXtinction model.

**Supplementary Figures:**

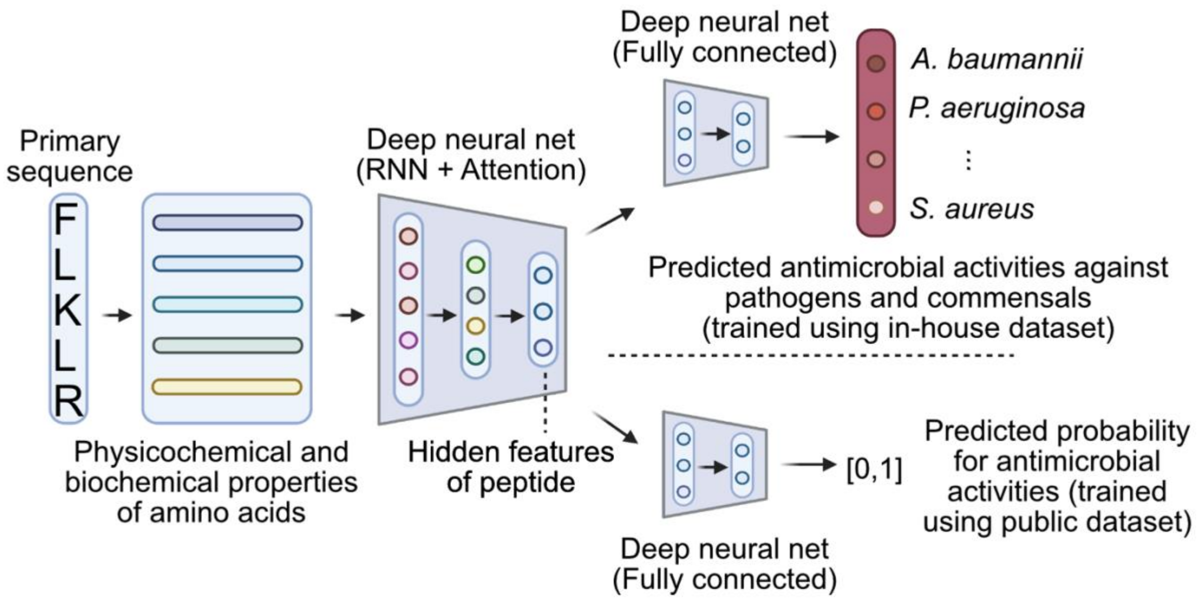


**Fig. S1. Schematic illustration of APEX.** APEX utilized a hybrid of recurrent and attention neural networks to extract peptide sequence information. Extracted hidden features for peptides from in-house or public datasets were processed by two fully connected neural networks to predict species-specific antimicrobial activities (i.e., a multi-output regression task) or general antimicrobial or not (i.e., a binary classification task), respectively. We adopted this multitask framework to accurately predict whether a given peptide sequence was likely to have antimicrobial activity.

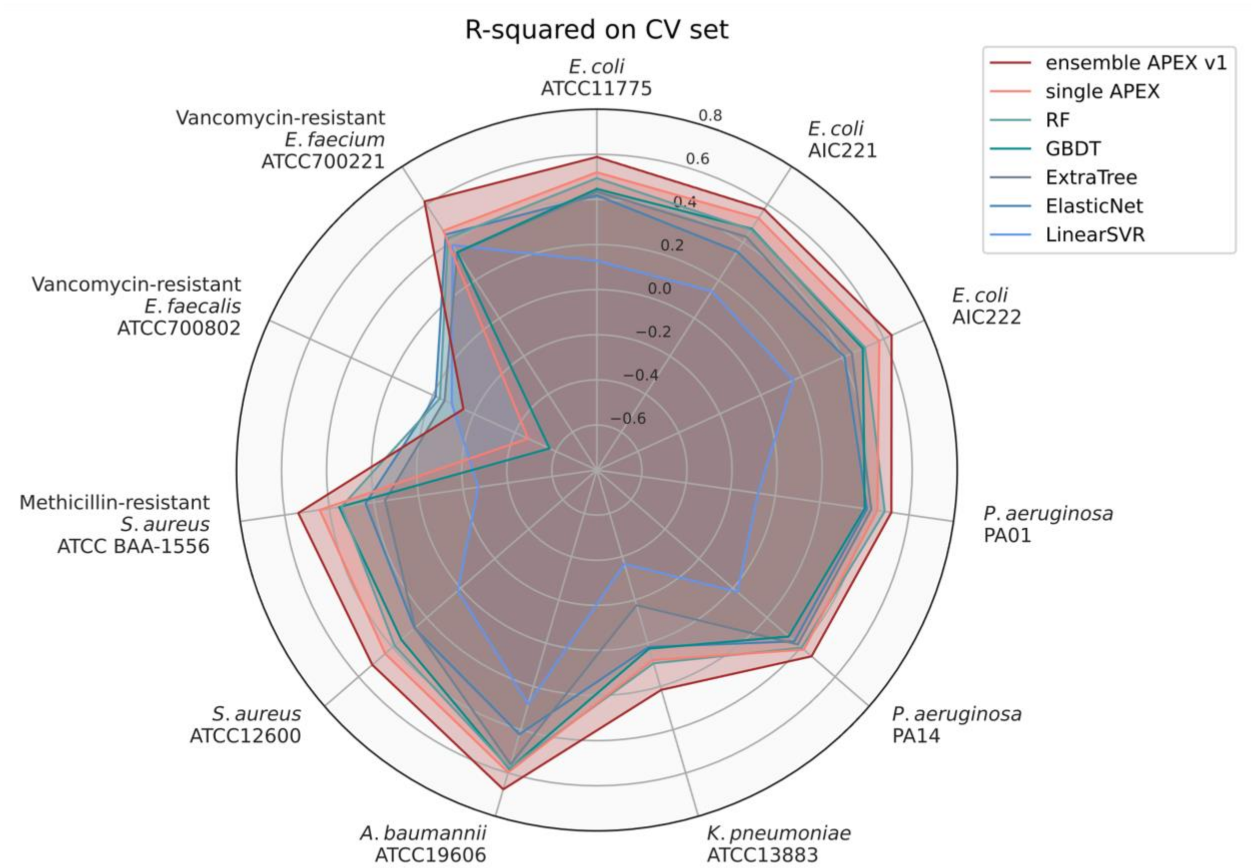


**Fig. S2. R-squared scores of various ML models on cross-validation (CV) set.** We divided our in-house peptide dataset into a CV set for hyperparameter tuning and ML model selection, and an independent dataset to evaluate the prediction performance. The figure shows the average species-specific prediction performance of 5-fold CV in terms of R-squared for various ML models on the CV set. The radius reflects the R-squared value. RF: random forest; GBDT: gradient boosting decision tree; ExtraTree: extra-tree regressor; ElasticNet: elastic net; LinearSVR: linear support vector regression.

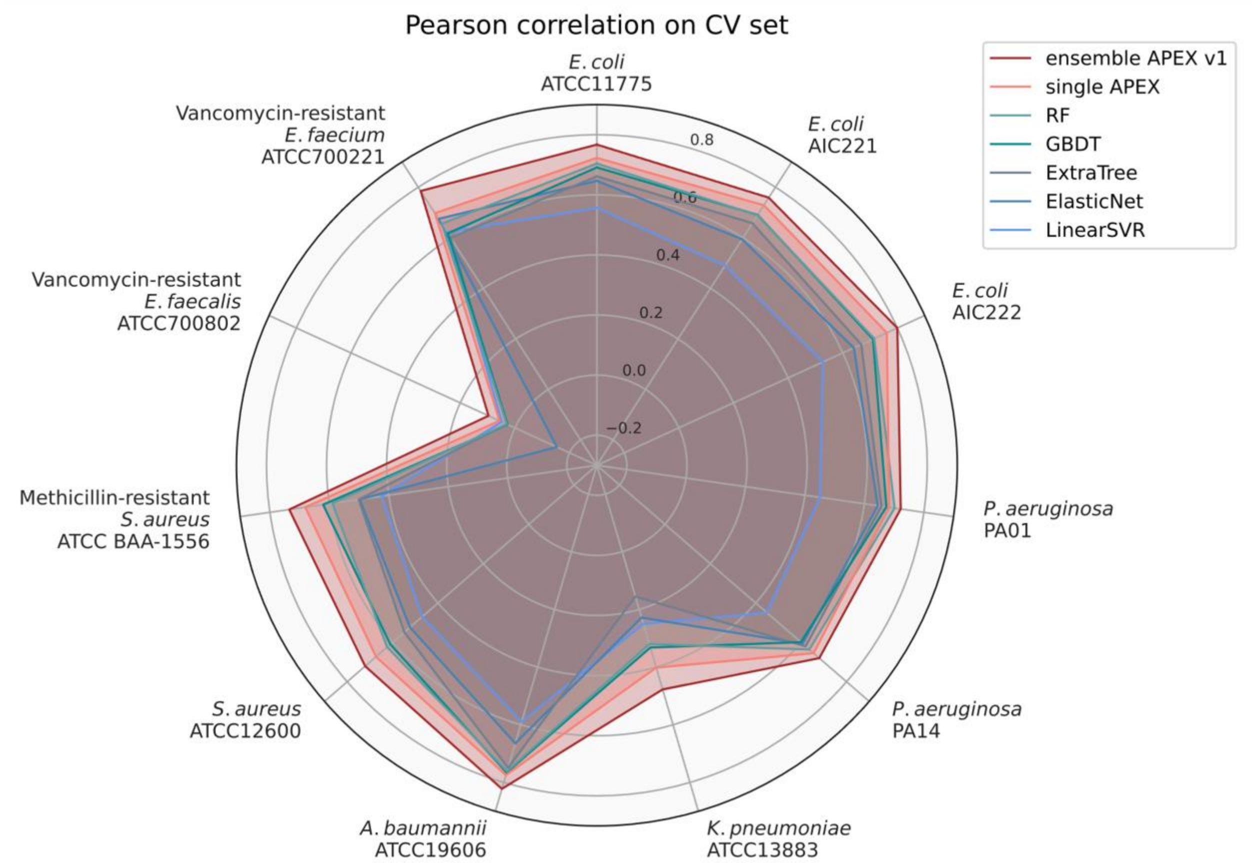


**Fig. S3. Pearson correlation scores of various ML models on cross-validation (CV) set.** We divided our in-house peptide dataset into a CV set for hyperparameter tuning and ML model selection, and an independent dataset to evaluate the prediction performance. The figure shows the average species-specific prediction performance of 5-fold CV in terms of Pearson correlation for various ML models on the CV set. The radius reflects the Pearson correlation value. RF: random forest; GBDT: gradient boosting decision tree; ExtraTree: extra-tree regressor; ElasticNet: elastic net; LinearSVR: linear support vector regression.

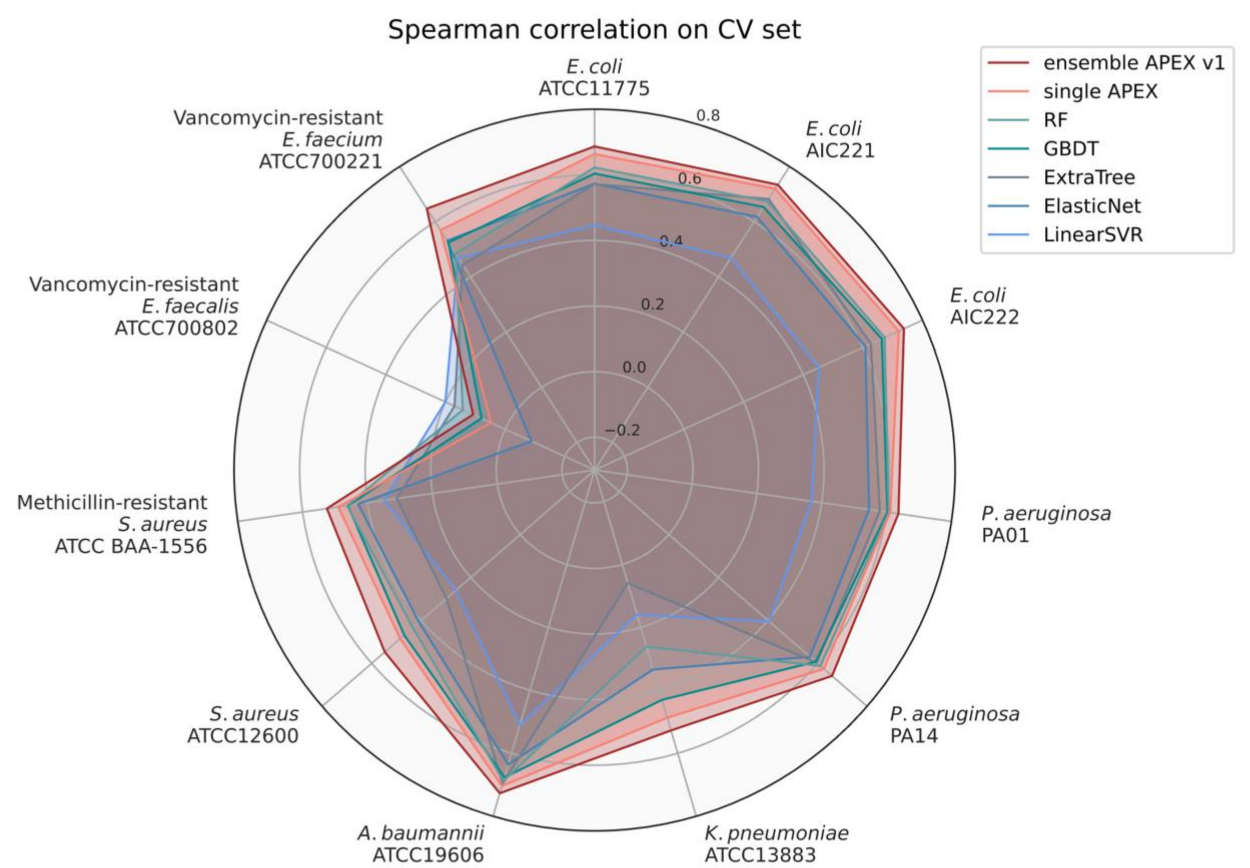


**Fig. S4. Spearman correlation scores of various ML models on cross-validation (CV) set.** We divided our in-house peptide dataset into a CV set for hyperparameter tuning and ML model selection, and an independent dataset to evaluate the prediction performance. The figure shows the average species-specific prediction performance of 5-fold CV in terms of Spearman correlation for various ML models on the CV set. The radius reflects the Spearman correlation value. RF: random forest; GBDT: gradient boosting decision tree; ExtraTree: extra-tree regressor; ElasticNet: elastic net; LinearSVR: linear support vector regression.

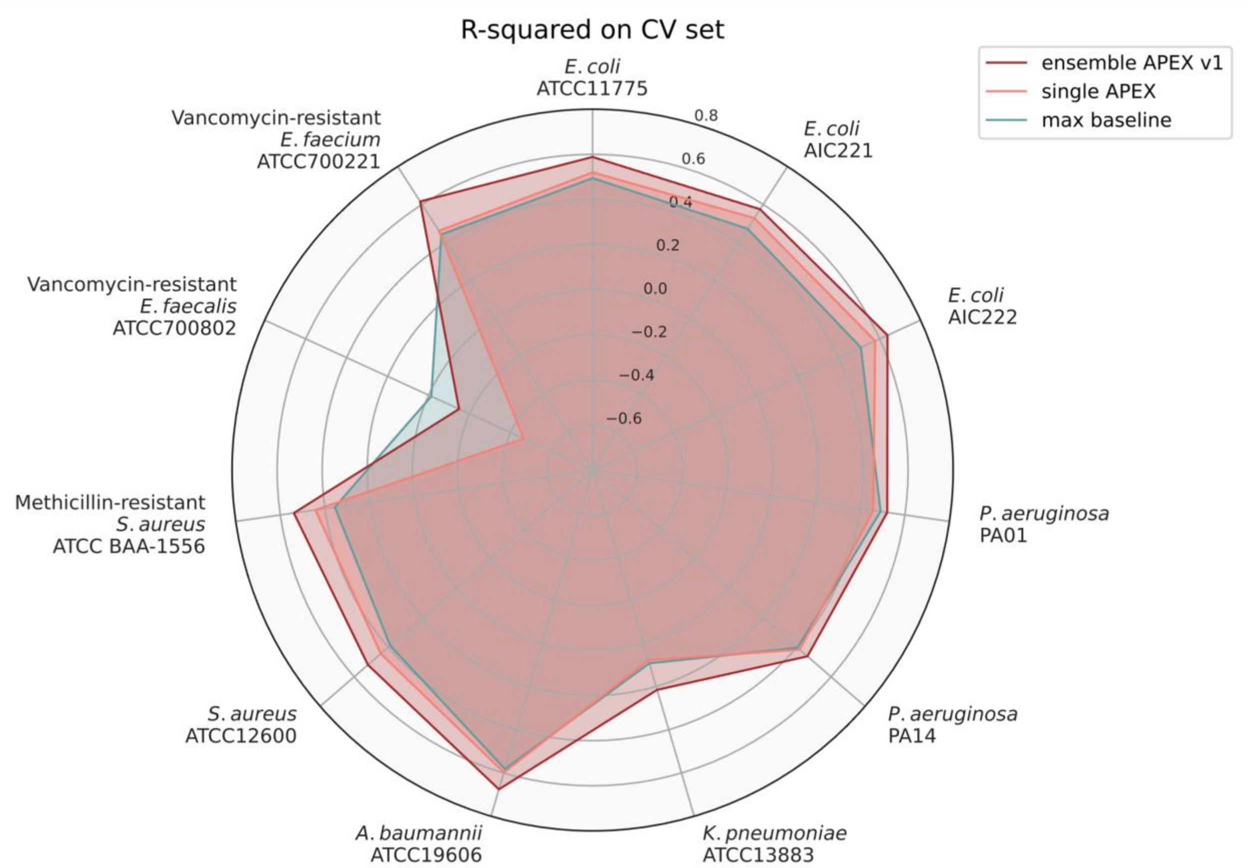


**Fig. S5. R-squared scores of various ML models on cross-validation (CV) set.** We divided our in-house peptide dataset into a CV set for hyperparameter tuning and ML model selection, and an independent dataset to evaluate the prediction performance. The figure shows the average species-specific prediction performance of 5-fold CV in terms of R-squared for APEX models and maximum performance from baseline ML models on the CV set. The radius reflects the R-squared value. Max baseline denotes the highest R-squared values from baseline ML models, including elastic net, linear support vector regression, extra-trees regressor, random forest and gradient boosting decision tree.

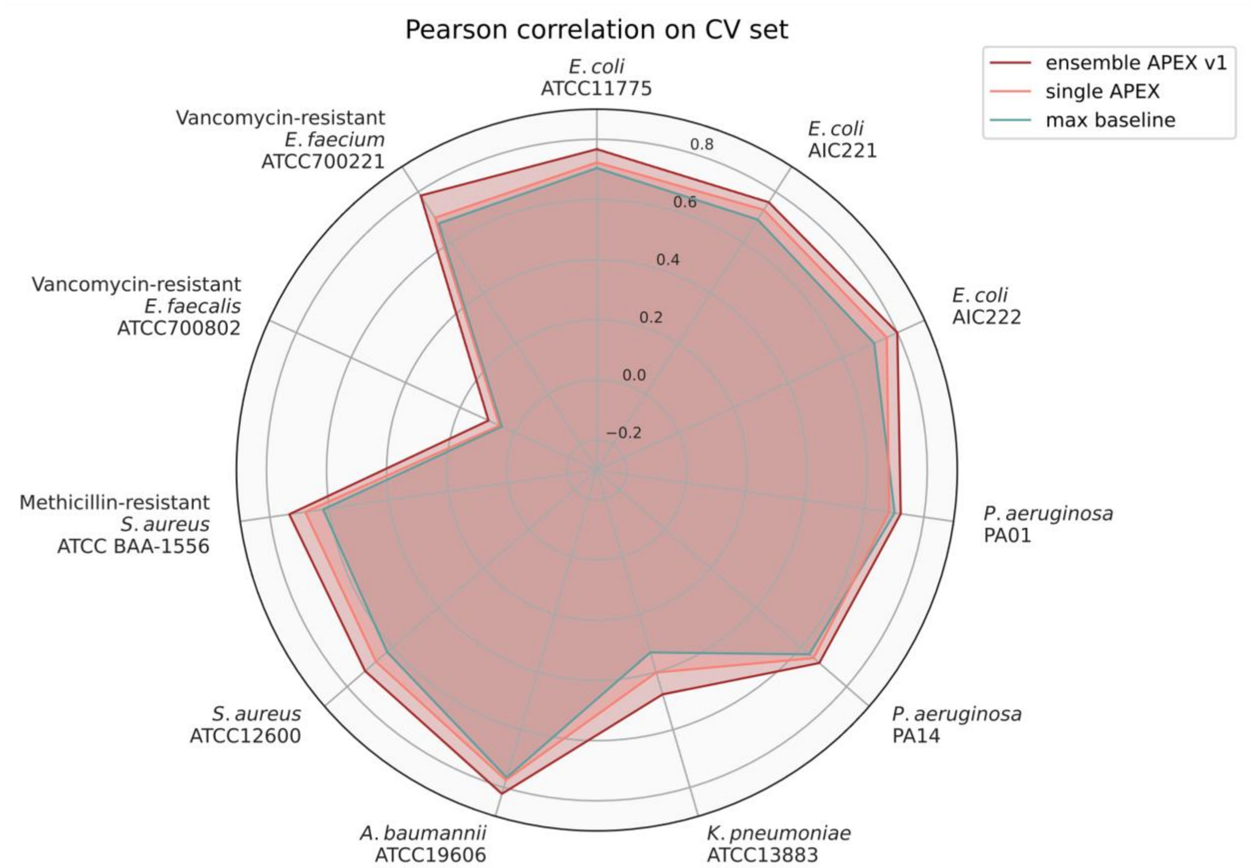


**Fig. S6. Pearson correlation scores of various ML models on CV set.** We divided our in-house peptide dataset into a CV set for hyperparameter tuning and ML model selection, and an independent dataset to evaluate the prediction performance. The figure shows the average species-specific prediction performance of 5-fold CV in terms of Pearson correlation for APEX models and maximum performance from baseline ML models on the CV set. The radius reflects the Pearson correlation value. Max baseline denotes the highest Pearson correlation values from baseline ML models, including elastic net, linear support vector regression, extra-trees regressor, random forest and gradient boosting decision tree.

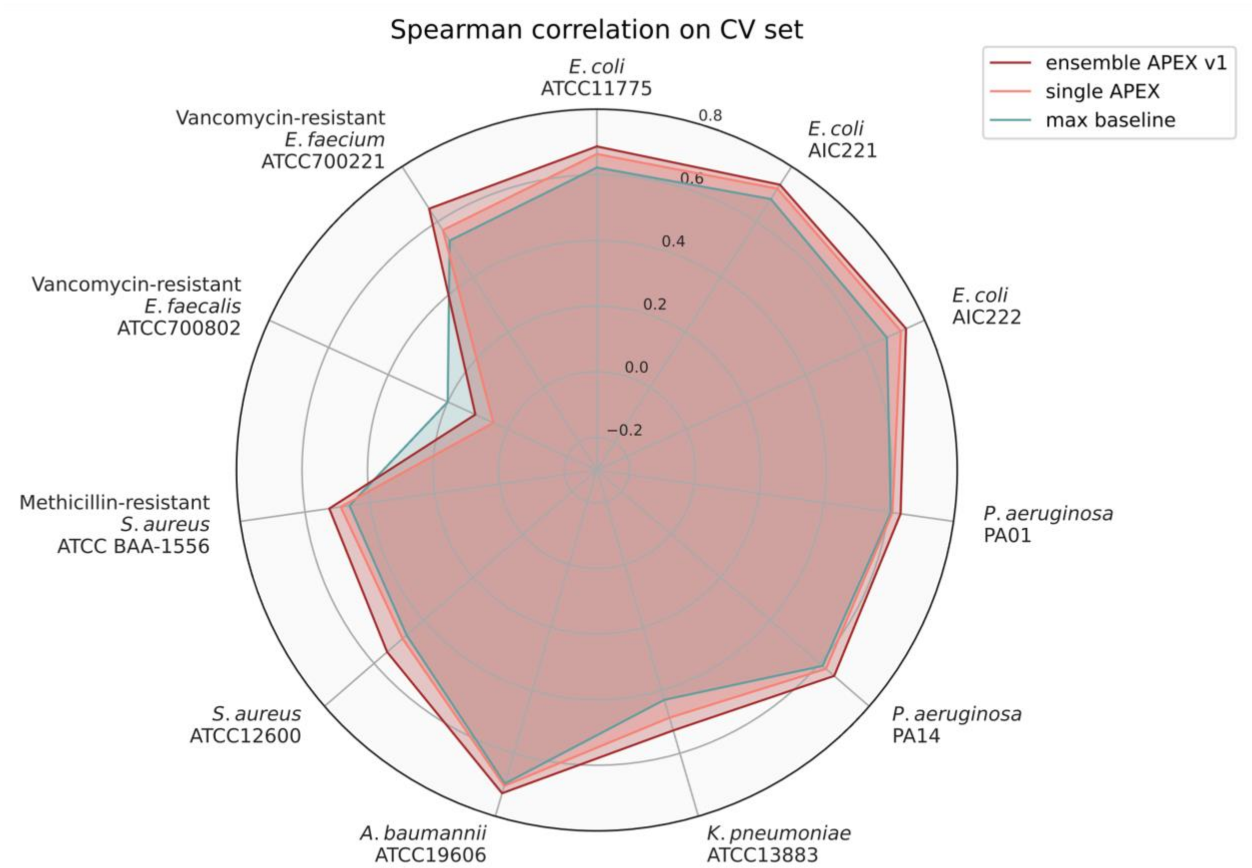


**Fig. S7. Spearman correlation scores of various ML models on CV set.** We divided our in-house peptide dataset into a CV set for hyperparameter tuning and ML model selection, and an independent dataset to evaluate the prediction performance. The figure shows the average species-specific prediction performance of 5-fold CV in terms of Spearman correlation for APEX models and maximum performance from baseline ML models on the CV set. The radius reflects the Spearman correlation value. Max baseline denotes the highest Spearman correlation values from baseline ML models, including elastic net, linear support vector regression, extra-trees regressor, random forest and gradient boosting decision tree.

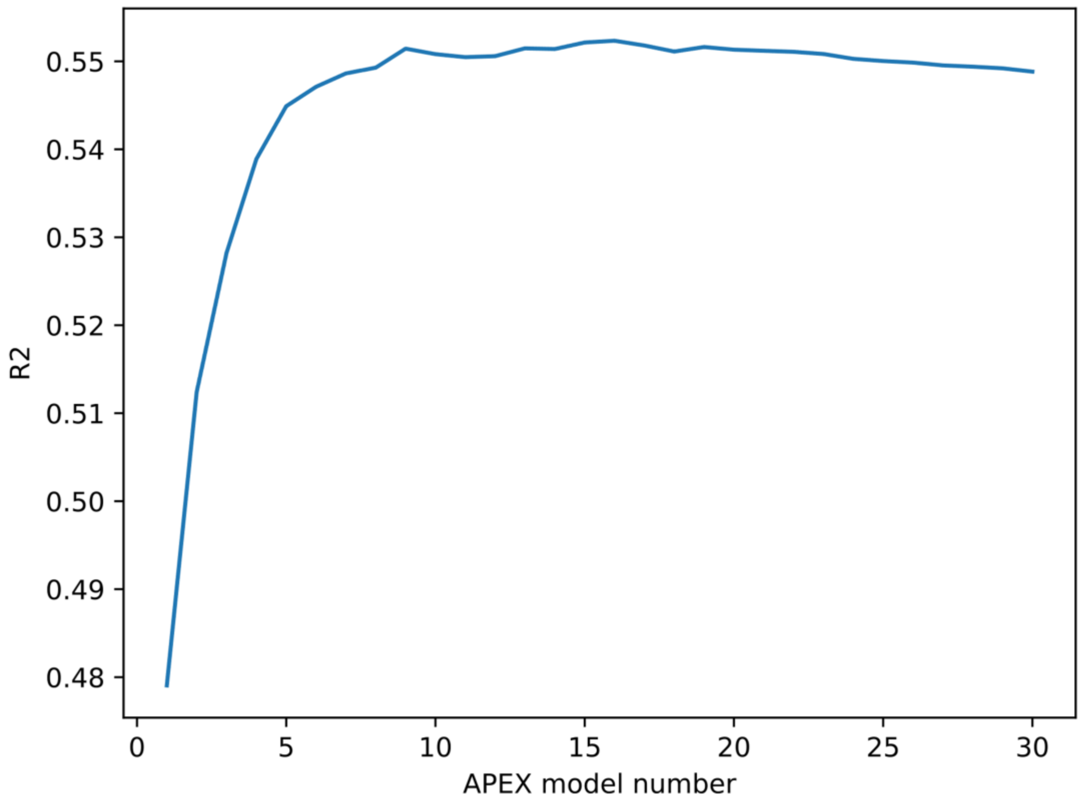


**Fig. S8. Relationship between R-squared and the number of APEX models used in ensemble learning on the CV set.** We divided our in-house peptide dataset into a CV set for hyperparameter tuning and ML model selection, and an independent dataset to evaluate prediction performance. On the CV set, we trained several APEX models under different deep learning architectures and training strategies. We ranked the APEX models by the averaged R-squared to predict species-specific antimicrobial activity. We averaged the predictions from the top-ranked models to create the ensemble APEX to improve antibiotic prediction performance. The figure shows the relationship between the averaged R-squared for species-specific antimicrobial activity prediction and the number of APEX models being averaged.

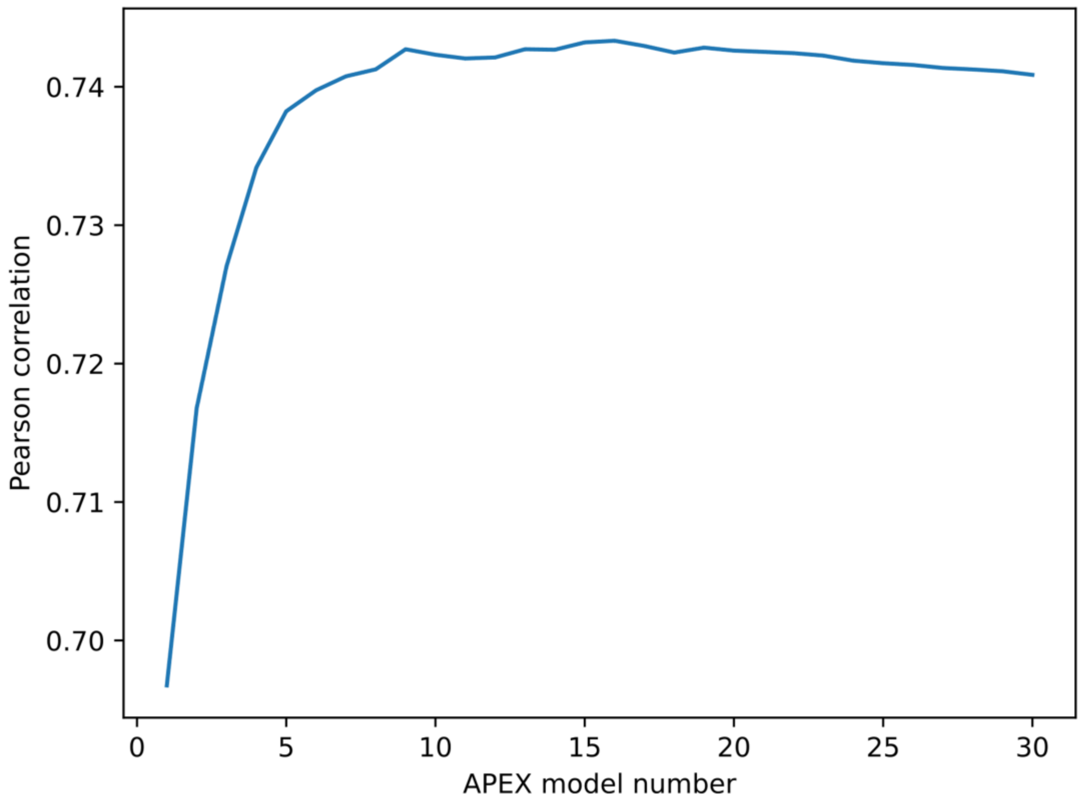


**Fig. S9. Relationship between Pearson correlation and the number of APEX models used in ensemble learning on the CV set.** We divided our in-house peptide dataset into a CV set for hyperparameter tuning and ML model selection, and an independent dataset to evaluate prediction performance. On CV set, we trained the APEX models under different deep learning architectures and training strategies, then ranked them by the averaged R-squared for species-specific antimicrobial activity prediction. To improve antibiotic prediction performance, we averaged the predictions from the top-ranked models creating ensemble APEX. The figure shows the relationship between the averaged Pearson correlation for species-specific antimicrobial activity prediction and the number of APEX models being averaged.

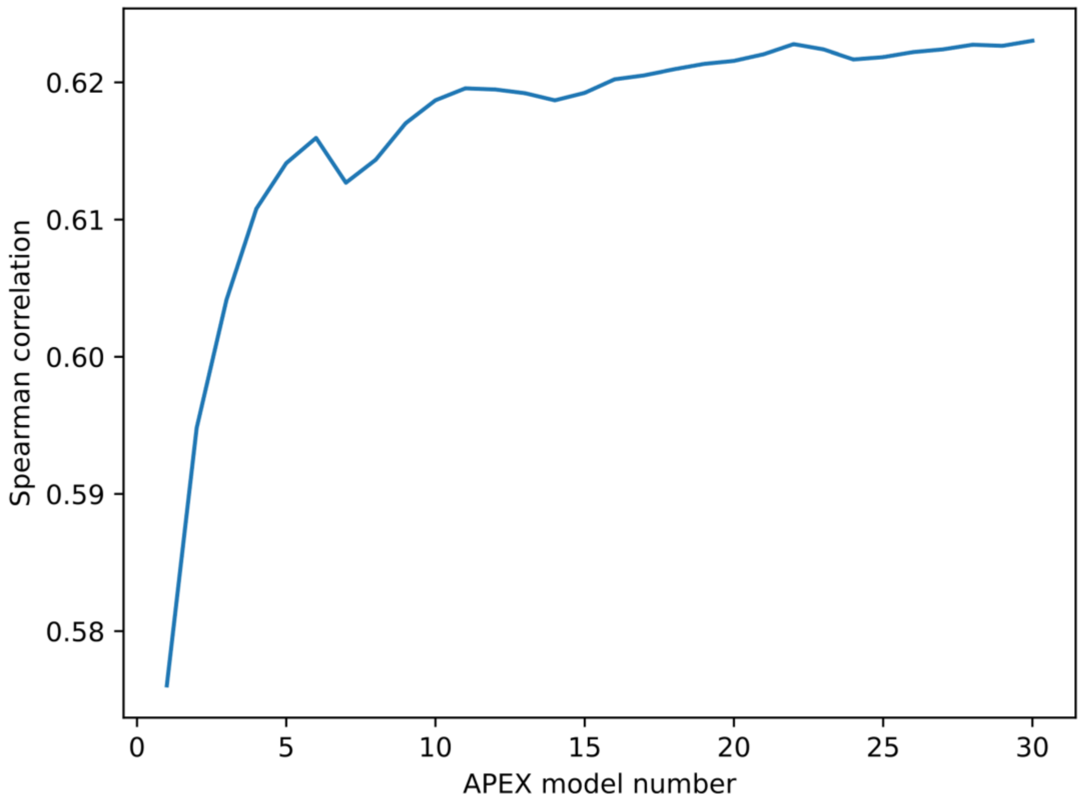


**Fig. S10. Relationship between Spearman correlation and the number of APEX models used in ensemble learning on CV set.** We divided our in-house peptide dataset into a CV set for hyperparameter tuning and ML model selection, and an independent dataset to evaluate prediction performance. On CV set, we trained APEX models under different deep learning architectures and training strategies. We ranked the APEX models by the averaged R-squared for species-specific antimicrobial activity prediction. To improve antibiotic prediction performance, we averaged the predictions from the top-ranked models to create ensemble APEX to improve antibiotic prediction performance. The figure shows the relationship between averaged Spearman correlation for species-specific antimicrobial activity prediction and the number of APEX models being averaged.

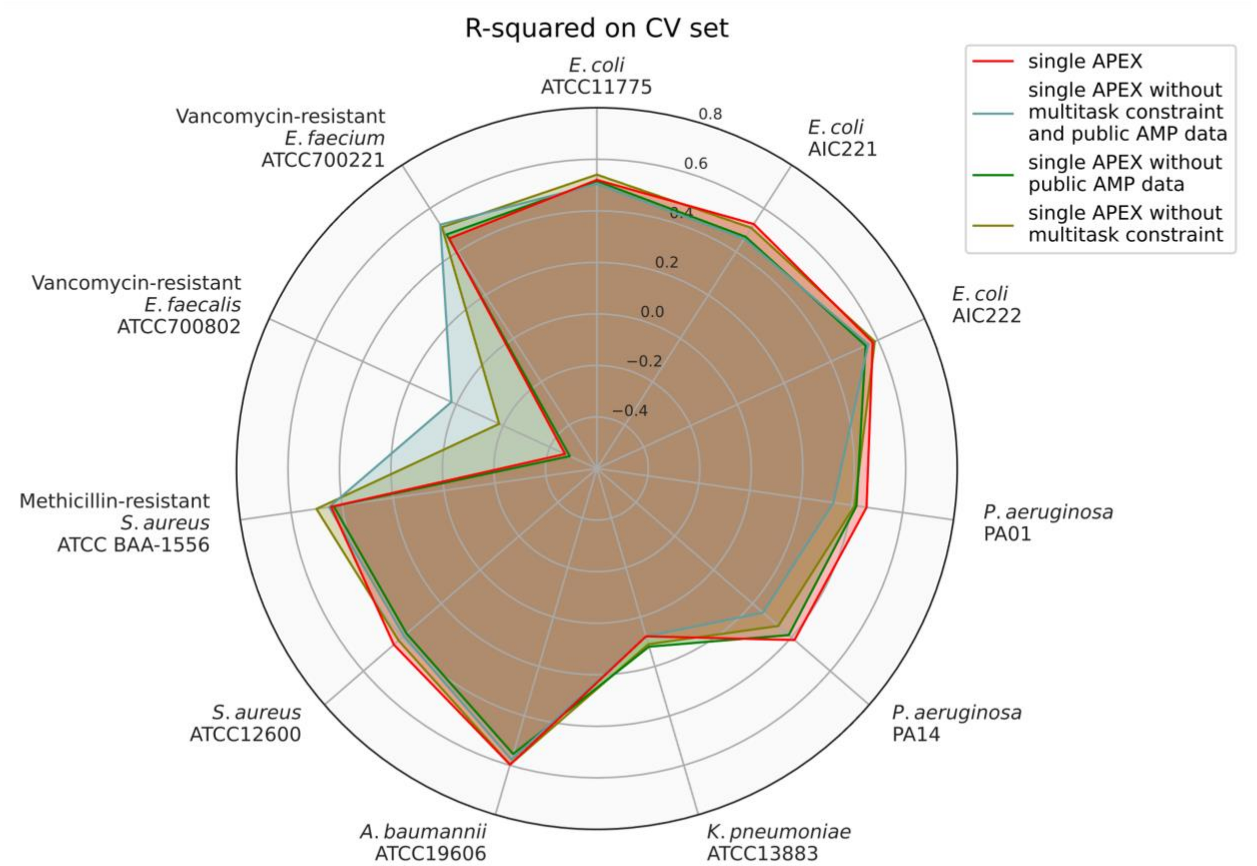


**Fig. S11. Ablation study of the multitask learning strategy of APEX in terms of R-squared on the CV set.** We divided our in-house peptide dataset into a CV set for hyperparameter tuning and ML model selection, and an independent dataset to evaluate prediction performance. The figure shows the average species-specific prediction performance of 5-fold cross validation in terms of R-squared for various APEX variants, including APEX without multitask constraint, APEX without using public AMP data during training, APEX without multitask constraint and public AMP data during training, and the original APEX (i.e., single APEX). The radius reflects the R-squared value.

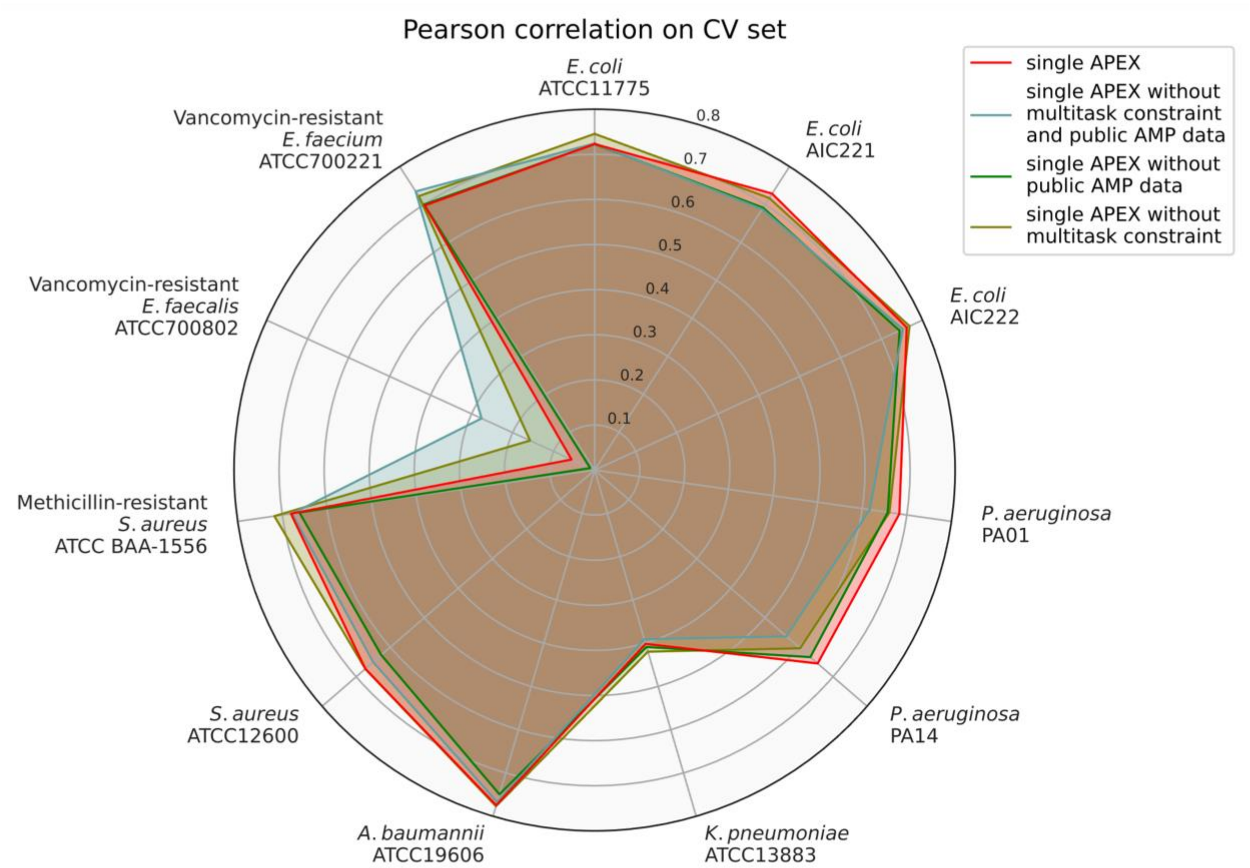


**Fig. S12. Ablation study of the multitask learning strategy of APEX in terms of Pearson correlation on CV set.** We divided our in-house peptide dataset into a CV set for hyperparameter tuning and ML model selection, and an independent dataset to evaluate prediction performance. The figure shows the average species-specific prediction performance of 5-fold cross validation in terms of Pearson correlation for various APEX variants, including APEX without multitask constraint, APEX without using public AMP data during training, APEX without multitask constraint and public AMP data during training, and the original APEX (i.e., single APEX). The radius reflects the Pearson correlation value.

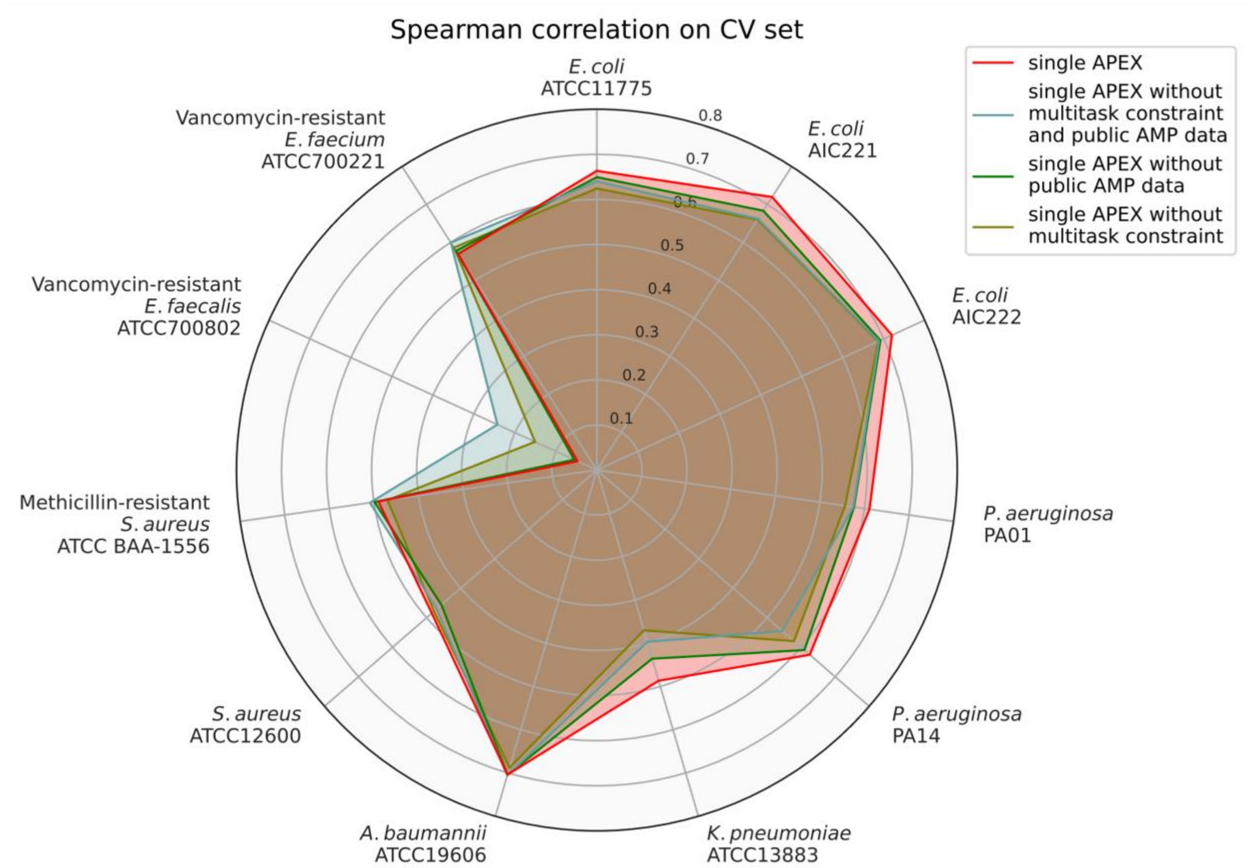


**Fig. S13. Ablation study of the multitask learning strategy of APEX in terms of Spearman correlation on CV set.** We divided our in-house peptide dataset into a CV set for hyperparameter tuning and ML model selection, and an independent dataset to evaluate prediction performance. The figure shows the average species-specific prediction performance of 5-fold cross validation in terms of Spearman correlation for various APEX variants, including APEX without multitask constraint, APEX without using public AMP data during training, APEX without multitask constraint and public AMP data during training, and the original APEX (i.e., single APEX). The radius reflects the Spearman correlation value.

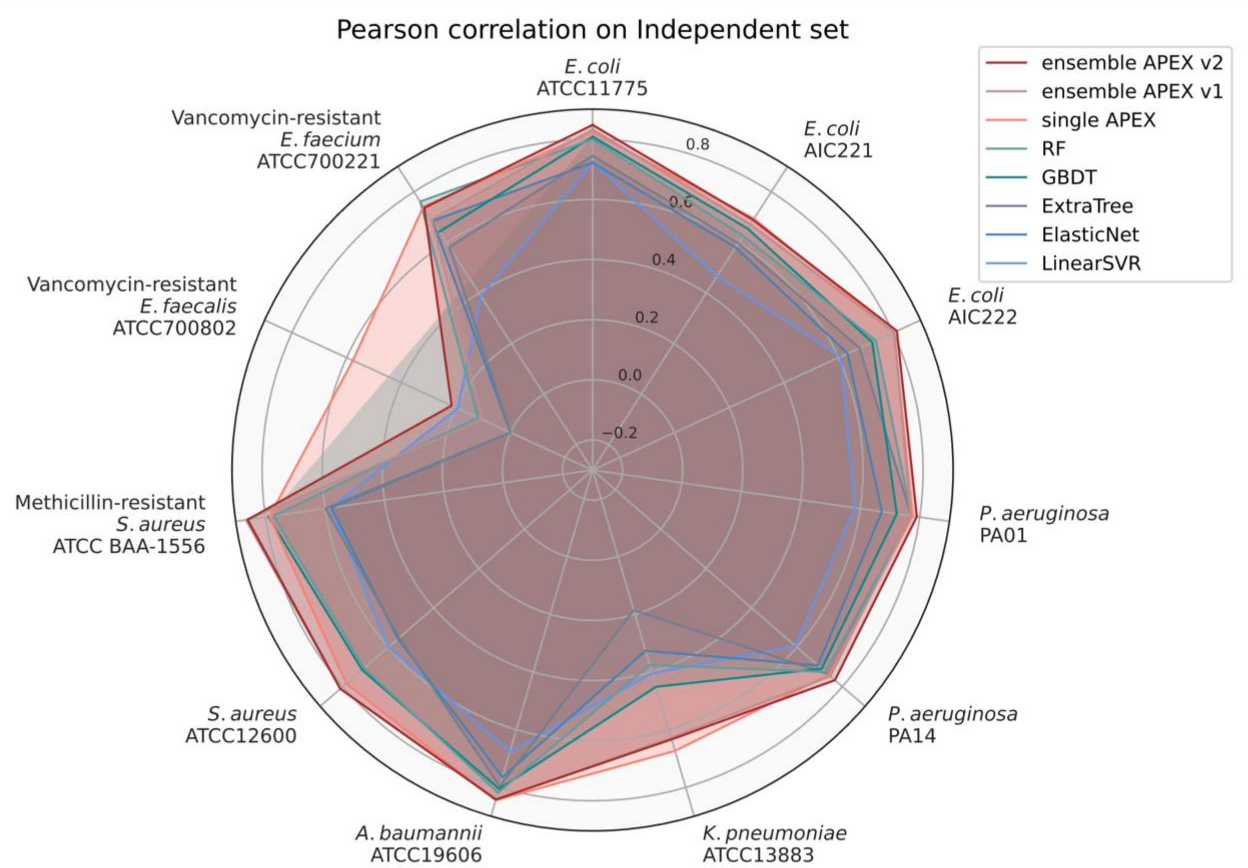


**Fig. S14. Pearson correlation scores of various ML models on an independent set.** We divided our in-house peptide dataset into a CV set for hyperparameter tuning and ML model selection, and an independent dataset to evaluate prediction performance. The figure shows the species-specific prediction performance in terms of Pearson correlation for various ML models that were trained on the CV set and evaluated on the independent set. The radius reflects the Pearson correlation value. RF: random forest; GBDT: gradient boosting decision tree; ExtraTree: extra-tree regressor; ElasticNet: elastic net; LinearSVR: linear support vector regression.

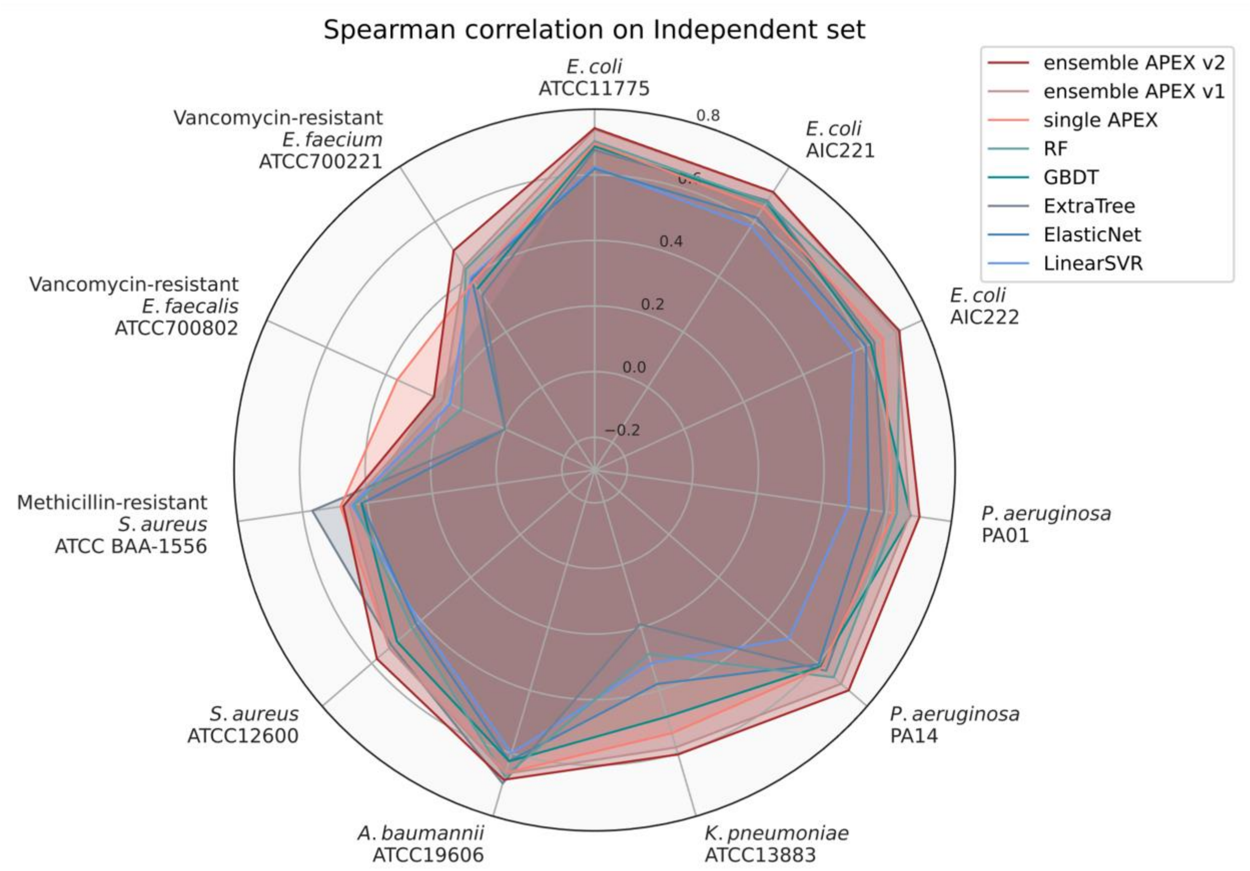


**Fig. S15. Spearman correlation scores of various ML models on an independent set.** We divided our in-house peptide dataset into a CV set for hyperparameter tuning and ML model selection, and an independent dataset to evaluate prediction performance. The figure shows the species-specific prediction performance in terms of Spearman correlation for various ML models that were trained on the CV set and evaluated on the independent set. The radius reflects the Spearman correlation value. RF: random forest; GBDT: gradient boosting decision tree; ExtraTree: extra-tree regressor; ElasticNet: elastic net; LinearSVR: linear support vector regression.

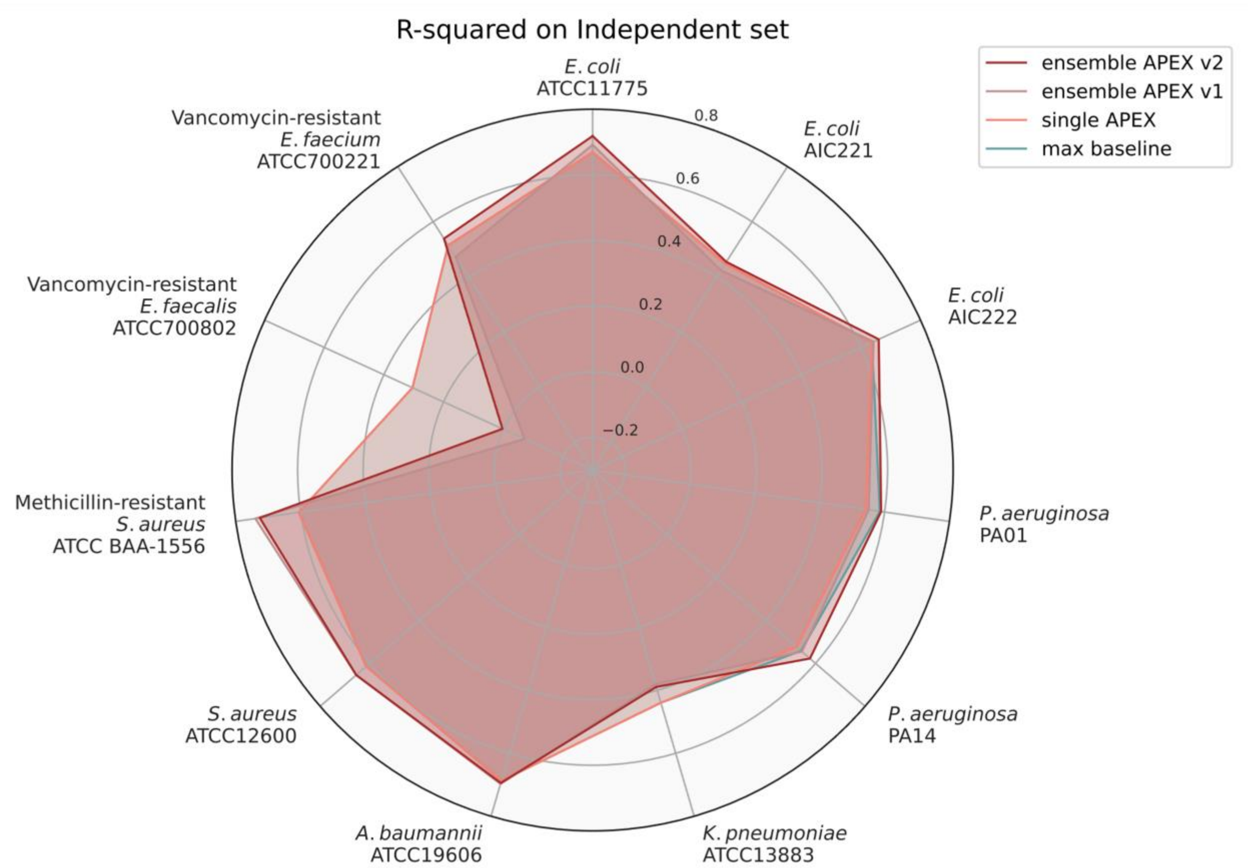


**Fig. S16. R-squared scores of various ML models on an independent set.** We divided our in-house peptide dataset into a CV set for hyperparameter tuning and ML model selection, and an independent dataset to evaluate prediction performance. The figure shows the species-specific prediction performance in terms of R-squared for APEX models and maximum performance from baseline ML models that were trained on the CV set and evaluated on the independent set. The radius reflects the R-squared value. Max baseline denotes the highest R-squared values from baseline ML models, including elastic net, linear support vector regression, extra-trees regressor, random forest and gradient boosting decision tree.

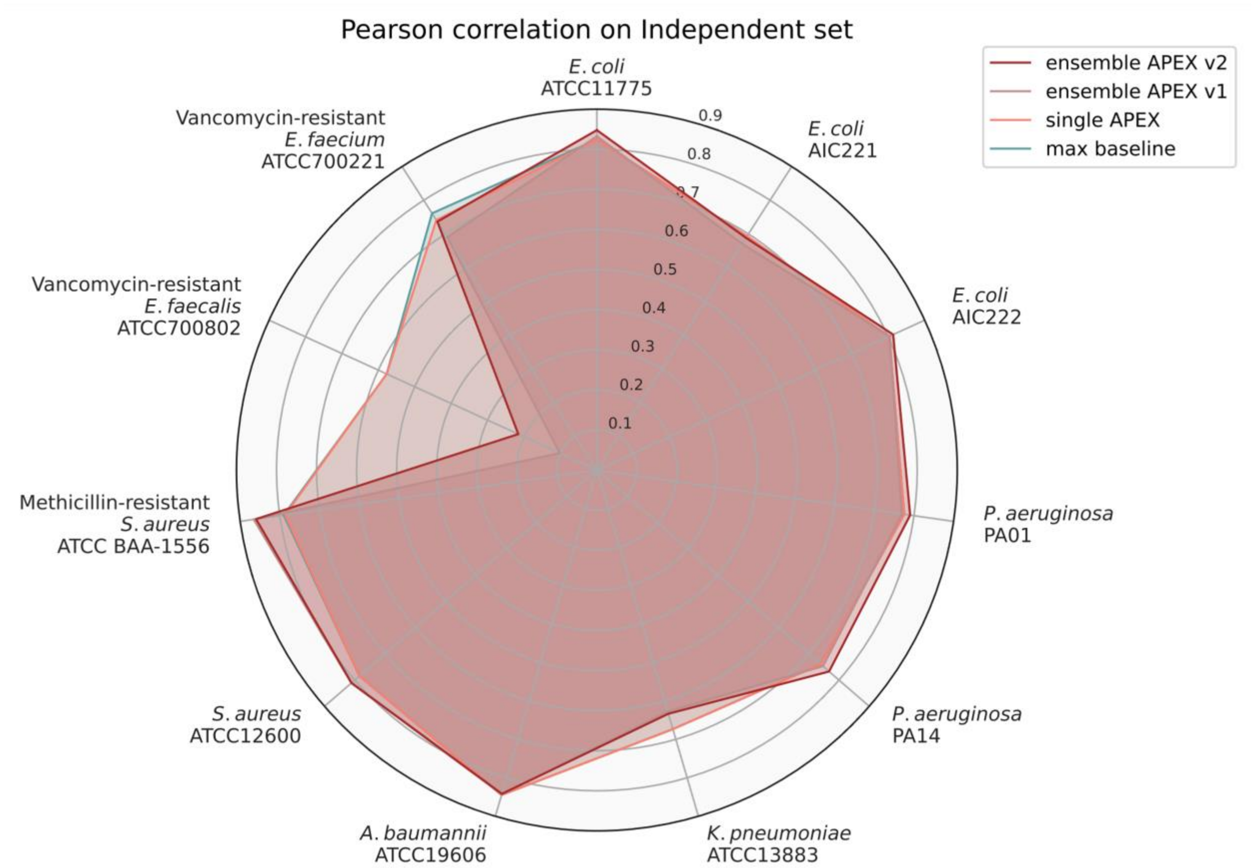


**Fig. S17. Pearson correlation scores of various ML models on an independent set.** We divided our in-house peptide dataset into a CV set for hyperparameter tuning and ML model selection, and an independent dataset to evaluate prediction performance. The figure shows the species-specific prediction performance in terms of Pearson correlation for APEX models and maximum performance from baseline ML models that were trained on the CV set and evaluated on the independent set. The radius reflects the Pearson correlation value. Max baseline denotes the highest Pearson correlation values from baseline ML models, including elastic net, linear support vector regression, extra-trees regressor, random forest and gradient boosting decision tree.

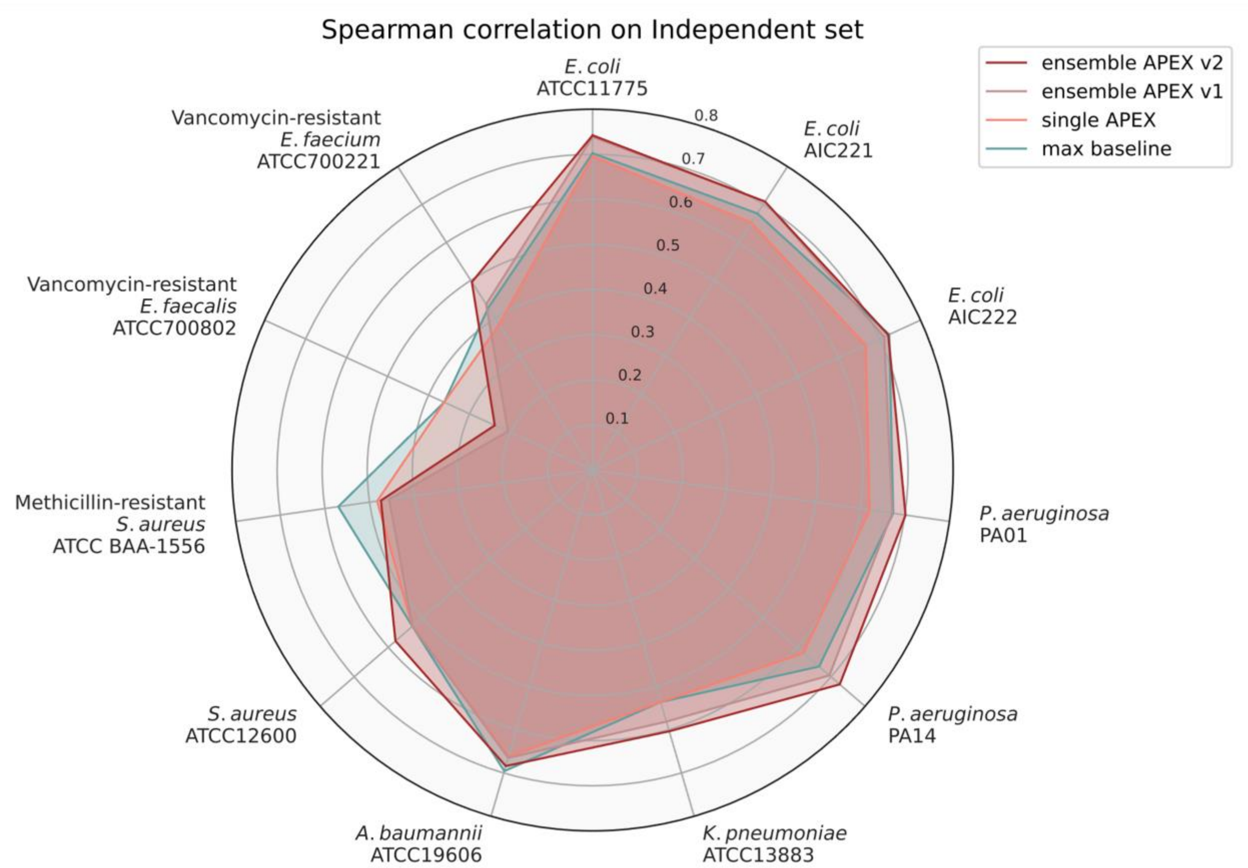


**Fig. S18. Spearman correlation scores of various ML models on an independent set.** We divided our in-house peptide dataset into a CV set for hyperparameter tuning and ML model selection, and an independent dataset to evaluate prediction performance. The figure shows the species-specific prediction performance in terms of Spearman correlation for APEX models and maximum performance from baseline ML models that were trained on the CV set and evaluated on the independent set. The radius reflects the Spearman correlation value. Max baseline denotes the highest Spearman correlation values from baseline ML models, including elastic net, linear support vector regression, extra-trees regressor, random forest and gradient boosting decision tree.

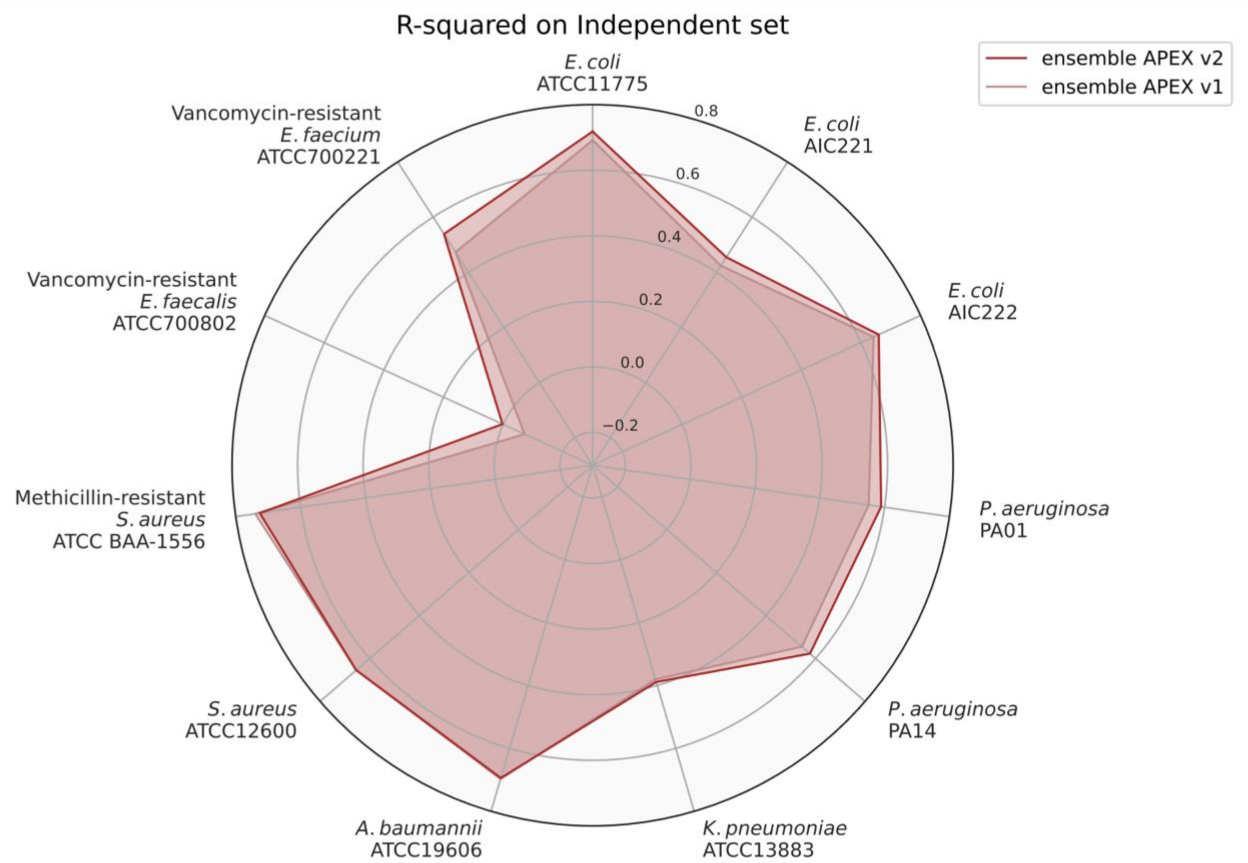


**Fig. S19. R-squared scores of ensemble APEX v2 and v1 on an independent set.** We divided our in-house peptide dataset into a CV set for hyperparameter tuning and ML model selection, and an independent dataset to evaluate prediction performance. The figure shows the species-specific prediction performance in terms of R-squared for two APEX variants that were trained on the CV set and evaluated on the independent set. The ensemble APEX v1 averaged the predictions from 8 different neural network architectures and training strategies. On top of ensemble APEX v1, ensemble APEX v2 further trained 5 copies under different random seeds for each base learner from ensemble APEX v1 to create 8×5 = 40 deep neural network predictors. The predictions from the 40 models were averaged to create the final prediction for ensemble APEX v2. The radius reflects the R-squared value.

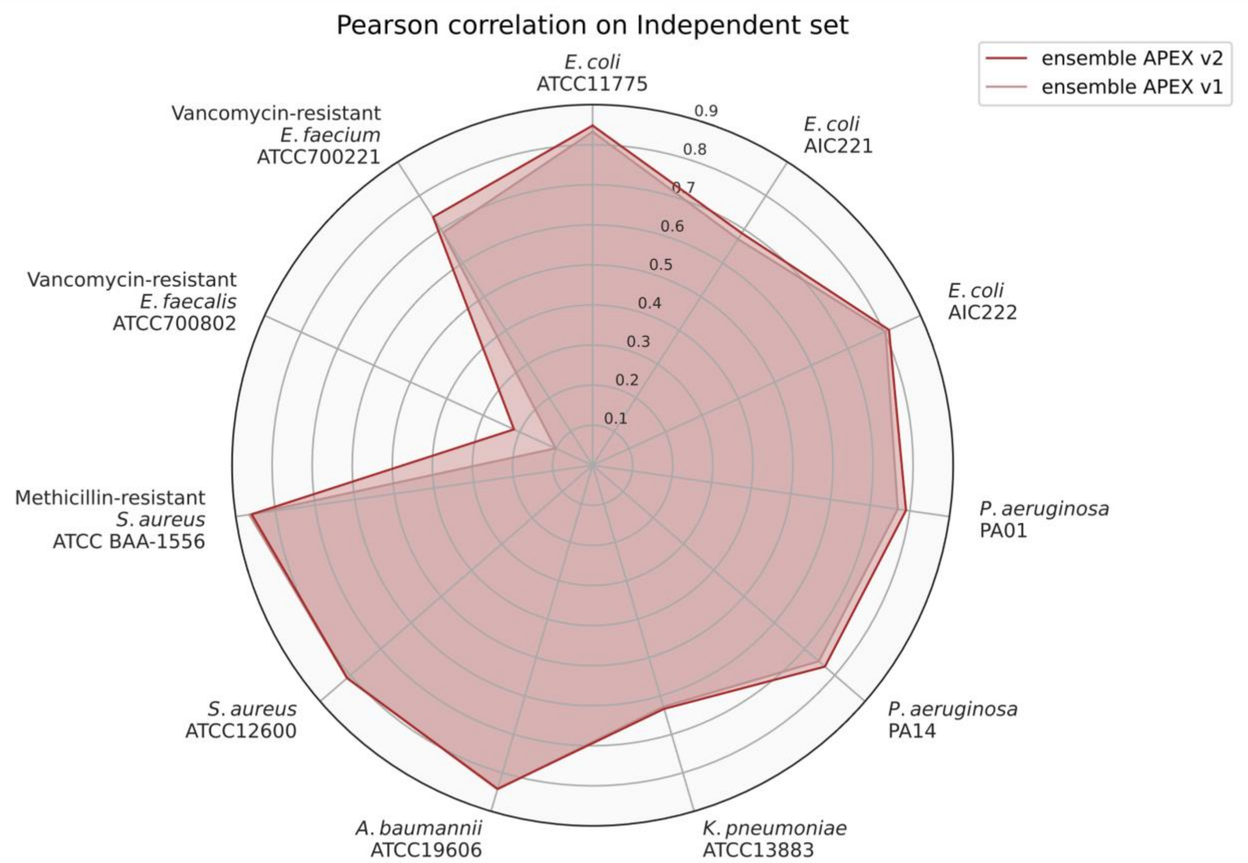


**Fig. S20. Pearson correlation scores of ensemble APEX v2 and v1 on an independent set.** We divided our in-house peptide dataset into a CV set for hyperparameter tuning and ML model selection, and an independent dataset to evaluate prediction performance. The figure shows the species-specific prediction performance in terms of Pearson correlation for two APEX variants that were trained on the CV set and evaluated on the independent set. The ensemble APEX v1 averaged the predictions from 8 different neural network architectures and training strategies. On top of ensemble APEX v1, ensemble APEX v2 further trained 5 copies under different random seeds for each base learner from ensemble APEX v1 to create 8×5 = 40 deep neural network predictors. The predictions from the 40 models were averaged to create the final prediction for ensemble APEX v2. The radius reflects the Pearson correlation value.

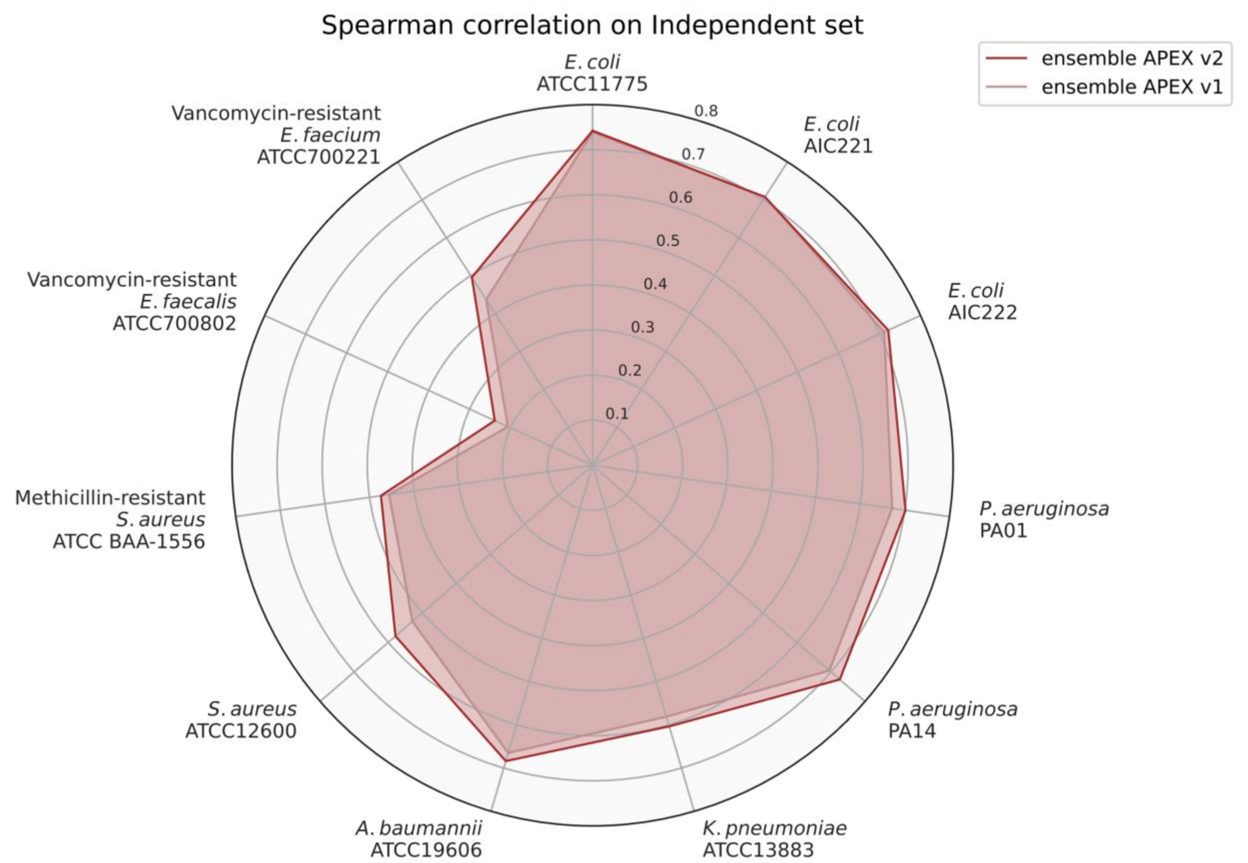


**Fig. S21. Spearman correlation scores of ensemble APEX v2 and v1 on an independent set.** We divided our in-house peptide dataset into a CV set for hyperparameter tuning and ML model selection, and an independent dataset to evaluate prediction performance. The figure shows the species-specific prediction performance in terms of Spearman correlation for two APEX variants that were trained on the CV set and evaluated on the independent set. The ensemble APEX v1 averaged the predictions from 8 different neural network architectures and training strategies. On top of ensemble APEX v1, ensemble APEX v2 further trained 5 copies under different random seeds for each base learner from ensemble APEX v1 to create 8×5 = 40 deep neural network predictors. The predictions from the 40 models were averaged to create the final prediction for ensemble APEX v2. The radius reflects the Spearman correlation value.

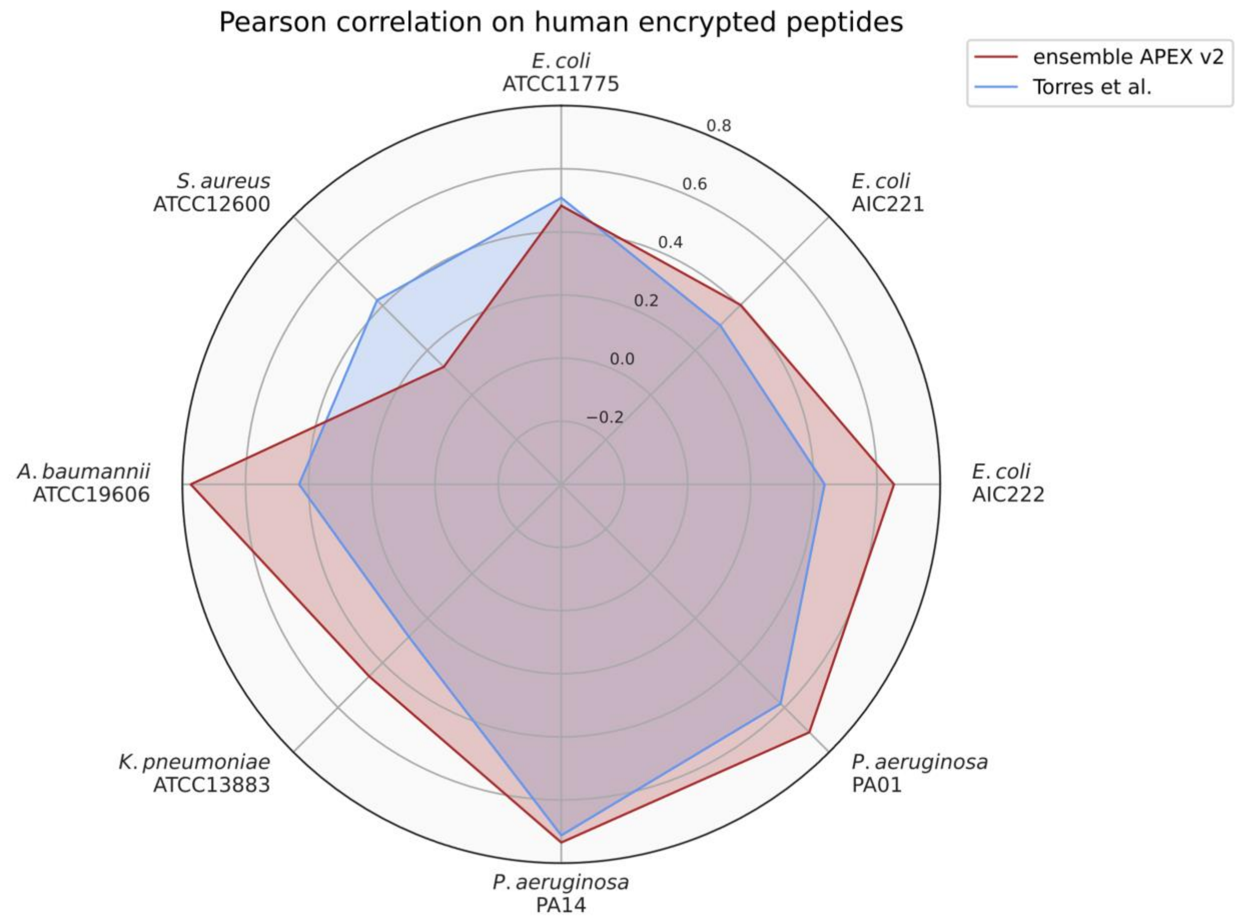


**Fig. S22. Pearson correlation scores of ensemble APEX v2 and the scoring function used to identify human encrypted peptides.** The human encrypted peptides from Torres et al.[1] constitute a subset of our in-house peptide dataset. We treated this subset as the test data and excluded it from ML model training. The figure shows the species-specific prediction performance in terms of Pearson correlation for ensemble APEX v2 and the AMP scoring function used by Torres et al.[1]. The radius reflects the Pearson correlation value.

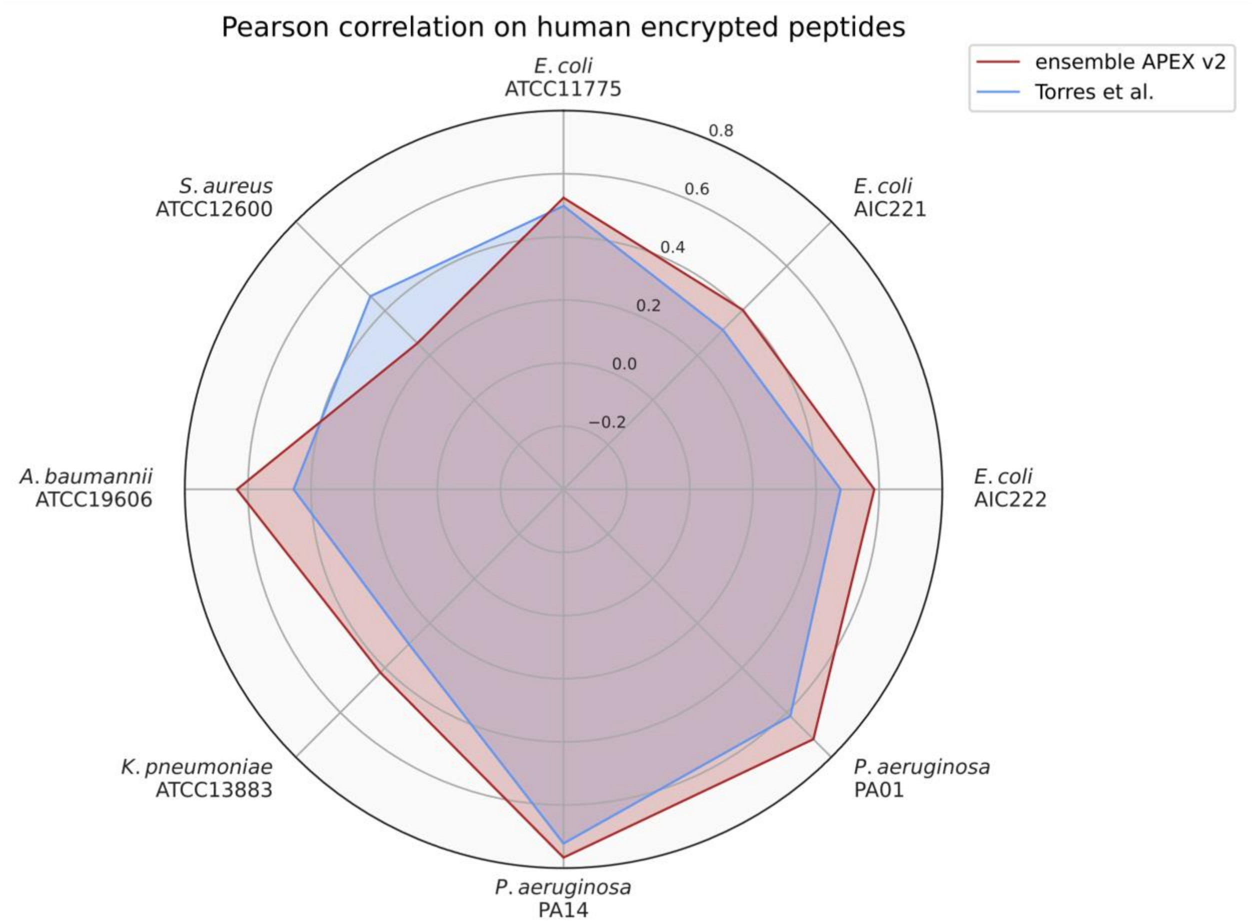


**Fig. S23. Spearman correlation scores of ensemble APEX v2 and the scoring function used to identify human encrypted peptides.** The human encrypted peptides from Torres et al.[1] constitute a subset of our in-house peptide dataset. We treated this subset as the test data and excluded it from ML model training. The figure shows the species-specific prediction performance in terms of Spearman correlation for ensemble APEX v2 and the AMP scoring function used in Torres et al.[1]. The radius reflects the Spearman correlation value.

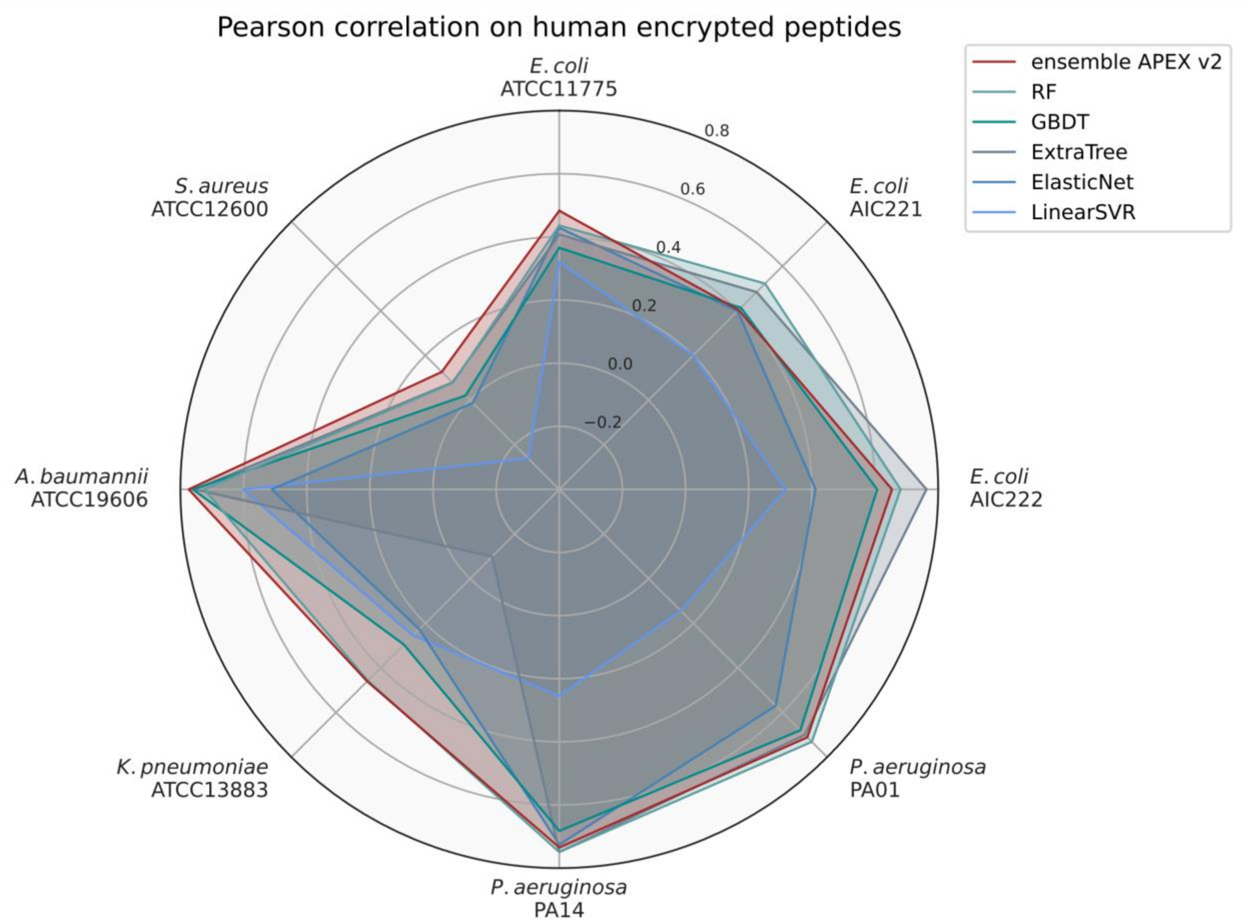


**Fig. S24. Pearson correlation scores of various ML models used to identify human encrypted peptides.** The human encrypted peptides from Torres et al.[1] constitute a subset of our in-house peptide dataset. We treated this subset as the test data and excluded it from ML model training. The figure shows the species-specific prediction performance in terms of Pearson correlation for various ML models. The radius reflects the Pearson correlation value. RF: random forest; GBDT: gradient boosting decision tree; ExtraTree: extra-tree regressor; ElasticNet: elastic net; LinearSVR: linear support vector regression.

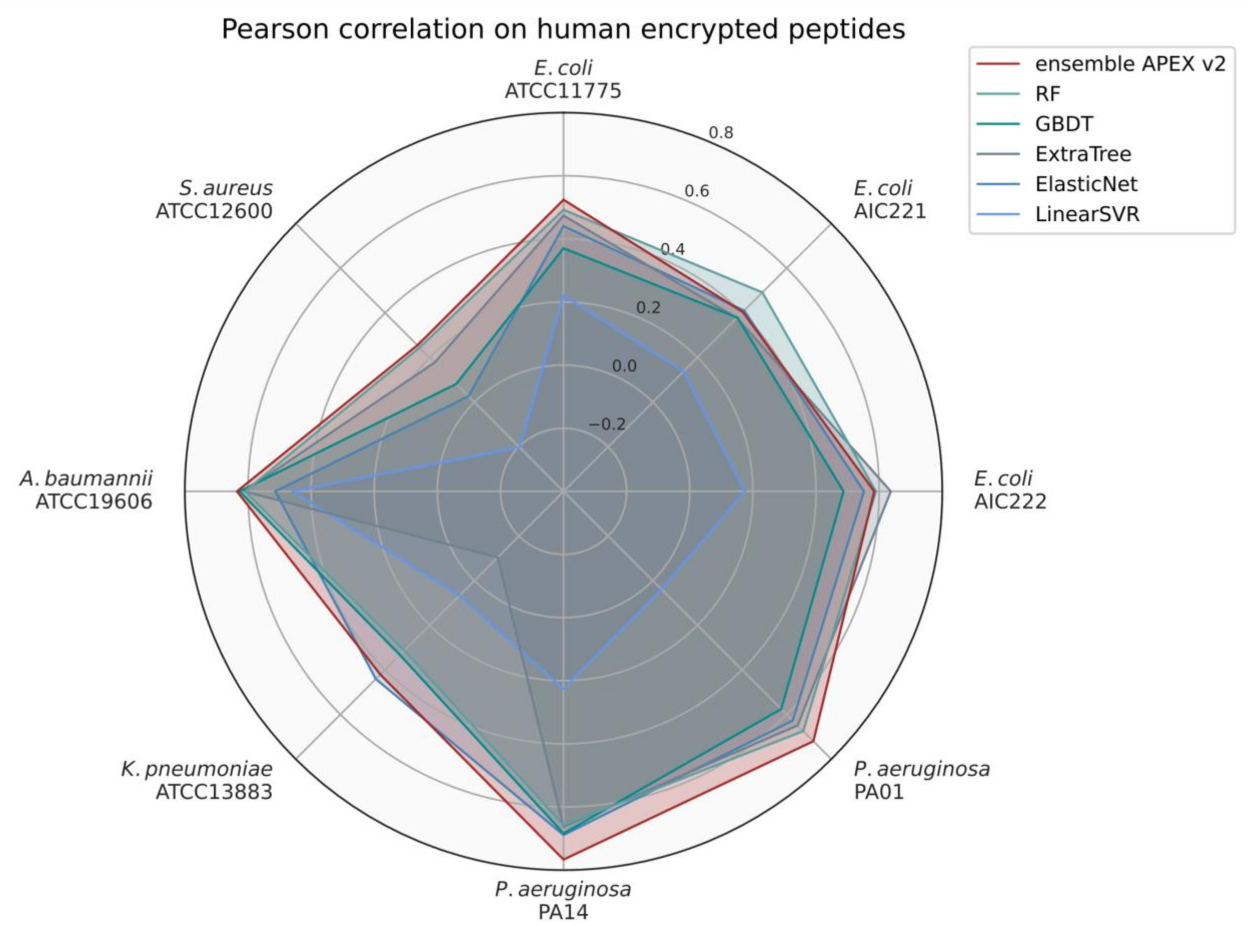


**Fig. S25. Spearman correlation scores of various ML models used to identify human encrypted peptides.** The human encrypted peptides from Torres et al.[1] constitute a subset of our in-house peptide dataset. We treated this subset as the test data and excluded it from ML model training. The figure shows the species-specific prediction performance in terms of Spearman correlation for various ML models. The radius reflects the Spearman correlation value. RF: random forest; GBDT: gradient boosting decision tree; ExtraTree: extra-tree regressor; ElasticNet: elastic net; LinearSVR: linear support vector regression.

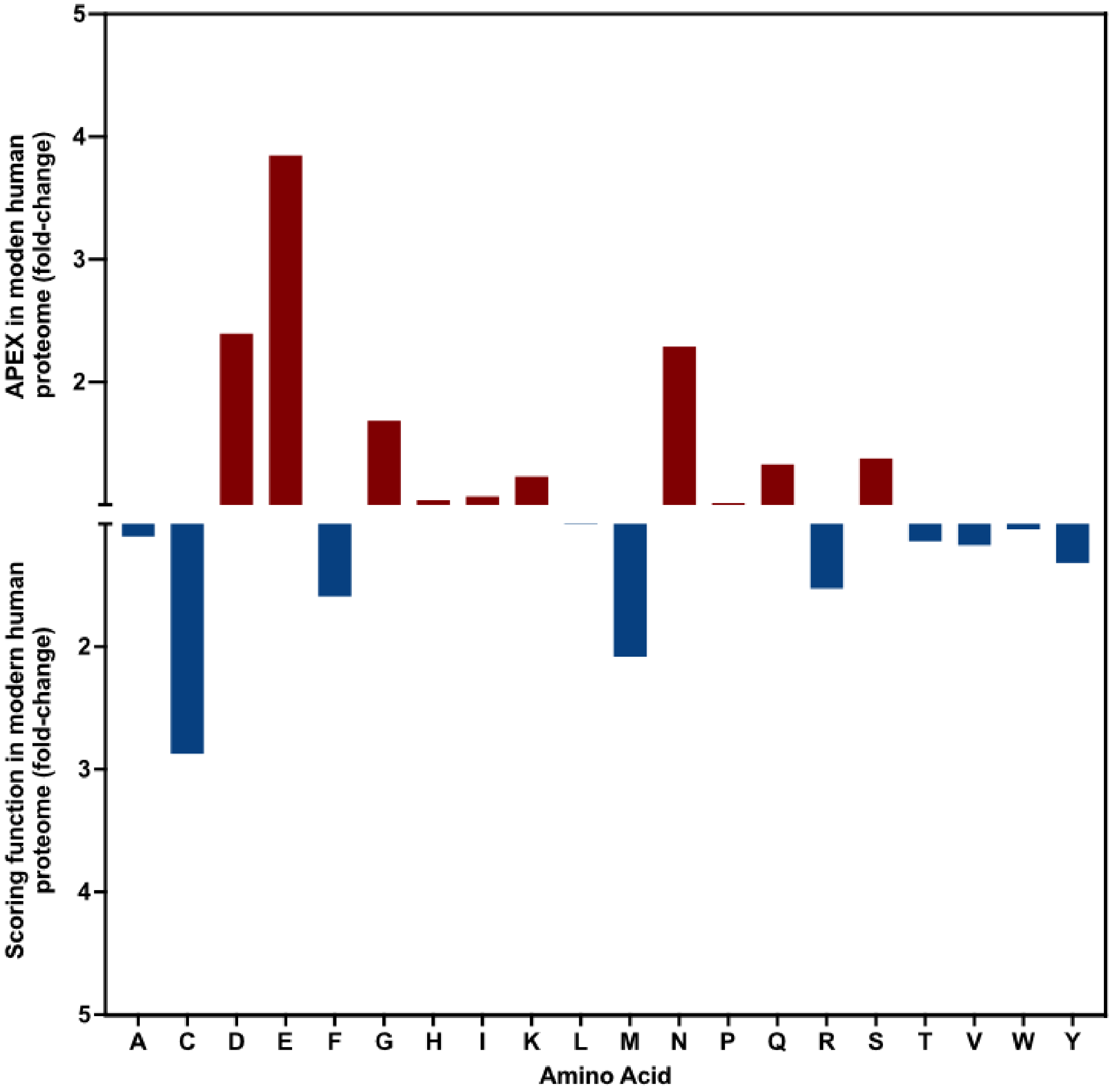


**Fig. S26. Relative abundance of the amino acid content of encrypted peptides (EPs) from the modern human proteome identified by APEX (top) and the scoring function (bottom).** The frequency of amino acid was normalized by the total number of amino acid residue counts. The ratio between normalized amino acid frequencies highlights the overrepresentation of negatively charged residues (D and E), glycine (G), and uncharged polar residues (N, Q, and S) in peptides identified by APEX. The scoring function preferably identified encrypted peptides with a high frequency of cysteine (C), methionine (M), arginine (R) and phenylalanine (F) residues.

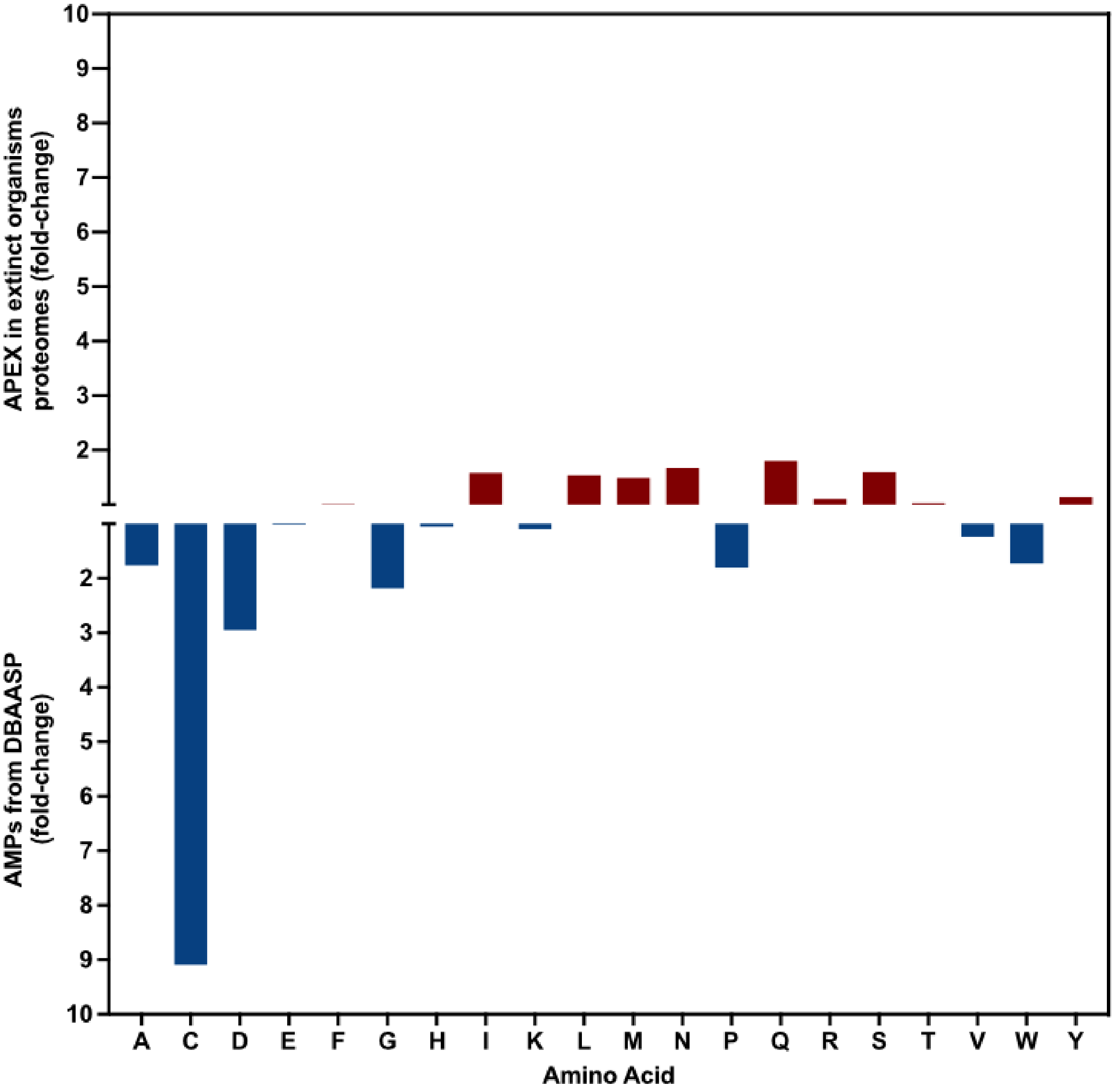


**Fig. S27. Relative abundance of the amino acid content of encrypted peptides (EPs) identified by APEX from the proteomes of extinct organisms (top) compared to known AMPs from the DBAASP (bottom).** The frequency of each amino acid residue was normalized by the total number of amino acid residue counts. The ratio between AEPs identified by APEX and known peptide from DBAASP highlights the relative abundance of aliphatic (L and I) and uncharged polar (M, N, Q, and S) residues in sequences from extinct organisms, whereas peptides from DBAASP present a higher content of negatively charged (D) and aromatic (W) residues, as well as residues known for their role in secondary structure (C, G, P).

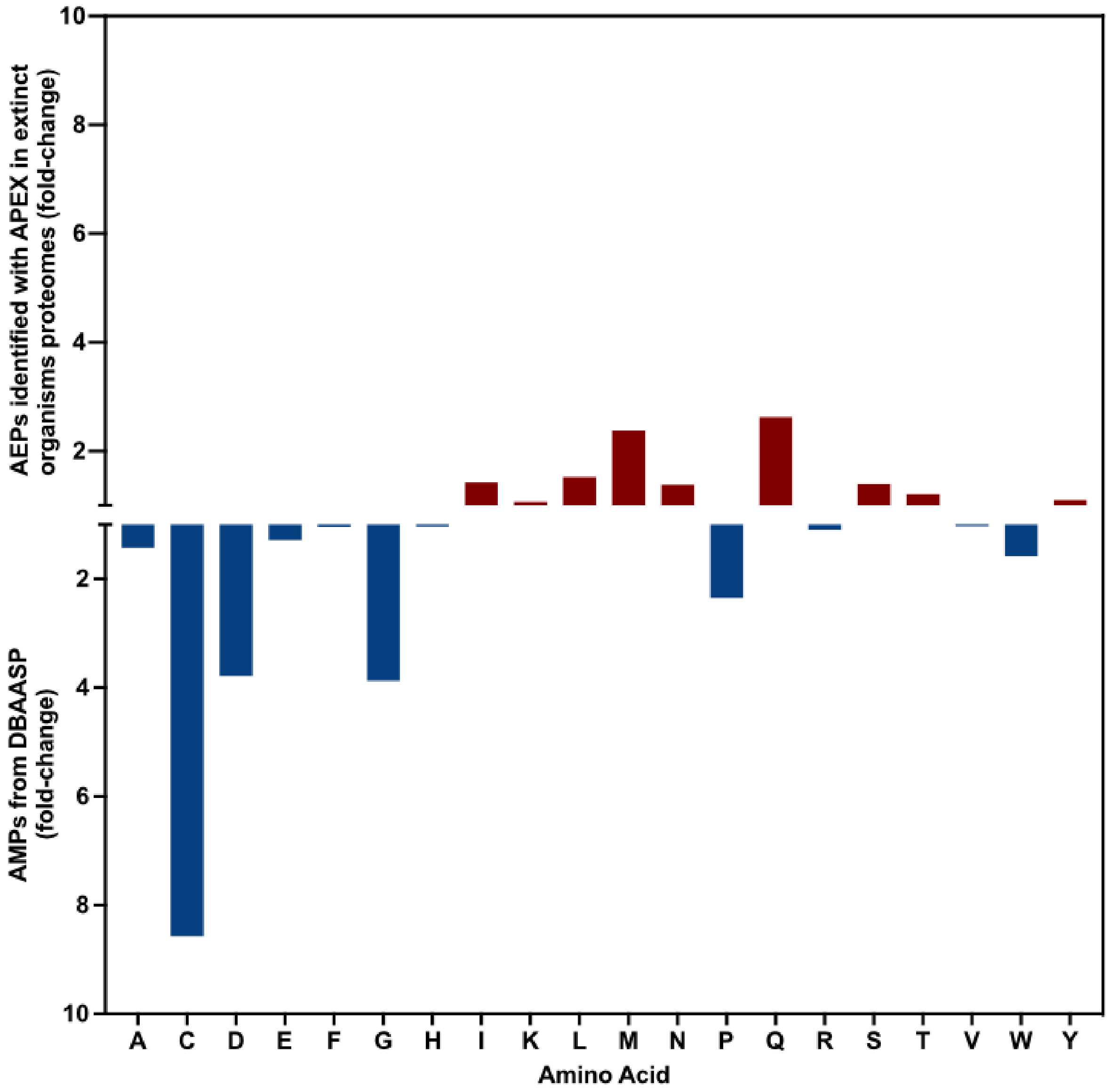


**Fig. S28. Relative abundance of the amino acid content of archaic encrypted peptides (AEPs) identified by APEX from the proteomes of extinct organisms (top) compared to known AMPs from the DBAASP (bottom).** The frequency of each amino acid was normalized by the total number of amino acid residue counts. The ratio between encrypted peptides from extinct proteins identified by APEX and known peptides from the DBAASP highlights the relative abundance of aliphatic (L and I) and uncharged polar (M, N, Q, and S) residues in sequences from extinct proteins, whereas peptides from the DBAASP had a higher content of negatively charged (D and E) and aromatic (W) residues, as well as residues known for their role in secondary structure (A, C, G, P).

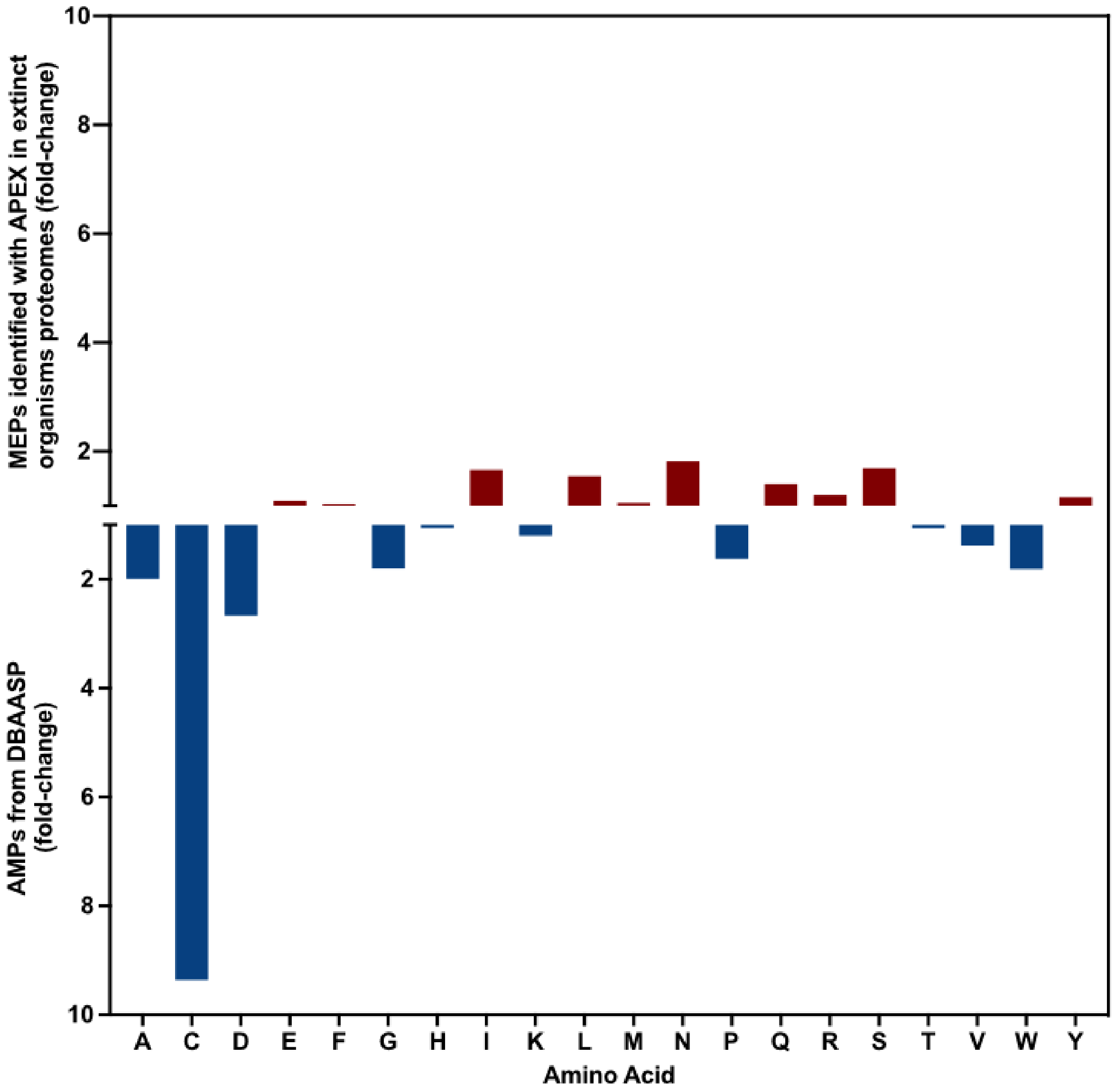


**Fig. S29. Relative abundance of the amino acid content of MEPs identified by APEX from the proteomes of extinct organisms compared to known AMPs from DBAASP.** The frequency of amino acid was normalized by the total number of amino acid residue counts. The ratio between encrypted peptides from extinct organisms that still exist in modern organisms proteomes identified by APEX and known peptide from DBAASP highlights the relative abundance of aliphatic (L and I) and polar non-charged (M, N, Q, and S) residues in sequences from extinct organisms proteins, whereas peptides from DBAASP present a higher content of negatively charged (D), aromatic (W), and residues known for their role on secondary structure (A, C, G, P).

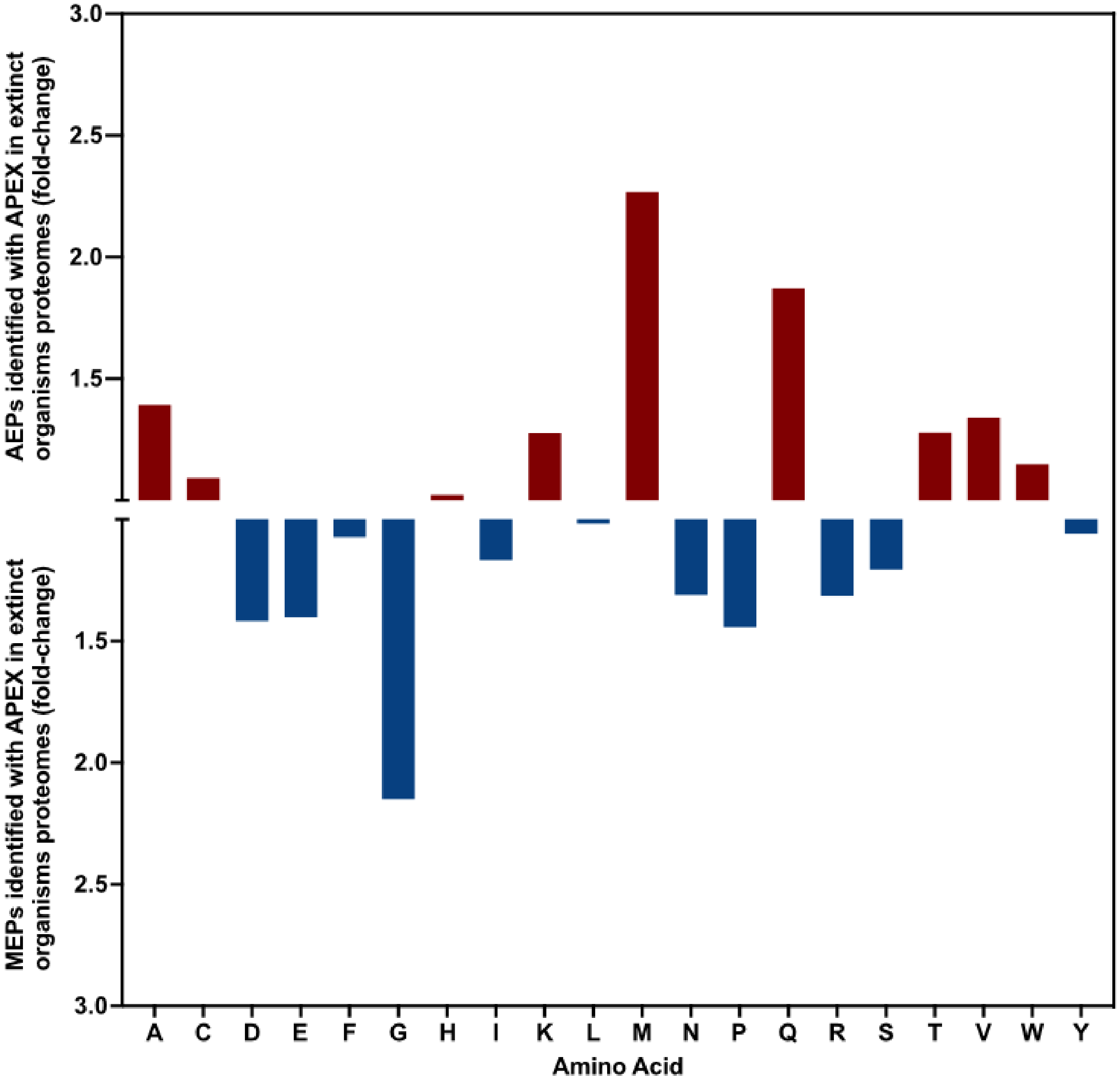


**Fig. S30. Relative abundance of the amino acid content of AEPs and MEPs identified by APEX from the proteomes of extinct organisms.** The frequency of amino acid was normalized by the total number of amino acid residue counts. The ratio between encrypted peptides from extinct proteins that are not present in modern organisms and from proteins that still exist in modern organisms identified by APEX highlights the relative abundance of polar non-charged (M and Q) residues in sequences from extinct proteins, whereas encrypted peptides from still existing proteins present a higher content of glycine.

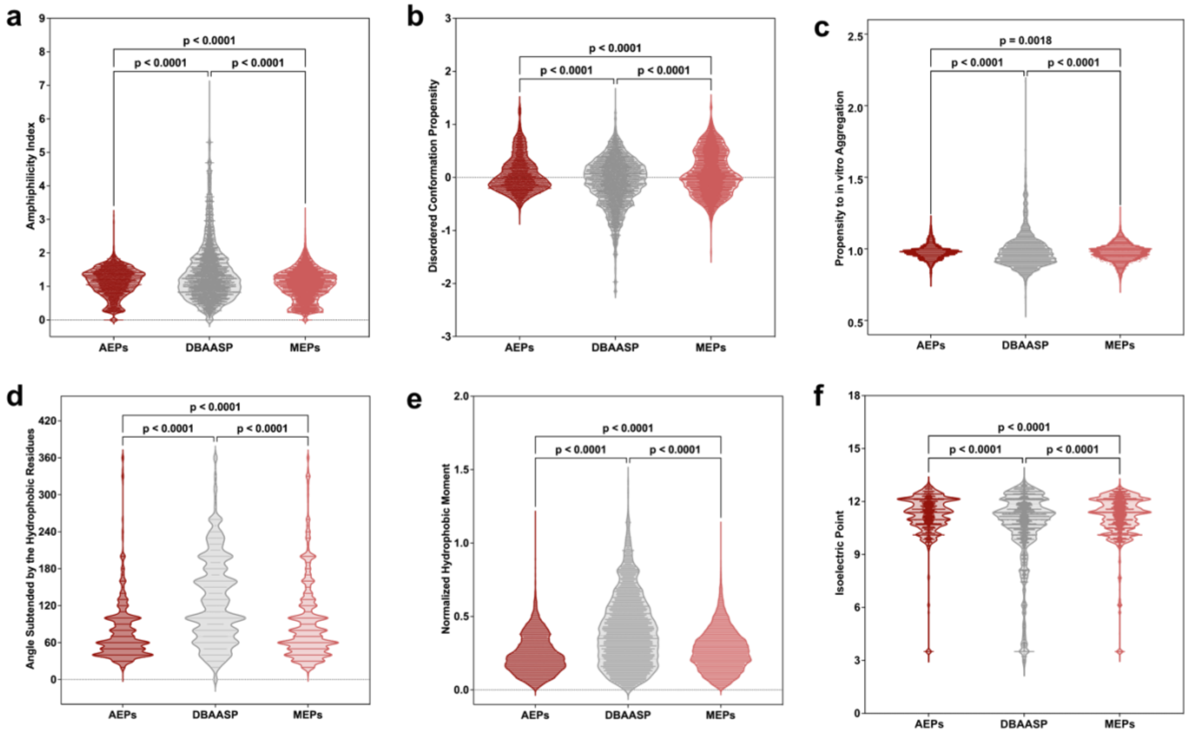


**Fig. S31. Physicochemical features of AEPs and MEPs identified by APEX in extinct organisms compared to AMPs from DBAASP. (a)** Amphiphilicity index and **(b)** disordered conformation propensity; both properties closely correlated with mechanism of action, *i.e.*, how the peptides interact with membrane lipids to exert antimicrobial activity. **(c)** Propensity to aggregate *in vitro* and **(d)** angle subtended by the hydrophobic residues; both properties are correlated with supramolecular arrangement of the molecules and toxicity. **(e)** Hydrophobic moment normalized by peptide length and **(f)** isoelectric point; both properties are also related to the amphipathicity of the molecules that influence directly on their interactions with bacterial membranes. Statistical significance was obtained using t-tests followed by Mann-Whitney test.

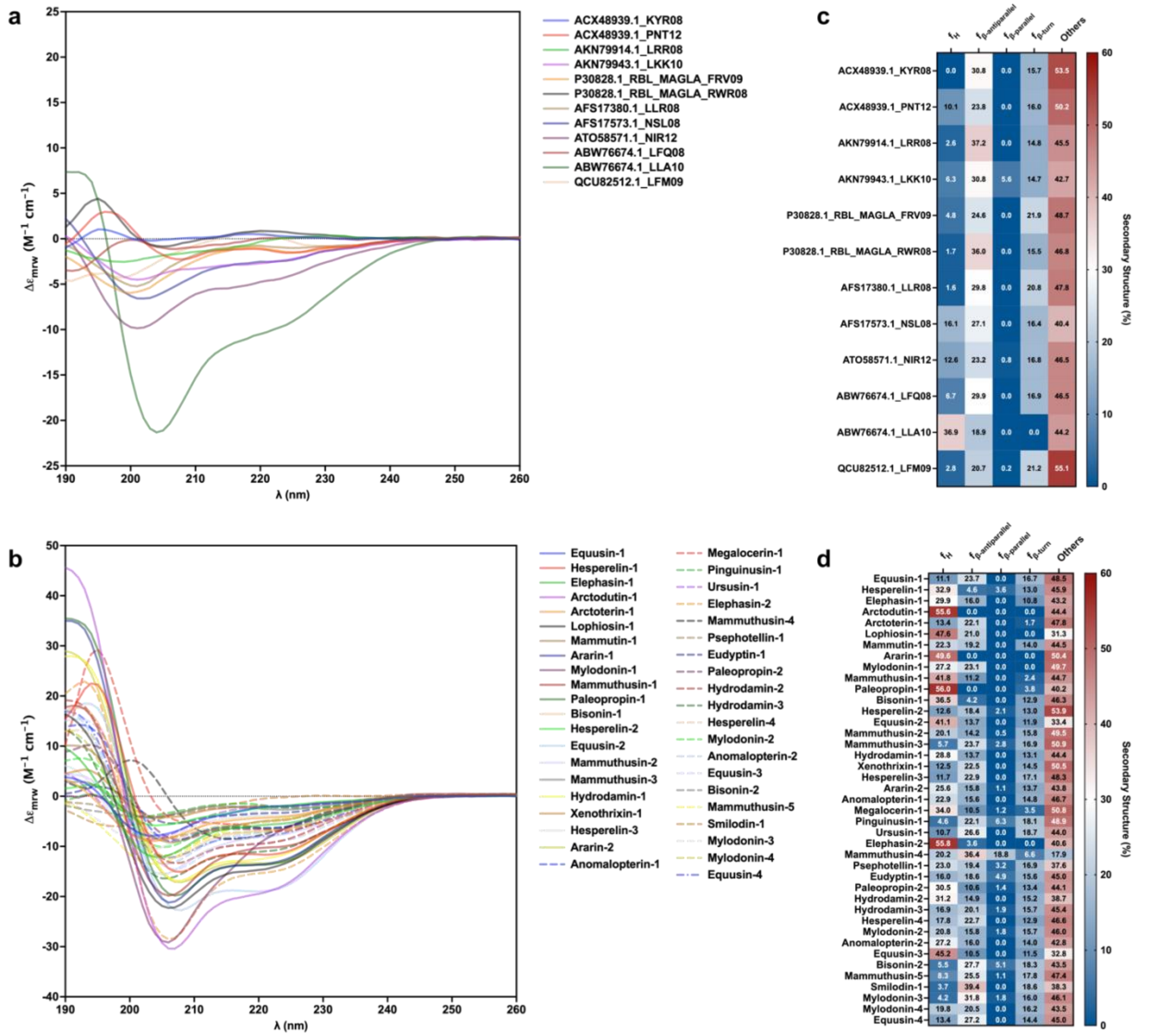


**Fig. S32. Secondary structure of active EPs predicted by the scoring function and APEX in helical inducer medium.** Circular dichroism experiments with encrypted peptides (EPs) from extinct organisms generated by **(a)** the scoring function based on length, hydrophobicity, and net charge, which predominantly identified unstructured peptides, and **(b)** the deep learning model, which identified EPs having a higher helical content than the EPs predicted by the scoring function. **(c-d)** Heat maps of the secondary structure basis components values obtained using the algorithm BeStSel[2] for EPs identified by the scoring function **(c)** or APEX **(d)**. Assays were performed in a J-1500 (Jasco circular dichroism spectrophotometer), and the circular dichroism spectra were recorded after three accumulations at 25 °C, using a 1 mm path length quartz cell, between 260 and 190 nm at 50 nm $min^{-1}$, with a bandwidth of 0.5 nm.

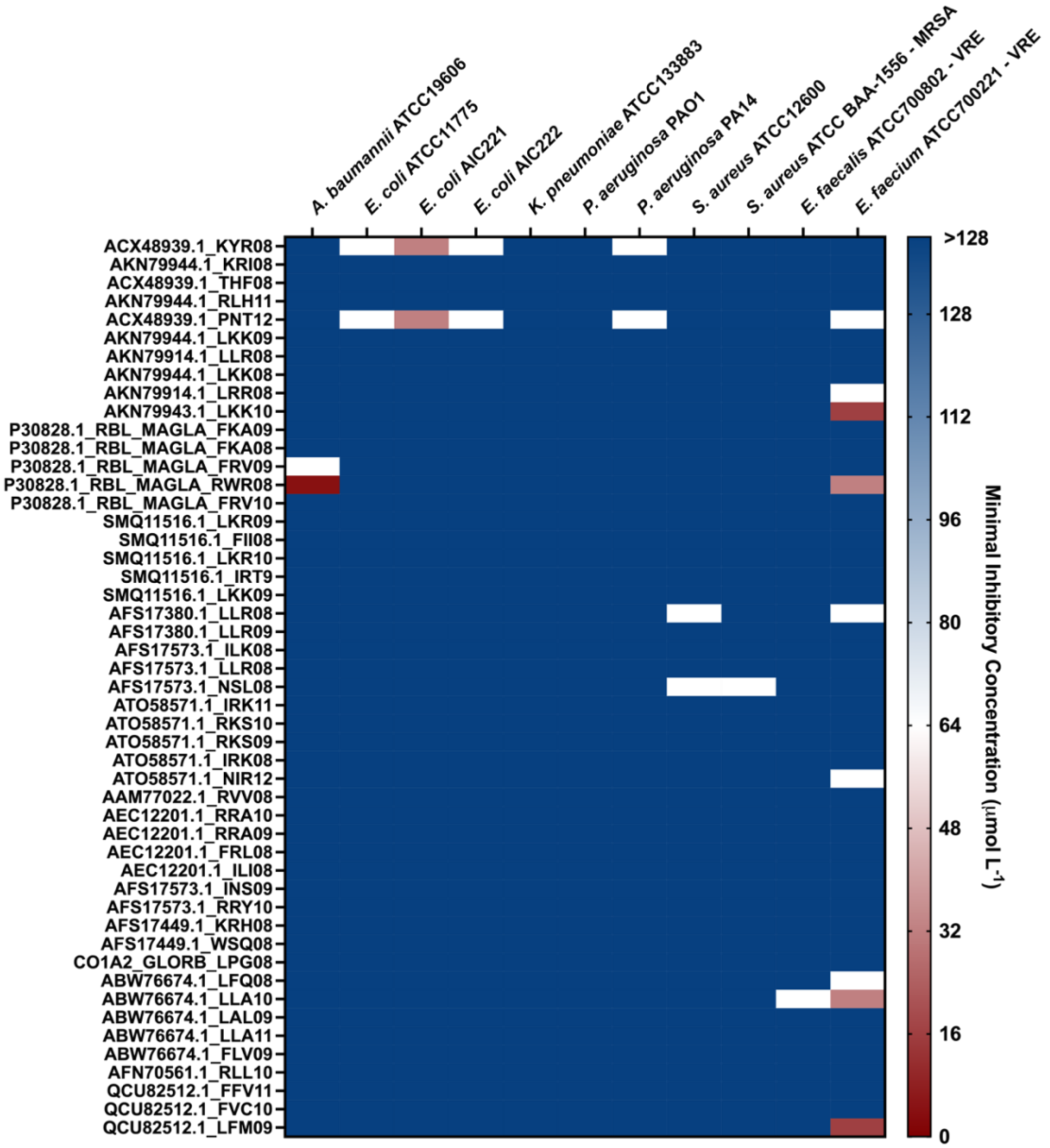


**Fig. S33. Antimicrobial activity of EPs from extinct organisms predicted by the scoring function.** Heat map of the antimicrobial activities (µmol $L^{-1}$) of the encrypted peptides (EPs) from extinct organisms predicted by the scoring function (Torres et al.[1]) based on length, hydrophobicity, and net charge, for 11 pathogens, including four strains resistant to conventional antibiotics. All the EPs identified by the scoring function were modern EPs (MEPs), *i.e.*, EPs identified in extinct organisms but also present in modern proteins. Briefly, $10^6$ bacterial cells and serially diluted MEPs (0–128 µmol $L^{-1}$) were incubated at 37 °C. One day post-treatment, the optical density at 600 nm was measured in a microplate reader to determine whether the EPs from extinct organisms inhibited bacterial growth *in vitro*. . Assays were performed in three independent replicates, and MIC values in the heat map are the arithmetic mean of the replicates in each condition.

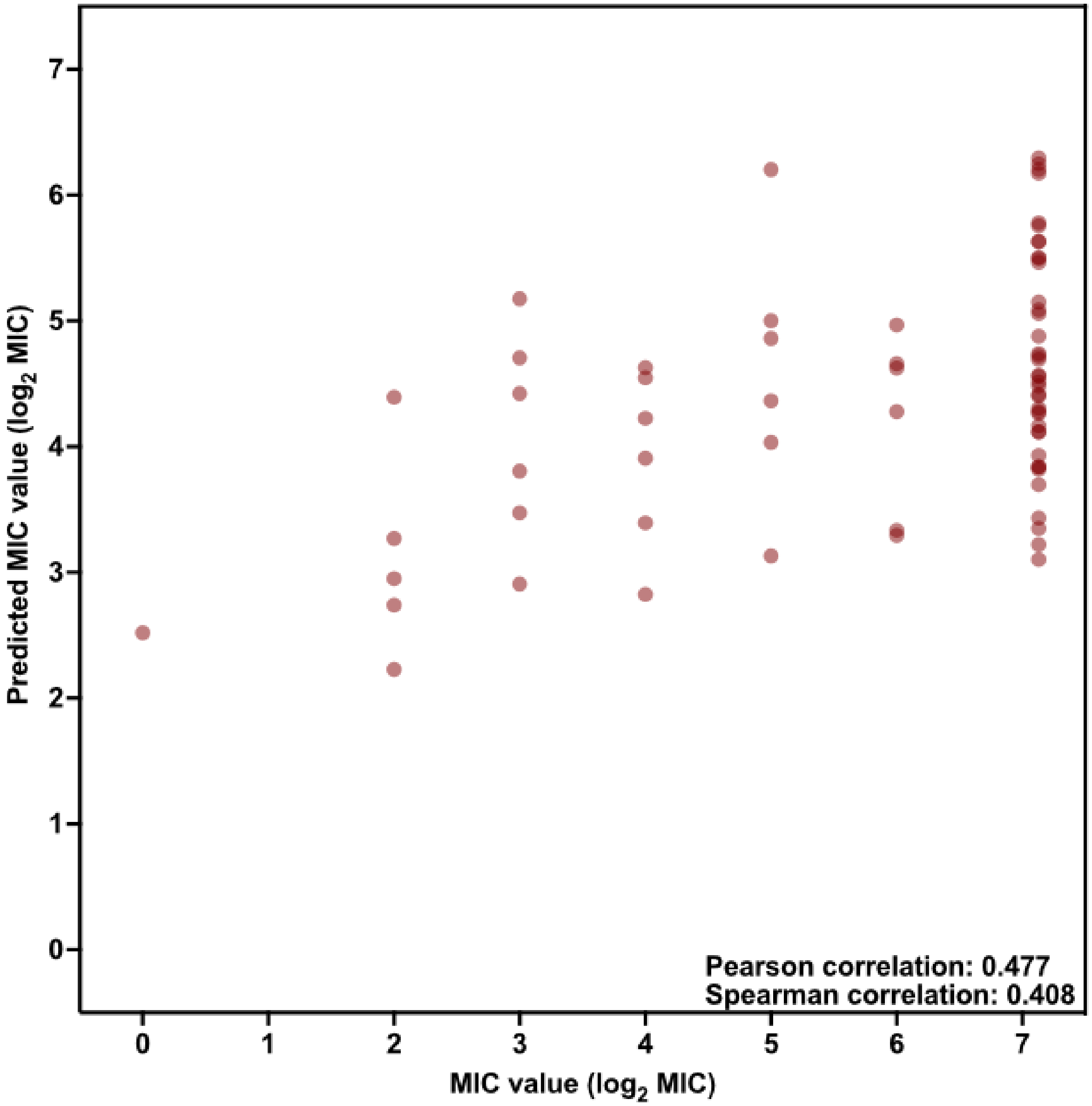


**Fig. S34. Predicted vs. experimental MICs of the EPs identified by APEX for *A. baumannii* ATCC 19606**. Each peptide on the scatter plot is represented by a red circle. Pearson and Spearman correlations values are shown as an inset.

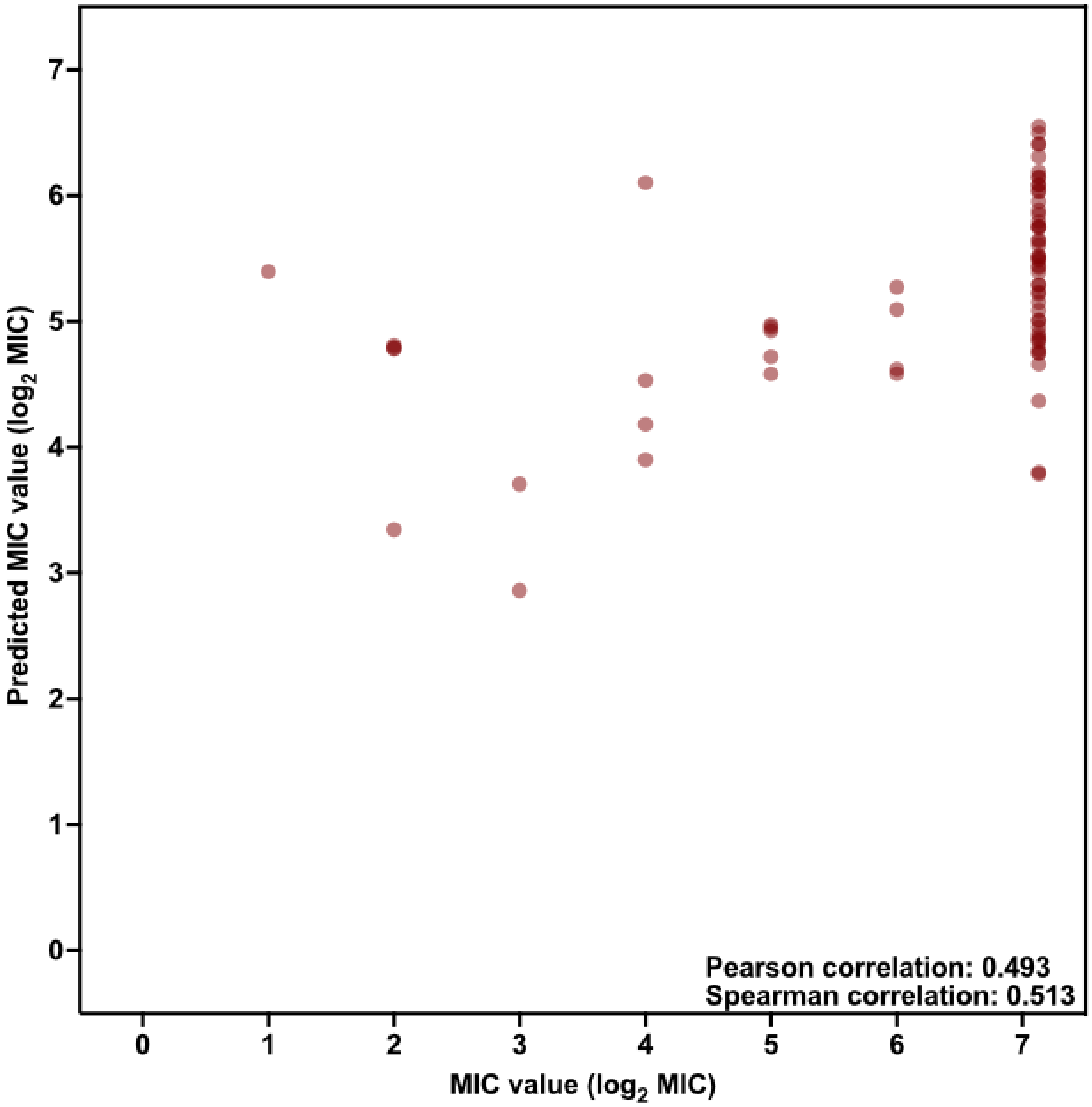


**Fig. S35. Predicted vs. experimental MICs of the EPs identified by APEX for *E. coli* AIC221**. Each peptide on the scatter plot is represented by a red circle. Pearson and Spearman correlations values are shown as an inset.

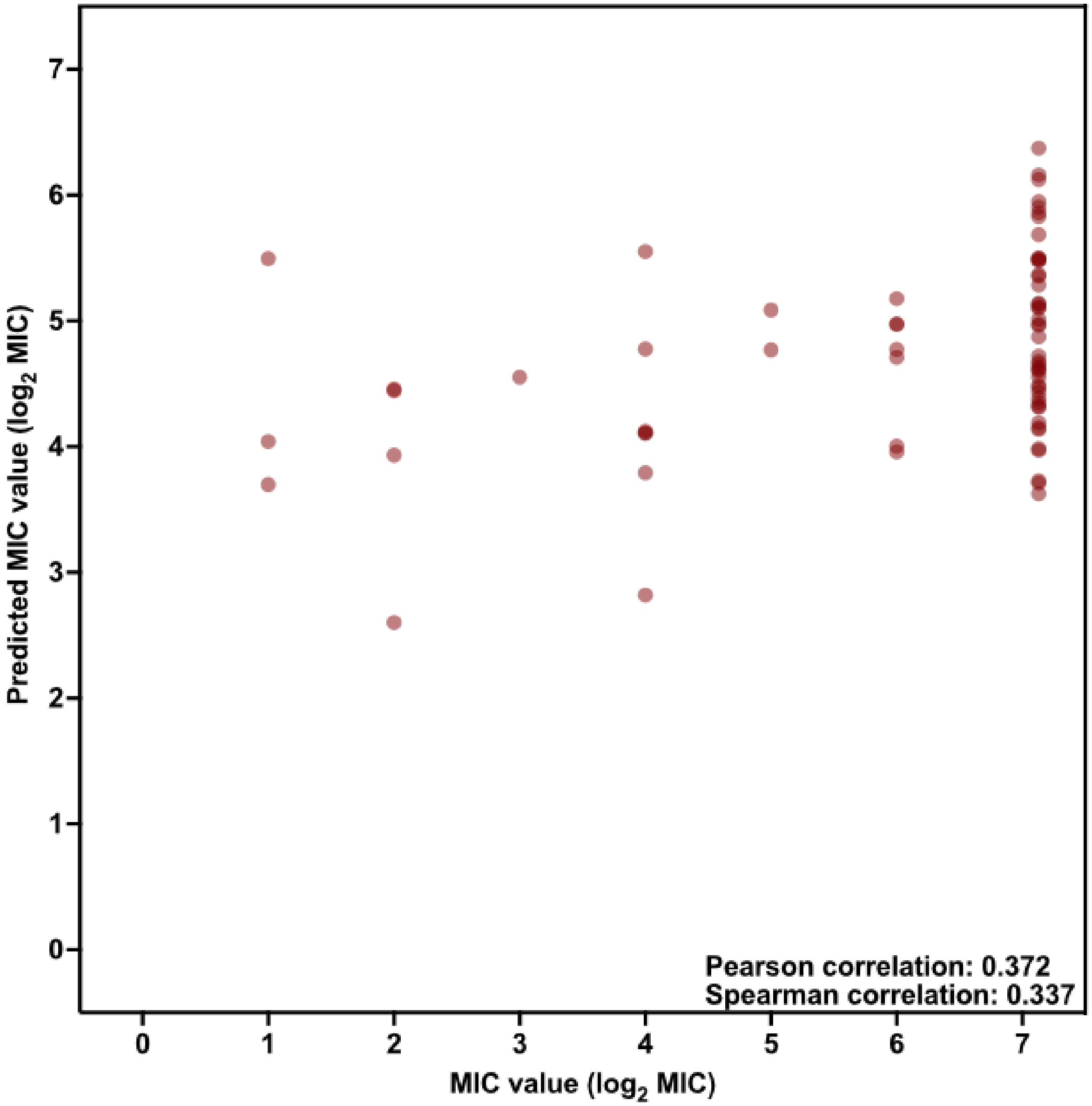


**Fig. S36. Predicted *vs* experimental MICs of the EPs identified by APEX for *E. coli* AIC222.** Each peptide on the scatter plot is represented by a red circle. Pearson and Spearman correlations values are shown as an inset.

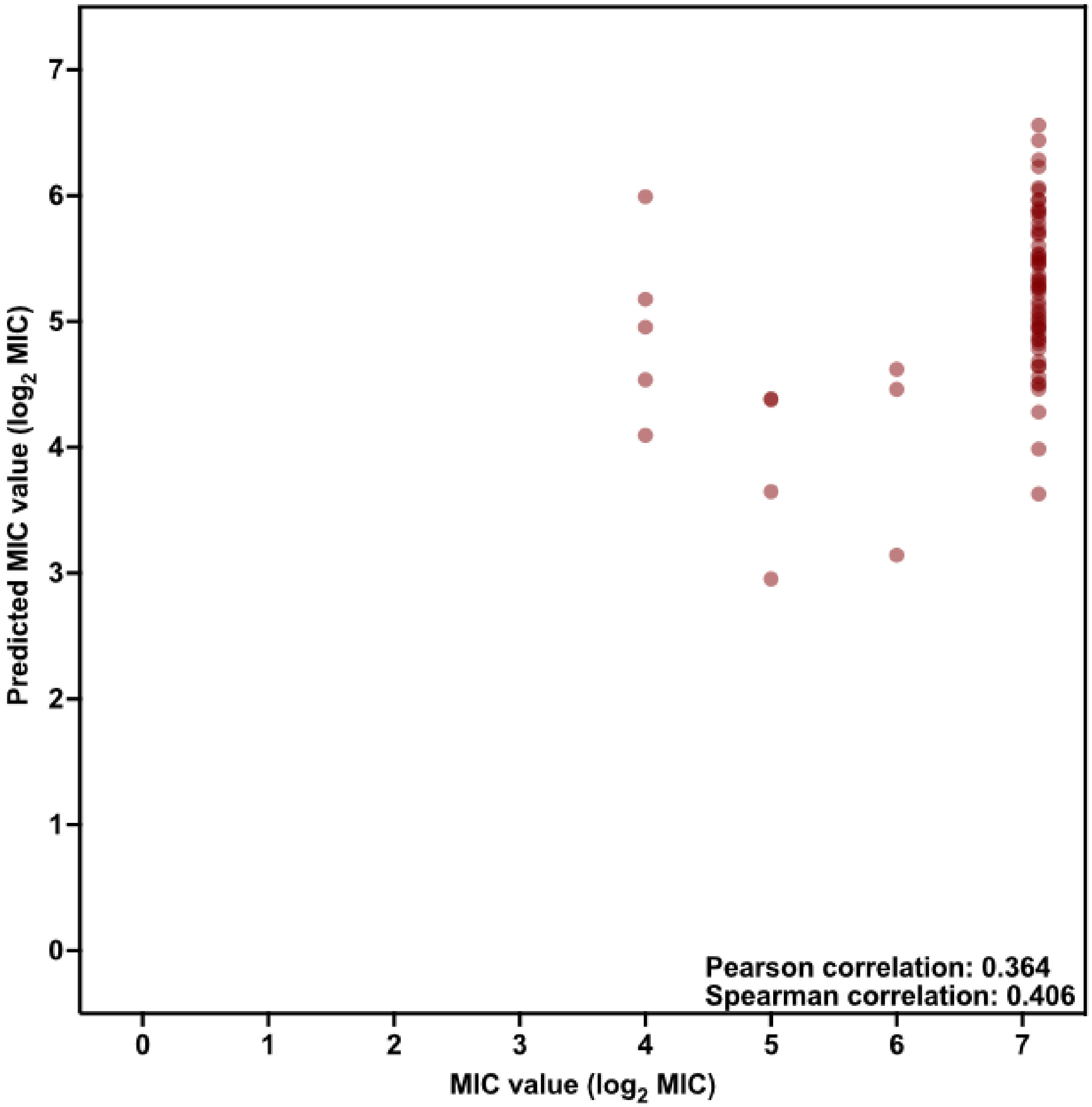


**Fig. S37. Predicted vs. experimental MICs of the EPs identified by APEX for *E. coli* ATCC 11775.** Each peptide on the scatter plot is represented by a red circle. Pearson and Spearman correlations values are shown as an inset.

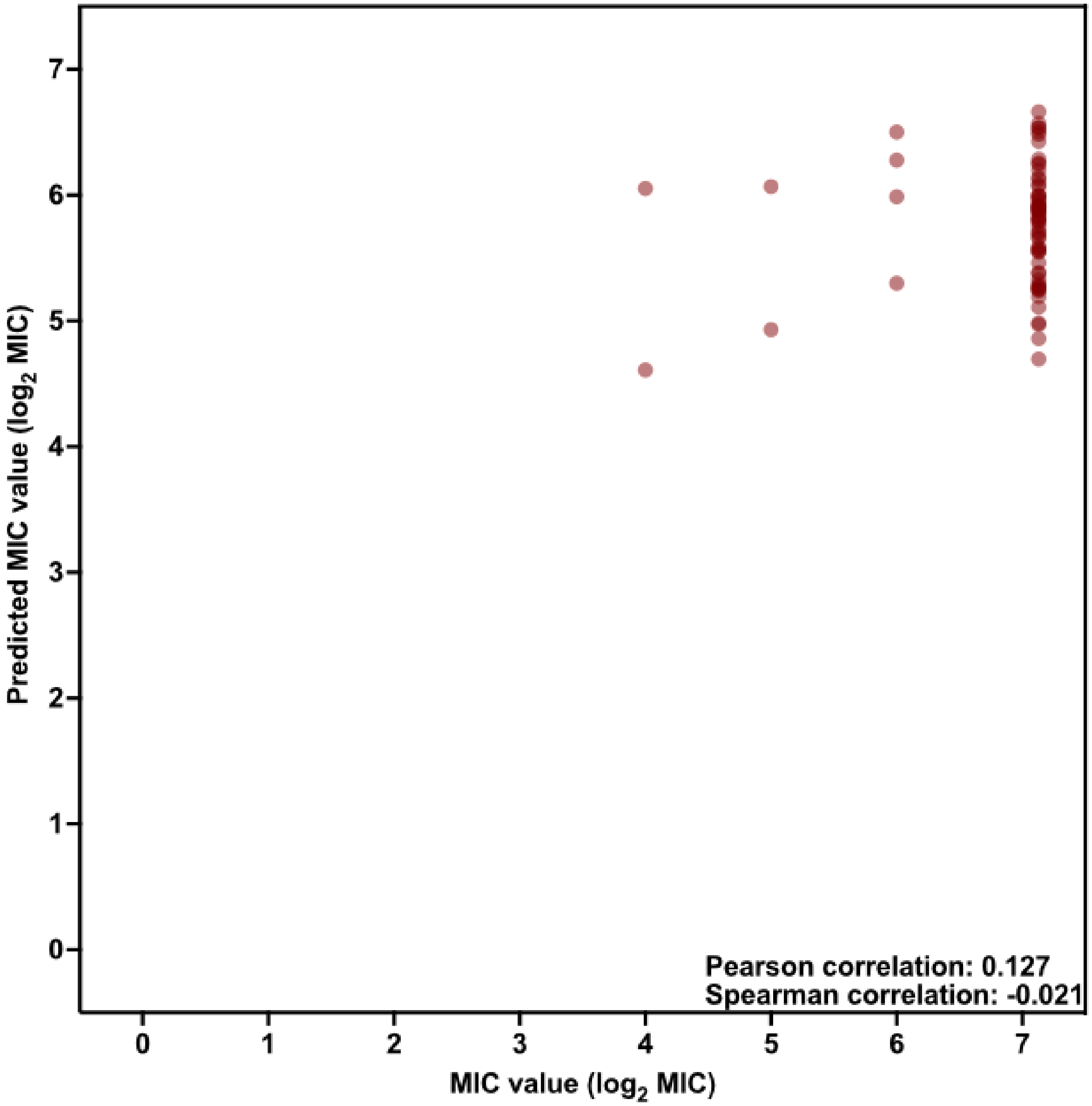


**Fig. S38. Predicted vs. experimental MICs of the EPs identified by APEX for *K. pneumoniae* ATCC 13883.** Each peptide on the scatter plot is represented by a red circle. Pearson and spearman correlations values are shown as an inset.

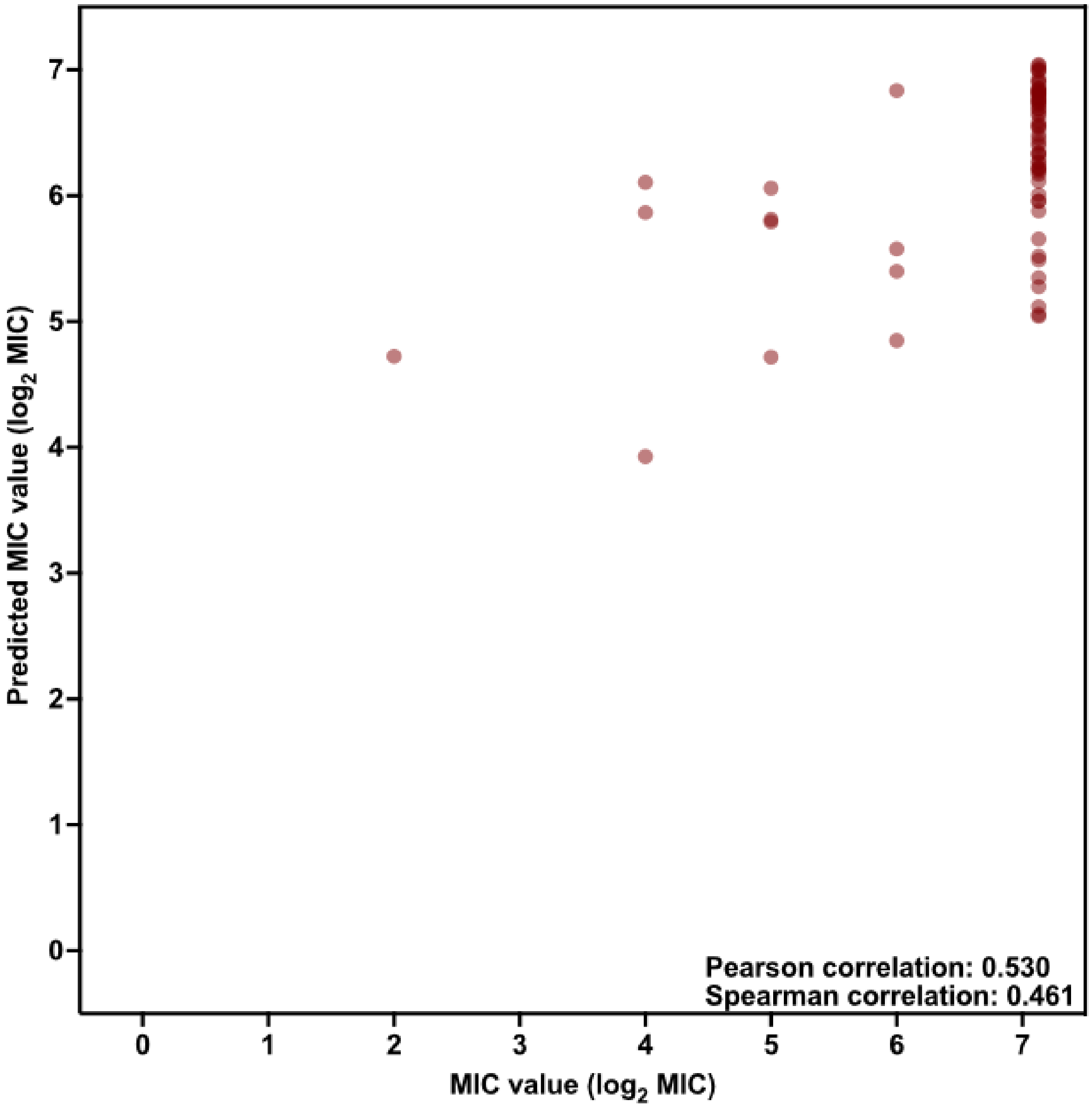


**Fig. S39. Predicted vs. experimental MICs of the EPs identified by APEX for *P. aeruginosa* PA14.** Each peptide on the scatter plot is represented by a red circle. Pearson and spearman correlations values are shown as an inset.

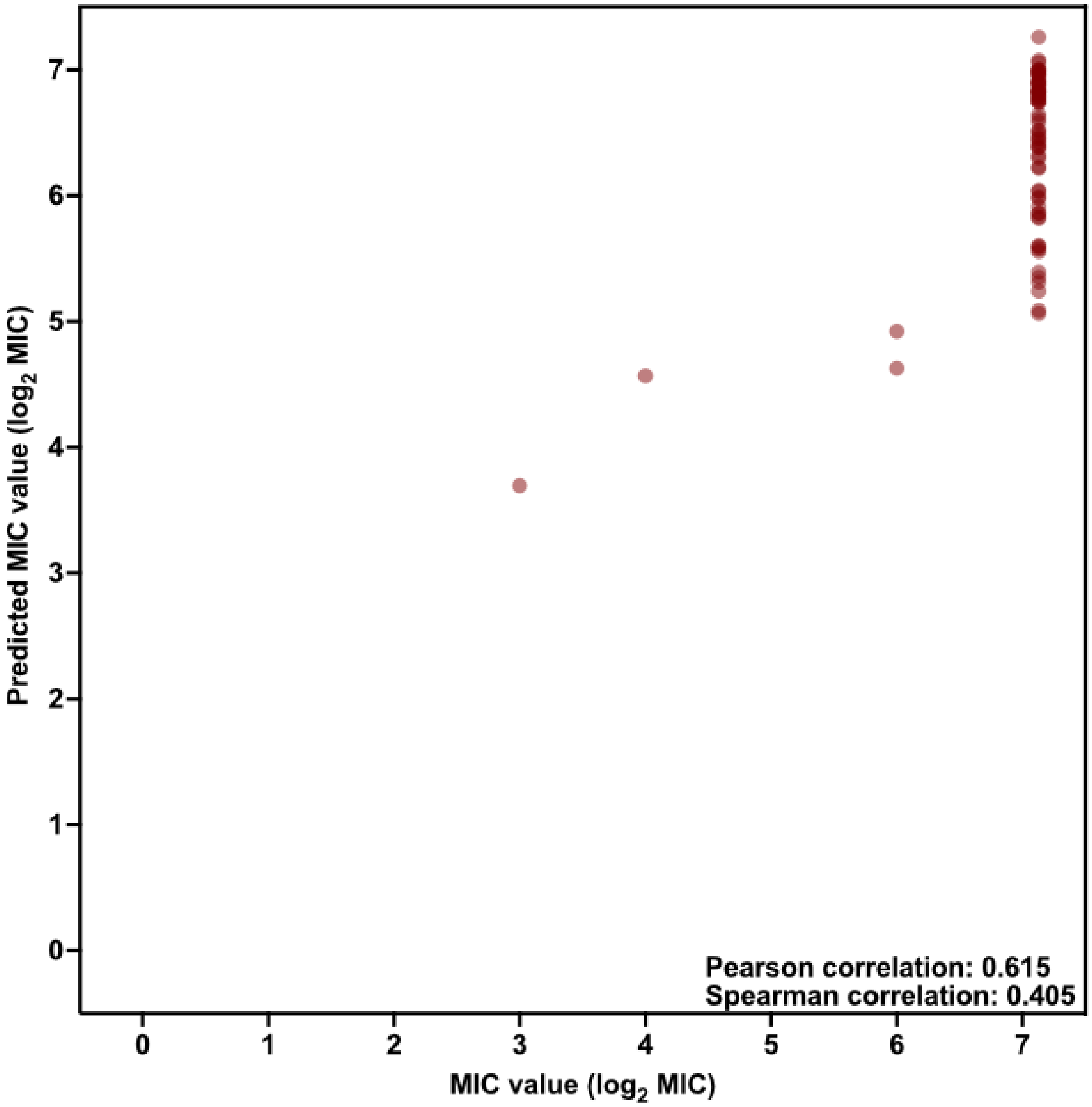


**Fig. S40. Predicted vs. experimental MICs of the EPs identified by APEX for *P. aeruginosa* PAO1.** Each peptide on the scatter plot is represented by a red circle. Pearson and spearman correlations values are shown as an inset.

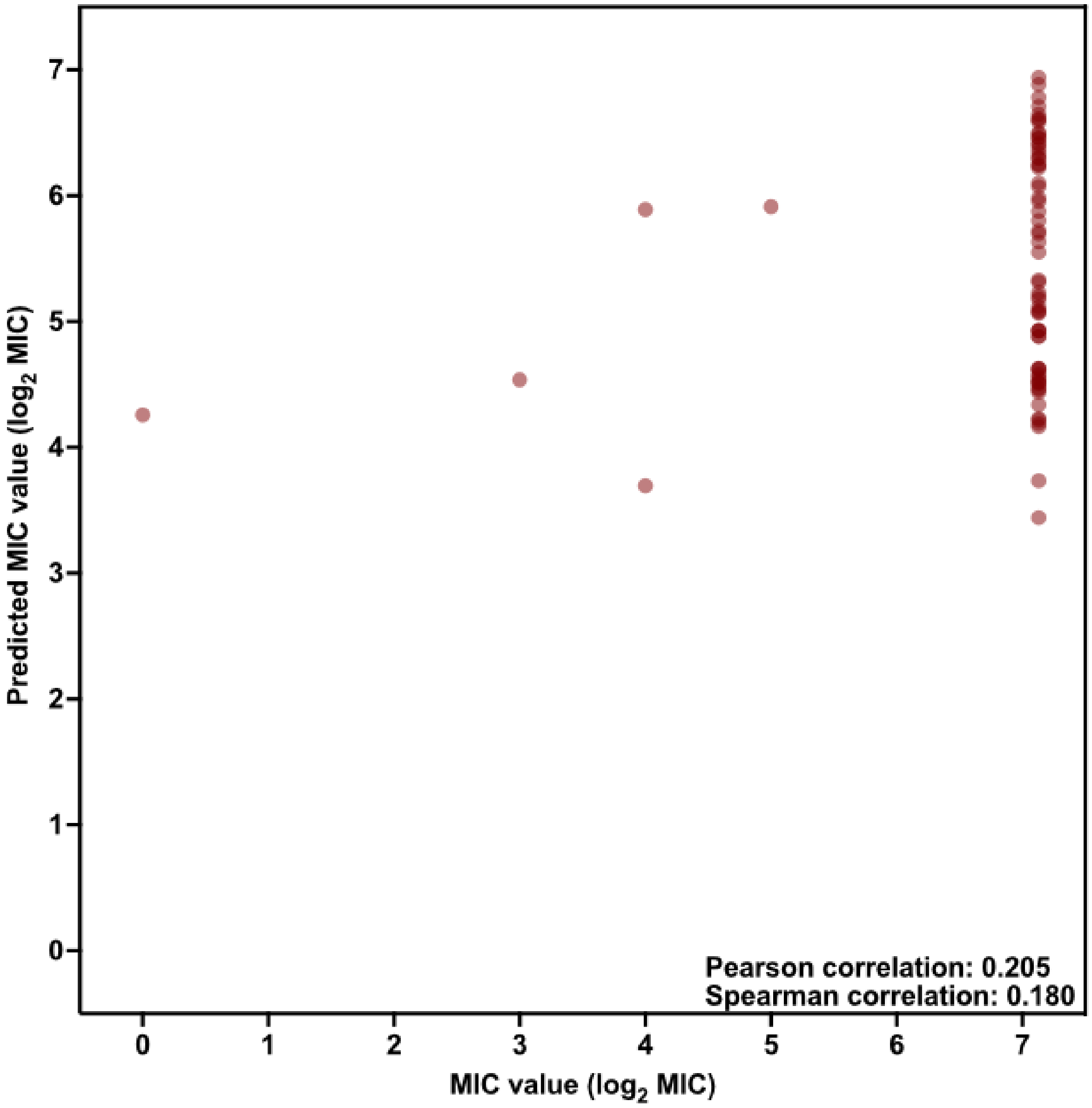


**Fig. S41. Predicted vs. experimental MICs of the EPs identified by APEX for methicillin-resistant *S. aureus* ATCC BAA-1556.** Each peptide on the scatter plot is represented by a red circle. Pearson and spearman correlations values are shown as an inset.

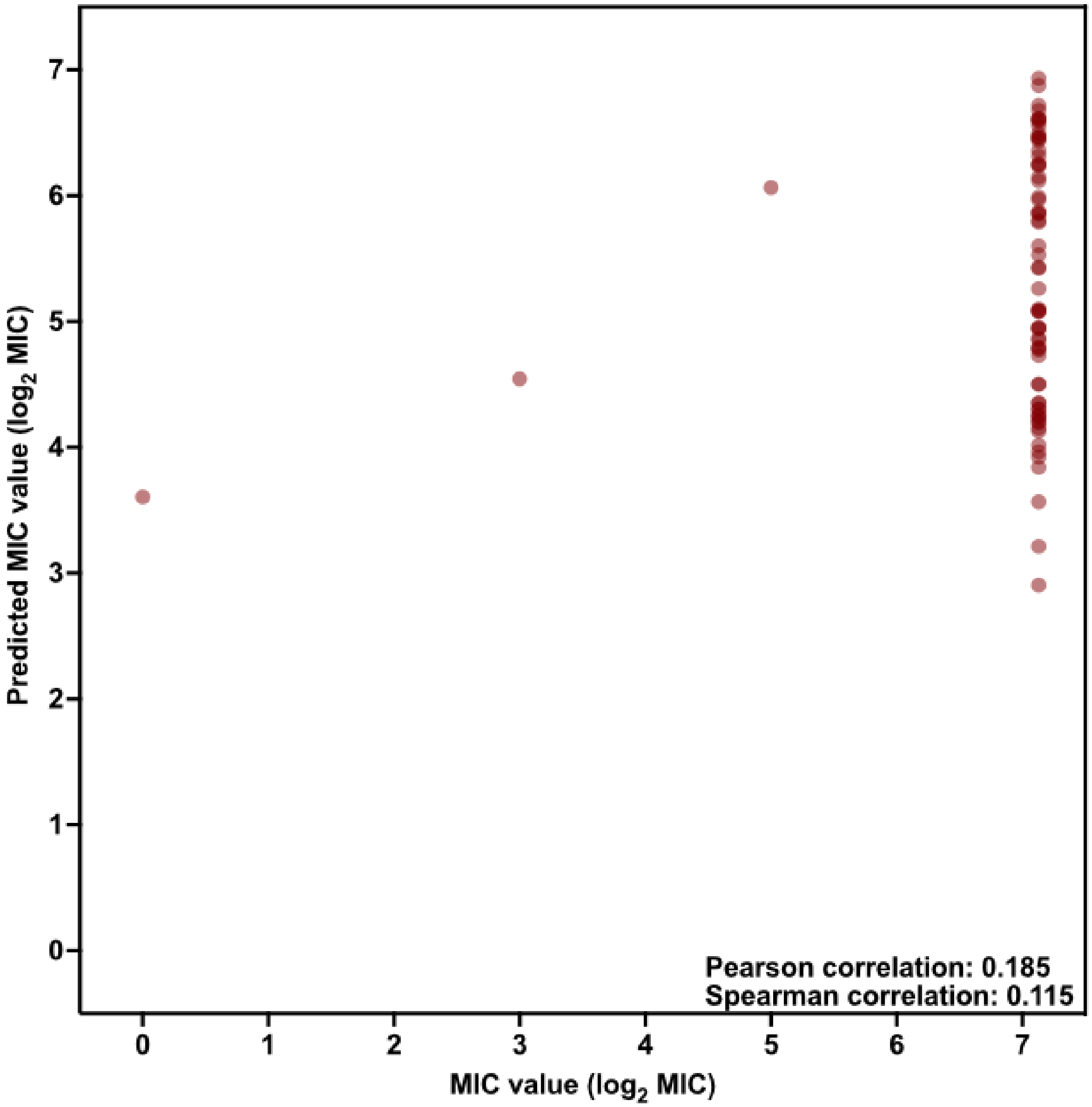


**Fig. S42. Predicted vs. experimental MICs of the EPs identified by APEX for *S. aureus* ATCC 12600.** Each peptide on the scatter plot is represented by a red circle. Pearson and spearman correlations values are shown as an inset.

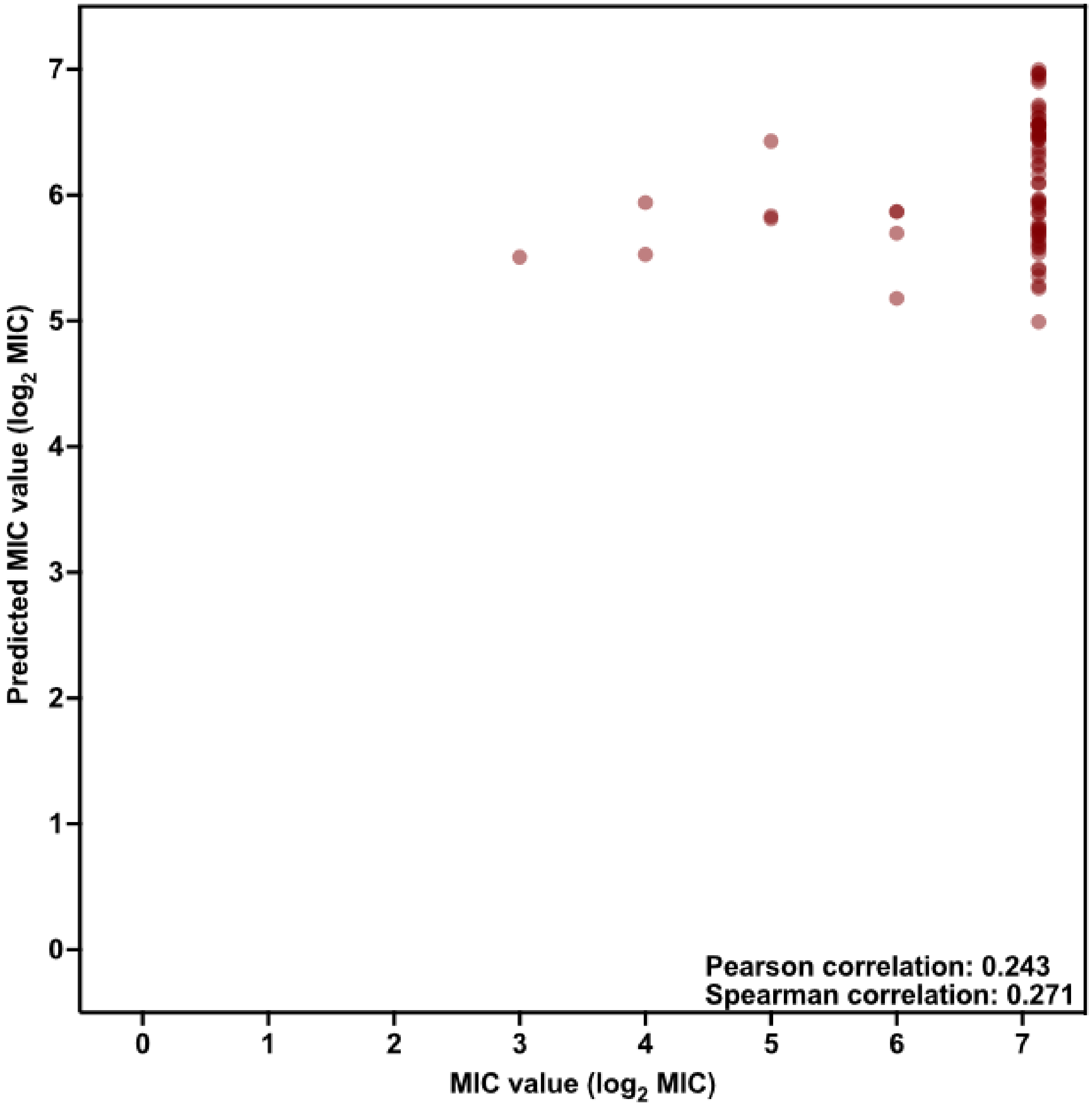


**Fig. S43. Predicted vs. experimental MICs of the EPs identified by APEX for vancomycin-resistant *E. faecalis* ATCC 700802.** Each peptide on the scatter plot is represented by a red circle. Pearson and spearman correlations values are shown as an inset.

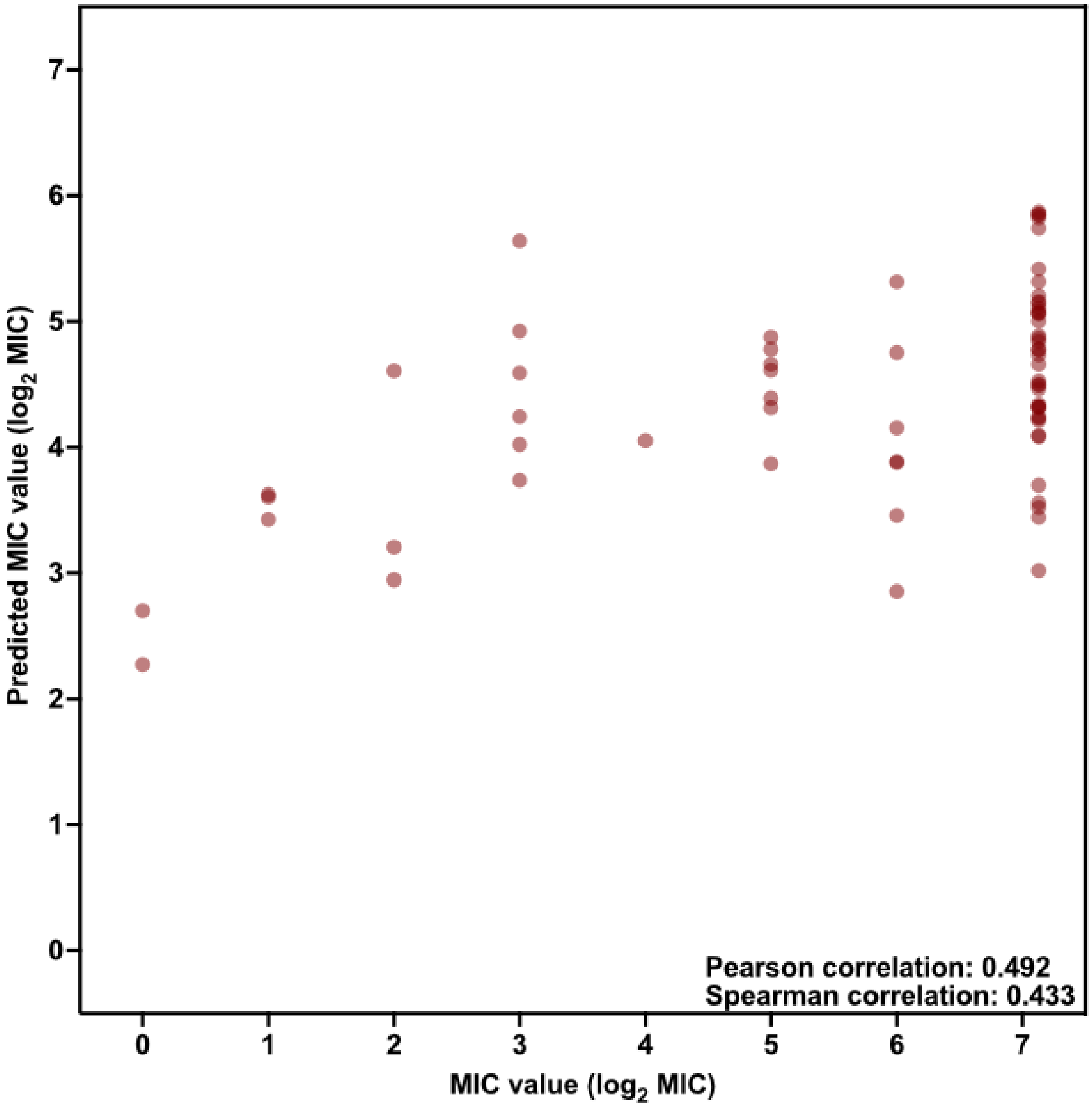


**Fig. S44. Predicted vs. experimental MICs of the EPs identified by APEX for vancomycin-resistant *E. faecium* ATCC 700221.** Each peptide on the scatter plot is represented by a red circle. Pearson and spearman correlations values are shown as an inset.

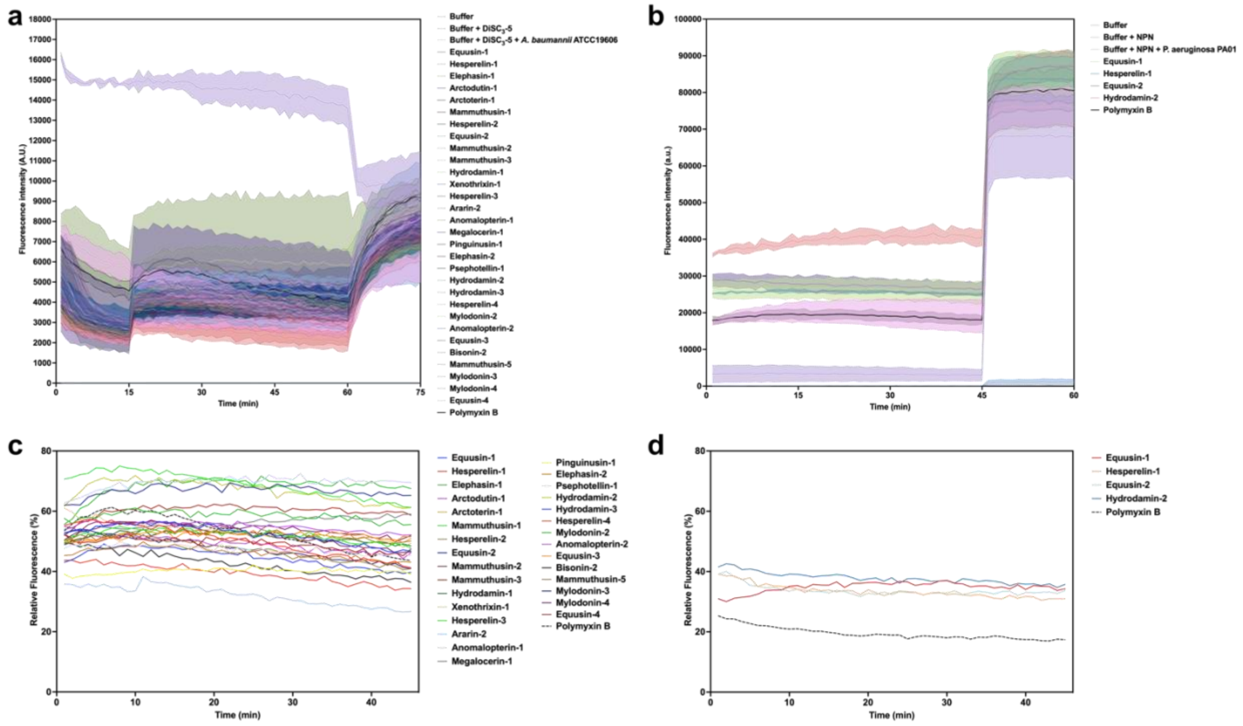


**Fig. S45.** **Cytoplasmic membrane depolarization of *A. baumannii* and *P. aeruginosa* triggered by AEPs and MEPs identified by APEX. (a-b)** Depolarization assays with the hydrophobic probe 3,3′-dipropylthiadicarbocyanine iodide [DiSC3-(5)] for all the encrypted peptides (EPs) from extinct organisms that were active against *A. baumannii* ATCC 19606 **(a)** and *P. aeruginosa* PAO1 **(b)**. Polymyxin B was used as positive control, and buffer, buffer with DiSC3-(5), and buffer with DiSC3-(5) and bacteria were used as baseline for fluorescence. The panels show the raw fluorescence intensity data obtained in the experiments. **(c-d)** Relative fluorescence values of the depolarization effect of EPs from extinct organisms compared to the untreated control on *A. baumannii* ATCC 19606 **(c)** and *P. aeruginosa* PAO1 **(d)** cytoplasmic membranes.

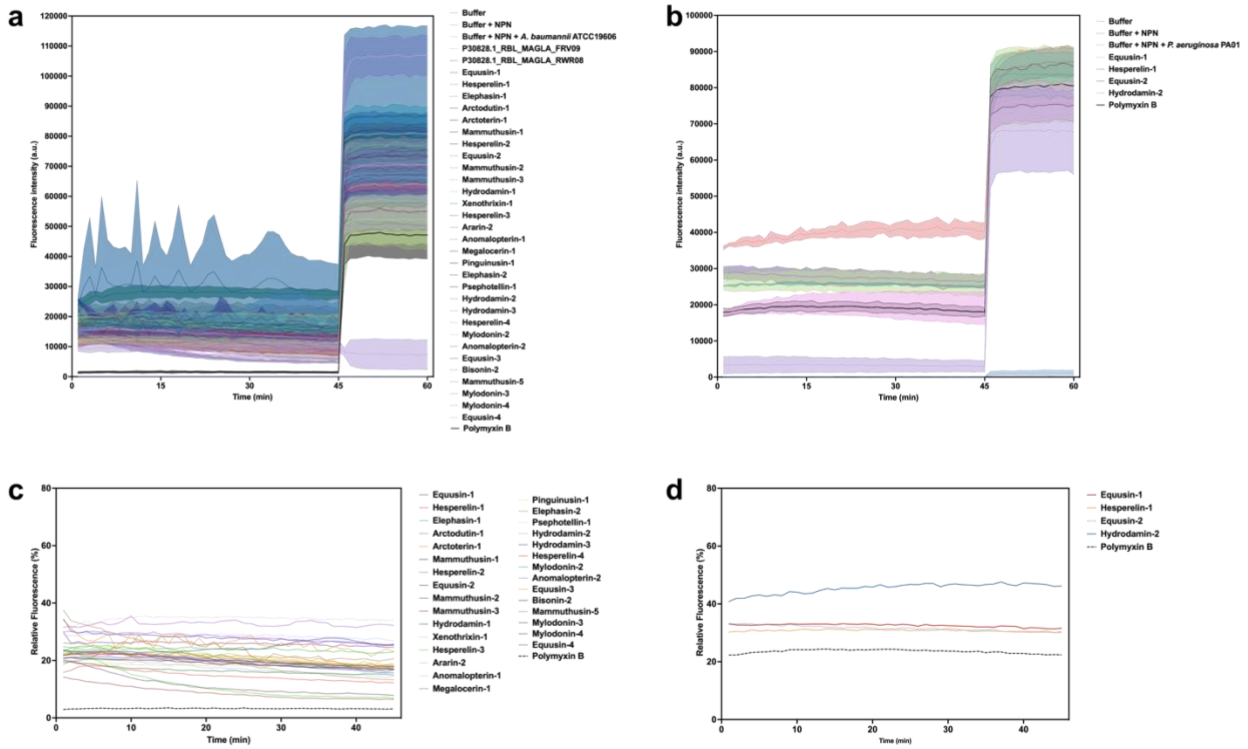


**Fig. S46. Outer membrane permeabilization of *A. baumannii* and *P. aeruginosa* cell membranes caused by encrypted peptides from extinct organisms.** The probe 1-(N-phenylamino)naphthalene (NPN) was used to detect permeabilization of the outer membrane. **(a-b)** Permeabilization by encrypted peptides (EPs) from extinct organisms active against each one of the strains: on *A. baumannii* ATCC 19606 **(a)** and *P. aeruginosa* PAO1 **(b)**. Polymyxin B was used as positive control. Buffer was used as the baseline for fluorescence, and buffer with NPN, and buffer with NPN and bacteria were used as baseline for fluorescence. The panels show the raw fluorescence intensity data obtained in the experiments. **(c-d)** Relative fluorescence values of the permeabilization effect of encrypted peptides from extinct organisms compared to the untreated control on *A. baumannii* ATCC 19606 **(c)** and *P. aeruginosa* PAO1 **(d)** outer membranes.

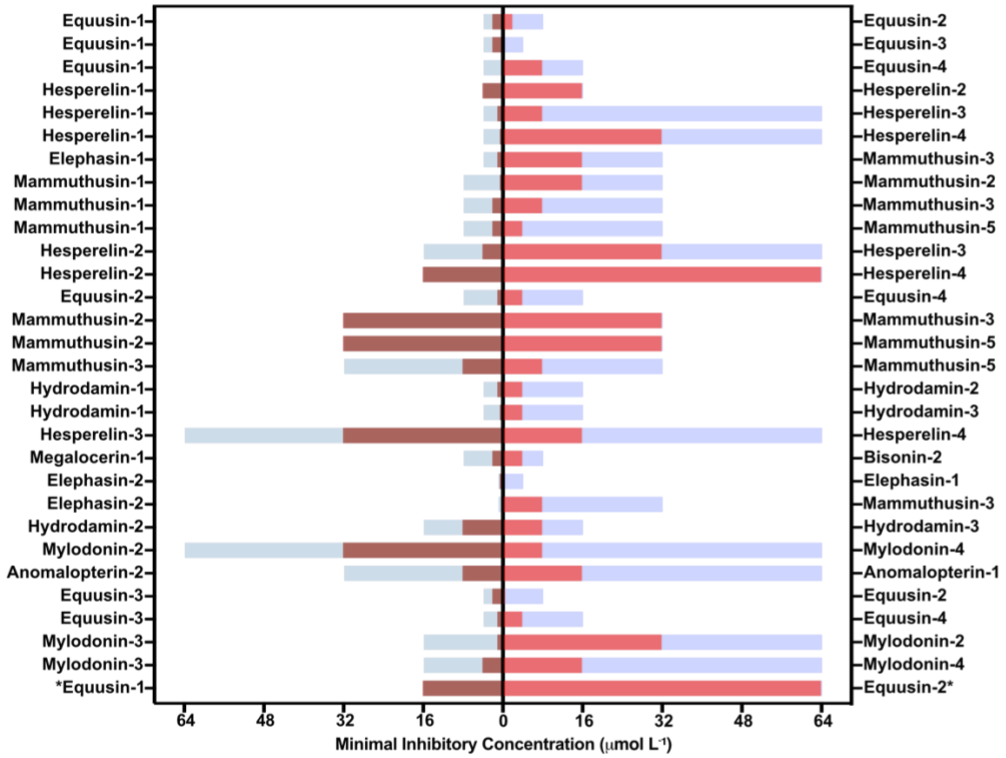


**Fig. S47. Synergy between AEPs and MEPs from extinct organisms.** Bar plot showing the synergistic interactions between EPs from the same extinct organism against *A. baumannii* and *P. aeruginosa* (*P. aeruginosa* result is indicated with an asterisk). Stacked bars represent the MICs in μmol $L^{-1}$ values in each condition. MICs of the individual peptides are shown in blue and light purple and when in combination, shown in brown and red. Each pair of peptides was placed in the same row for side-by-side comparison of MIC-fold change before and after they were tested in combination.

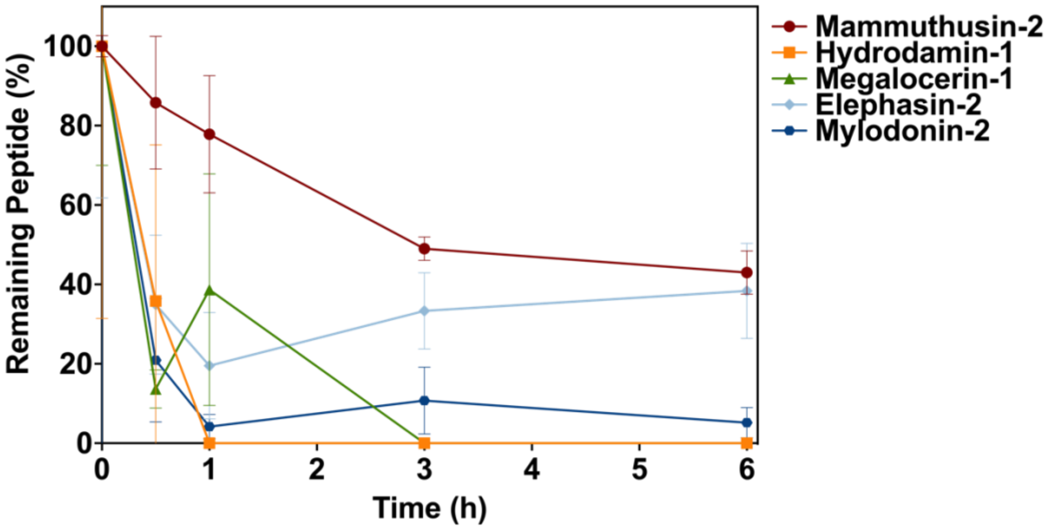


**Fig. S48. Resistance to proteolytic degradation.** The AEPs hydrodamin-1, elephasin-2, and mylodonin-2 and the MEPs mammuthusin-2 and megalocerin-1 were exposed for a total period of 6 h to human serum that contains several active proteases. Aliquots of the resulting solution were analyzed by liquid chromatography coupled to mass spectrometry. In summary, the AEP elephasin-2 and the MEP mammuthusin-2 exhibited the highest resistance to proteolytic degradation with ~40% peptide remaining after 6 h of exposure. All other AEPs and MEP tested degraded at varying degrees within the duration of the experiment. Experiments were performed in three independent replicates.

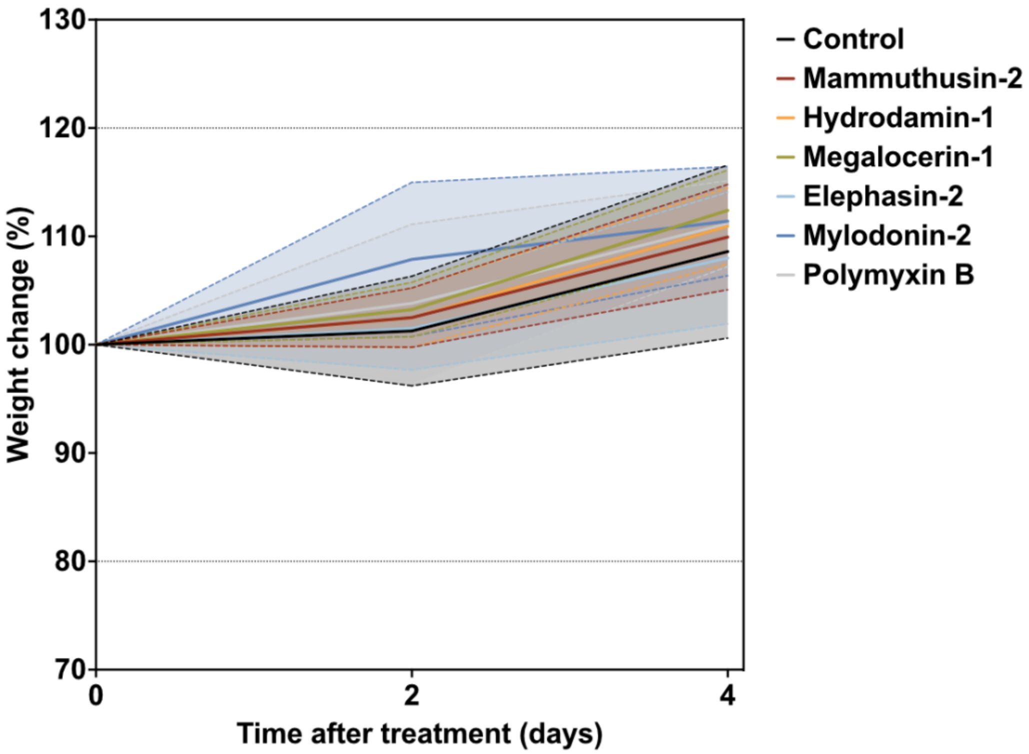


**Fig. S49. Weight change monitoring in skin abscess infection mouse model infected with *A. baumannii* cells.** Mouse weight was monitored throughout the duration of the skin abscess assay (4 days total) to rule out the potentially toxic effects of bacterial load and the encrypted peptides (EPs).

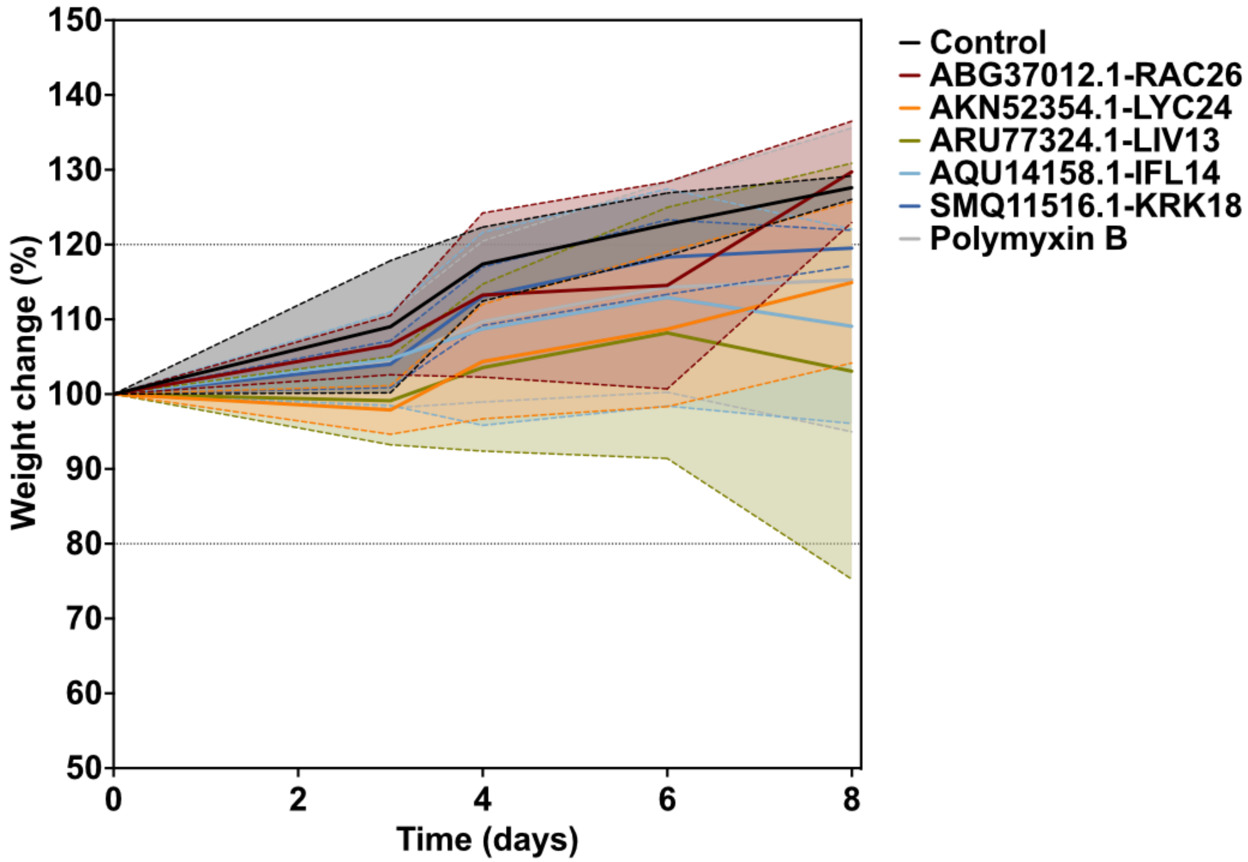


**Fig. S50. Weight change monitoring in thigh infection mouse model infected with *A. baumannii* cells.** Mouse weight was monitored throughout the duration of the thigh infection model (8 days total) to rule out the potentially toxic effects of bacterial load or the encrypted peptides (EPs).

### Supplementary Tables:

#### Table S1. Hyperparameter ranges explored for APEX.

| Hyperparameters | number of RNN layers | n | m | λ_l2 | λ_(multitask_constraint) | λ_BCE |
|---|---|---|---|---|---|---|
| Hyperparameter range | {1, 2, 3} | {128, 256} | {512, 1024, 2048} | {1e-5, 1e-6} | {0.1, 0.01, 0.001, 0.0} | {1.0, 0.1, 0.0} |

#### Table S2. Hyperparameter ranges searched for elastic net.

| Elastic Net hyperparameter | alpha |
|---|---|
| Hyperparameter range | {1.0, 0.5, 0.1, 0.05, 0.01, 0.005, 0.001, 0.0005, 0.0001} |

#### Table S3. Hyperparameter ranges searched for linear support vector machine.

| Linear support vector regression hyperparameter | C |
|---|---|
| Hyperparameter range | {1, 10, 20, 30, …, 1000} |

#### Table S4. Hyperparameter ranges searched for tree-based models.

| Random forest, gradient boosting decision tree and, extra-trees regressor hyperparameter hyperparameters | Num estimators | Max depth |
|---|---|---|
| Hyperparameter range | {128, 256, 512, 1024} | {8, 16, 32, 64} |

**Table S5. R-squared of various ML models on CV set.** We divided our in-house peptide dataset into a CV set for hyperparameter tuning and ML model selection, and an independent dataset to evaluate prediction performance. The table shows the average species-specific prediction performance of 5-fold CV in terms of R-squared for various ML models on the CV set. RF: random forest; GBDT: gradient boosting decision tree; ExtraTree: extra-tree regressor; ElasticNet: elastic net; LinearSVR: linear support vector regression.

| Strain | Ensemble APEX v1 | Single APEX | RF | GBDT | ExtraTree | ElasticNet | LinearSVR |
|---|---|---|---|---|---|---|---|
| *E. coli* ATCC11775 | 0.588229 | 0.519899 | 0.494097 | 0.446439 | 0.434624 | 0.417793 | 0.128045 |
| *P. aeruginosa* PAO1 | 0.520804 | 0.458351 | 0.490921 | 0.401458 | 0.430824 | 0.41167 | -0.07728 |
| *P. aeruginosa* PA14 | 0.461245 | 0.415884 | 0.402854 | 0.327131 | 0.38079 | 0.360004 | 0.02522 |
| *S. aureus* ATCC12600 | 0.519007 | 0.443486 | 0.388353 | 0.348654 | 0.267847 | 0.271189 | 0.017144 |
| *E. coli* AIC221 | 0.574544 | 0.527619 | 0.46737 | 0.471732 | 0.428688 | 0.351852 | 0.142403 |
| *E. coli* AIC222 | 0.638583 | 0.578155 | 0.50825 | 0.497999 | 0.445026 | 0.408777 | 0.159808 |
| *K. pneumoniae* ATCC13883 | 0.215734 | 0.078356 | 0.093001 | 0.025148 | -0.17556 | 0.017496 | -0.36587 |
| *A. baumannii* ATCC19606 | 0.675531 | 0.597275 | 0.577164 | 0.581414 | 0.559864 | 0.420903 | 0.281722 |
| Methicillin-resistant *S. aureus* ATCC BAA-1556 | 0.539041 | 0.442817 | 0.346027 | 0.355517 | 0.15017 | 0.237193 | -0.2676 |
| Vancomycin-resistant *E. faecalis* ATCC700802 | -0.14859 | -0.46429 | -0.03412 | -0.56905 | -0.05681 | -0.01311 | -0.09197 |
| Vancomycin-resistant *E. faecium* ATCC700221 | 0.615554 | 0.460972 | 0.422509 | 0.347594 | 0.345839 | 0.441332 | 0.385905 |
| **Average** | **0.472699** | **0.368956** | **0.377857** | **0.294003** | **0.291936** | **0.302282** | **0.030685** |

**Table S6. Pearson correlation of various ML models on CV set.** We divided our in-house peptide dataset into a CV set for hyperparameter tuning and ML model selection, and an independent dataset to evaluate prediction performance. The table shows the average species-specific prediction performance of 5-fold CV in terms of Pearson correlation for various ML models on the CV set. RF: random forest; GBDT: gradient boosting decision tree; ExtraTree: extra-tree regressor; ElasticNet: elastic net; LinearSVR: linear support vector regression.

| **Strain** | **Ensemble APEX v1** | **Single APEX** | **RF** | **GBDT** | **ExtraTree** | **ElasticNet** | **LinearSVR** |
|---|---|---|---|---|---|---|---|
| *E. coli* ATCC11775 | 0.767053 | 0.722816 | 0.703998 | 0.691083 | 0.661328 | 0.646422 | 0.556482 |
| *P. aeruginosa* PAO1 | 0.72198 | 0.683082 | 0.701559 | 0.673932 | 0.656995 | 0.644568 | 0.450686 |
| *P. aeruginosa* PA14 | 0.679904 | 0.654815 | 0.635966 | 0.598879 | 0.619795 | 0.606475 | 0.449355 |
| *S. aureus* ATCC12600 | 0.720695 | 0.672914 | 0.623975 | 0.609784 | 0.548119 | 0.523365 | 0.469692 |
| *E. coli* AIC221 | 0.758512 | 0.728434 | 0.688737 | 0.6902 | 0.65761 | 0.594226 | 0.490013 |
| *E. coli* AIC222 | 0.799682 | 0.761297 | 0.715085 | 0.710545 | 0.667721 | 0.640531 | 0.52739 |
| *K. pneumoniae* ATCC13883 | 0.477322 | 0.401615 | 0.319796 | 0.332036 | 0.155075 | 0.228322 | 0.250313 |
| *A. baumannii* ATCC19606 | 0.822192 | 0.774607 | 0.761093 | 0.765975 | 0.749168 | 0.664209 | 0.58983 |
| Methicillin-resistant *S. aureus* ATCC BAA-1556 | 0.734576 | 0.680613 | 0.589402 | 0.620643 | 0.500931 | 0.497521 | 0.424084 |
| Vancomycin-resistant *E. faecalis* ATCC700802 | 0.096459 | 0.056318 | 0.029826 | 0.026884 | 0.029424 | -0.15356 | 0.048211 |
| Vancomycin-resistant *E. faecium* ATCC700221 | 0.784896 | 0.697111 | 0.655262 | 0.617407 | 0.600566 | 0.674514 | 0.622007 |
| **Average** | **0.669388** | **0.621238** | **0.584063** | **0.576124** | **0.531521** | **0.506053** | **0.44346** |

**Table S7. Spearman correlation of various ML models on CV set.** We divided our in-house peptide dataset into a CV set for hyperparameter tuning and ML model selection, and an independent dataset to evaluate prediction performance. The table shows the average species-specific prediction performance of 5-fold CV in terms of Spearman correlation for various ML models on the CV set. RF: random forest; GBDT: gradient boosting decision tree; ExtraTree: extra-tree regressor; ElasticNet: elastic net; LinearSVR: linear support vector regression.

| Strain | Ensemble APEX v1 | Single APEX | RF | GBDT | ExtraTree | ElasticNet | LinearSVR |
|---|---|---|---|---|---|---|---|
| *E. coli* ATCC11775 | 0.686485 | 0.66331 | 0.622101 | 0.603803 | 0.571779 | 0.572859 | 0.446956 |
| *P. aeruginosa* PAO1 | 0.636473 | 0.611234 | 0.606478 | 0.602546 | 0.579222 | 0.54594 | 0.368697 |
| *P. aeruginosa* PA14 | 0.658294 | 0.625266 | 0.611538 | 0.592355 | 0.596603 | 0.569181 | 0.405066 |
| *S. aureus* ATCC12600 | 0.546967 | 0.484758 | 0.433656 | 0.468351 | 0.297889 | 0.408671 | 0.260263 |
| *E. coli* AIC221 | 0.733874 | 0.719559 | 0.673859 | 0.653163 | 0.681991 | 0.617147 | 0.469486 |
| *E. coli* AIC222 | 0.737407 | 0.720051 | 0.672798 | 0.6632 | 0.626752 | 0.607942 | 0.452432 |
| *K. pneumoniae* ATCC13883 | 0.527278 | 0.486858 | 0.260868 | 0.430102 | 0.058973 | 0.333547 | 0.159662 |
| *A. baumannii* ATCC19606 | 0.726801 | 0.703749 | 0.687577 | 0.676496 | 0.696205 | 0.635361 | 0.510874 |
| Methicillin-resistant *S. aureus* ATCC BAA-1556 | 0.525053 | 0.489835 | 0.462725 | 0.460328 | 0.309507 | 0.428855 | 0.348216 |
| Vancomycin-resistant *E. faecalis* ATCC700802 | 0.107576 | 0.047122 | 0.141981 | 0.079331 | 0.169632 | -0.08813 | 0.200453 |
| Vancomycin-resistant *E. faecium* ATCC700221 | 0.646947 | 0.569296 | 0.484717 | 0.524636 | 0.444265 | 0.531174 | 0.468919 |
| **Average** | **0.593923** | **0.556458** | **0.514391** | **0.523119** | **0.457529** | **0.469322** | **0.371911** |

**Table S8. R-squared of single APEX variants on CV set.** We divided our in-house peptide dataset into a CV set for hyperparameter tuning and ML model selection, and an independent dataset to evaluate prediction performance. The table shows the average species-specific prediction performance of 5-fold CV in terms of R-squared for various APEX variants, including the original APEX (*i.e.*, single APEX), APEX without multitask constraint, APEX without using public AMP data during training, and APEX without multitask constraint and public AMP data during training.

| **Strain** | **Single APEX** | **Single APEX without multitask constraint** | **Single APEX without public AMP data** | **Single APEX without multitask constraint and public AMP data** |
|---|---|---|---|---|
| *E. coli* ATCC11775 | 0.519899 | 0.540226 | 0.515468 | 0.504094 |
| *P. aeruginosa* PAO1 | 0.458351 | 0.411795 | 0.418864 | 0.328494 |
| *P. aeruginosa* PA14 | 0.415884 | 0.33248 | 0.386427 | 0.253563 |
| *S. aureus* ATCC12600 | 0.443486 | 0.418941 | 0.378119 | 0.391915 |
| *E. coli* AIC221 | 0.527619 | 0.509886 | 0.468906 | 0.459699 |
| *E. coli* AIC222 | 0.578155 | 0.586917 | 0.547914 | 0.559102 |
| *K. pneumoniae* ATCC13883 | 0.078356 | 0.110211 | 0.12016 | 0.078548 |
| *A. baumannii* ATCC19606 | 0.597275 | 0.597209 | 0.554144 | 0.57716 |
| Methicillin-resistant *S. aureus* ATCC BAA-1556 | 0.442817 | 0.500468 | 0.432583 | 0.45067 |
| Vancomycin-resistant *E. faecalis* ATCC700802 | -0.46429 | -0.18286 | -0.48424 | 0.021012 |
| Vancomycin-resistant *E. faecium* ATCC700221 | 0.460972 | 0.512912 | 0.479714 | 0.52491 |
| **Average** | **0.368956** | **0.39438** | **0.347096** | **0.377197** |

**Table S9. Pearson correlation of single APEX variants on CV set.** We divided our in-house peptide dataset into a CV set for hyperparameter tuning and ML model selection, and an independent dataset to evaluate prediction performance. The table shows the average species-specific prediction performance of 5-fold CV in terms of Pearson correlation for various APEX variants, including the original APEX (i.e., single APEX), APEX without multitask constraint, APEX without using public AMP data during training, and APEX without multitask constraint and public AMP data during training.

| Strain | Single APEX | Single APEX without multitask constraint | Single APEX without public AMP data | Single APEX without multitask constraint and public AMP data |
|---|---|---|---|---|
| *E. coli* ATCC11775 | 0.722816 | 0.745399 | 0.72278 | 0.724131 |
| *P. aeruginosa* PAO1 | 0.683082 | 0.660926 | 0.657109 | 0.616645 |
| *P. aeruginosa* PA14 | 0.654815 | 0.603326 | 0.633283 | 0.563075 |
| *S. aureus* ATCC12600 | 0.672914 | 0.673138 | 0.62627 | 0.650051 |
| *E. coli* AIC221 | 0.728434 | 0.716857 | 0.691317 | 0.687469 |
| *E. coli* AIC222 | 0.761297 | 0.76824 | 0.744002 | 0.752025 |
| *K. pneumoniae* ATCC13883 | 0.401615 | 0.419686 | 0.408857 | 0.390867 |
| *A. baumannii* ATCC19606 | 0.774607 | 0.776853 | 0.749409 | 0.767272 |
| Methicillin-resistant *S. aureus* ATCC BAA-1556 | 0.680613 | 0.717745 | 0.661616 | 0.67801 |
| Vancomycin-resistant *E. faecalis* ATCC700802 | 0.056318 | 0.157377 | 0.010612 | 0.275715 |
| Vancomycin-resistant *E. faecium* ATCC700221 | 0.697111 | 0.721689 | 0.699877 | 0.734134 |
| **Average** | **0.621238** | **0.63284** | **0.600467** | **0.621763** |

**Table S10. Spearman correlation of single APEX variants on CV set.** We divided our in-house peptide dataset into a CV set for hyperparameter tuning and ML model selection, and an independent dataset to evaluate prediction performance. The table shows the average species-specific prediction performance of 5-fold CV in terms of Spearman correlation for various APEX variants, including the original APEX (i.e., single APEX), APEX without multitask constraint, APEX without using public AMP data during training, and APEX without multitask constraint and public AMP data during training.

| **Strain** | **Single APEX** | **Single APEX without multitask constraint** | **Single APEX without public AMP data** | **Single APEX without multitask constraint and public AMP data** |
|---|---|---|---|---|
| *E. coli* ATCC11775 | 0.66331 | 0.624219 | 0.6493 | 0.640146 |
| *P. aeruginosa* PAO1 | 0.611234 | 0.556474 | 0.576812 | 0.575338 |
| *P. aeruginosa* PA14 | 0.625266 | 0.578997 | 0.609458 | 0.544803 |
| *S. aureus* ATCC12600 | 0.484758 | 0.478392 | 0.456472 | 0.47019 |
| *E. coli* AIC221 | 0.719559 | 0.659801 | 0.683423 | 0.661957 |
| *E. coli* AIC222 | 0.720051 | 0.686905 | 0.691819 | 0.688436 |
| *K. pneumoniae* ATCC13883 | 0.486858 | 0.370328 | 0.435828 | 0.396926 |
| *A. baumannii* ATCC19606 | 0.703749 | 0.687933 | 0.703927 | 0.705705 |
| Methicillin-resistant *S. aureus* ATCC BAA-1556 | 0.489835 | 0.469302 | 0.498565 | 0.508692 |
| Vancomycin-resistant *E. faecalis* ATCC700802 | 0.047122 | 0.150755 | 0.055165 | 0.242589 |
| Vancomycin-resistant *E. faecium* ATCC700221 | 0.569296 | 0.586361 | 0.577632 | 0.59948 |
| **Average** | **0.556458** | **0.53177** | **0.539855** | **0.548569** |

**Table S11. Hyperparameters of top eight APEX ranked by R-squared on CV set.**

| Top eight APEX models | number of RNN layers | n | m | λ_l2 | λ_(multitask_constraint) | λ_BCE |
|---|---|---|---|---|---|---|
| 1 | 3 | 128 | 2048 | 1.00E-05 | 0.1 | 1 |
| 2 | 3 | 256 | 2048 | 1.00E-06 | 0.1 | 1 |
| 3 | 2 | 128 | 512 | 1.00E-05 | 0.01 | 1 |
| 4 | 3 | 128 | 512 | 1.00E-05 | 0.001 | 1 |
| 5 | 2 | 128 | 2048 | 1.00E-06 | 0 | 1 |
| 6 | 3 | 256 | 512 | 1.00E-06 | 0 | 1 |
| 7 | 2 | 128 | 2048 | 1.00E-05 | 0.01 | 1 |
| 8 | 2 | 256 | 2048 | 1.00E-06 | 0.1 | 1 |

**Table S12. R-squared of various ML models on independent set.** We divided our in-house peptide dataset into a CV set for hyperparameter tuning and ML model selection, and an independent dataset to evaluate prediction performance. The table shows the species-specific prediction performance in terms of R-squared for various ML models that were trained on the CV set and evaluated on the independent set. RF: random forest; GBDT: gradient boosting decision tree; ExtraTree: extra-tree regressor; ElasticNet: elastic net; LinearSVR: linear support vector regression.

| **Strain** | **Ensemble APEX v2** | **Ensemble APEX v1** | **Single APEX** | **RF** | **GBDT** | **ExtraTree** | **ElasticNet** | **LinearSVR** |
|---|---|---|---|---|---|---|---|---|
| *E. coli* ATCC11775 | 0.718159 | 0.691395 | 0.668556 | 0.63738 | 0.648213 | 0.551901 | 0.520619 | 0.401562 |
| *P. aeruginosa* PAO1 | 0.58922 | 0.550826 | 0.542209 | 0.585178 | 0.437675 | 0.583725 | 0.444805 | -0.30991 |
| *P. aeruginosa* PA14 | 0.577267 | 0.544865 | 0.523449 | 0.525311 | 0.469315 | 0.541576 | 0.464316 | 0.082555 |
| *S. aureus* ATCC12600 | 0.654147 | 0.65415 | 0.613936 | 0.498575 | 0.496227 | 0.286083 | 0.309557 | 0.254665 |
| *E. coli* AIC221 | 0.454362 | 0.423644 | 0.448303 | 0.380448 | 0.35911 | 0.338844 | 0.330079 | -0.07492 |
| *E. coli* AIC222 | 0.659494 | 0.642316 | 0.636876 | 0.540003 | 0.476026 | 0.456006 | 0.400548 | 0.208218 |
| *K. pneumoniae* ATCC13883 | 0.388908 | 0.380317 | 0.439253 | 0.134236 | 0.174184 | -0.16017 | 0.095197 | -0.17792 |
| *A. baumannii* ATCC19606 | 0.694184 | 0.697003 | 0.687141 | 0.664341 | 0.625813 | 0.627206 | 0.541836 | 0.250524 |
| Methicillin-resistant *S. aureus* ATCC BAA-1556 | 0.725662 | 0.739452 | 0.60538 | 0.578394 | 0.587113 | 0.221953 | 0.325214 | -0.12777 |
| Vancomycin-resistant *E. faecalis* ATCC700802 | 0.002173 | -0.07186 | 0.304024 | -0.04891 | -0.09351 | -0.09296 | -0.01733 | -0.01604 |
| Vancomycin-resistant *E. faecium* ATCC700221 | 0.538558 | 0.472891 | 0.514716 | 0.448163 | 0.375496 | 0.28212 | 0.34944 | 0.130954 |
| **Average** | **0.545648** | **0.520455** | **0.543986** | **0.449374** | **0.414151** | **0.330572** | **0.342208** | **0.056538** |

**Table S13. Pearson correlation of various ML models on independent set.** We divided our in-house peptide dataset into a CV set for hyperparameter tuning and ML model selection, and an independent dataset to evaluate prediction performance. The table shows the species-specific prediction performance in terms of Pearson correlation for various ML models that were trained on the CV set and evaluated on the independent set. RF: random forest; GBDT: gradient boosting decision tree; ExtraTree: extra-tree regressor; ElasticNet: elastic net; LinearSVR: linear support vector regression.

| Strain | Ensemble APEX v2 | Ensemble APEX v1 | Single APEX | RF | GBDT | ExtraTree | ElasticNet | LinearSVR |
|---|---|---|---|---|---|---|---|---|
| *E. coli* ATCC11775 | 0.847805 | 0.832404 | 0.824249 | 0.800211 | 0.808419 | 0.744944 | 0.722128 | 0.723382 |
| *P. aeruginosa* PAO1 | 0.790273 | 0.769464 | 0.775476 | 0.771463 | 0.722799 | 0.771821 | 0.672709 | 0.58249 |
| *P. aeruginosa* PA14 | 0.767182 | 0.746836 | 0.739504 | 0.725186 | 0.708727 | 0.736129 | 0.69213 | 0.593574 |
| *S. aureus* ATCC12600 | 0.810676 | 0.81136 | 0.784346 | 0.70632 | 0.714723 | 0.551515 | 0.55703 | 0.598799 |
| *E. coli* AIC221 | 0.688587 | 0.671029 | 0.693506 | 0.62584 | 0.654544 | 0.597245 | 0.580406 | 0.466094 |
| *E. coli* AIC222 | 0.813293 | 0.803721 | 0.802091 | 0.736241 | 0.723161 | 0.676554 | 0.632948 | 0.607374 |
| *K. pneumoniae* ATCC13883 | 0.633471 | 0.627873 | 0.67345 | 0.37494 | 0.451051 | 0.185642 | 0.325999 | 0.404414 |
| *A. baumannii* ATCC19606 | 0.841729 | 0.842787 | 0.84471 | 0.818252 | 0.805014 | 0.796043 | 0.764364 | 0.674816 |
| Methicillin-resistant *S. aureus* ATCC BAA-1556 | 0.859695 | 0.865534 | 0.786457 | 0.772415 | 0.78984 | 0.598027 | 0.578747 | 0.580319 |
| Vancomycin-resistant *E. faecalis* ATCC700802 | 0.21569 | 0.101714 | 0.576916 | 0.119566 | -0.51239 | 0 | 0 | 0.193324 |
| Vancomycin-resistant *E. faecium* ATCC700221 | 0.736427 | 0.690436 | 0.741292 | 0.761104 | 0.642366 | 0.581336 | 0.684935 | 0.387822 |
| Average | 0.727712 | 0.705742 | 0.749272 | 0.655595 | 0.59166 | 0.567205 | 0.564672 | 0.528401 |

**Table S14. Spearman correlation of various ML models on independent set.** We divided our in-house peptide dataset into a CV set for hyperparameter tuning and ML model selection, and an independent dataset to evaluate prediction performance. The table shows the species-specific prediction performance in terms of Spearman correlation for various ML models that were trained on the CV set and evaluated on the independent set. RF: random forest; GBDT: gradient boosting decision tree; ExtraTree: extra-tree regressor; ElasticNet: elastic net; LinearSVR: linear support vector regression.

| **Strain** | **Ensemble APEX v2** | **Ensemble APEX v1** | **Single APEX** | **RF** | **GBDT** | **ExtraTree** | **ElasticNet** | **LinearSVR** |
|---|---|---|---|---|---|---|---|---|
| *E. coli* ATCC11775 | 0.742143 | 0.738599 | 0.695429 | 0.70242 | 0.686571 | 0.675388 | 0.618169 | 0.623012 |
| *P. aeruginosa* PAO1 | 0.701432 | 0.671522 | 0.620927 | 0.631221 | 0.673711 | 0.592132 | 0.545197 | 0.481929 |
| *P. aeruginosa* PA14 | 0.725533 | 0.695124 | 0.619324 | 0.665001 | 0.613896 | 0.63482 | 0.604422 | 0.484608 |
| *S. aureus* ATCC12600 | 0.578662 | 0.528803 | 0.52903 | 0.435649 | 0.497982 | 0.518397 | 0.419481 | 0.407387 |
| *E. coli* AIC221 | 0.707444 | 0.707646 | 0.651341 | 0.670901 | 0.670596 | 0.676166 | 0.614667 | 0.583788 |
| *E. coli* AIC222 | 0.721146 | 0.711075 | 0.666143 | 0.723491 | 0.626716 | 0.638027 | 0.609947 | 0.570258 |
| *K. pneumoniae* ATCC13883 | 0.603606 | 0.581167 | 0.53605 | 0.28463 | 0.483702 | 0.190172 | 0.379504 | 0.31371 |
| *A. baumannii* ATCC19606 | 0.683889 | 0.664927 | 0.661991 | 0.673022 | 0.625251 | 0.695621 | 0.624478 | 0.600003 |
| Methicillin-resistant *S. aureus* ATCC BAA-1556 | 0.474364 | 0.456122 | 0.482894 | 0.44037 | 0.418149 | 0.570131 | 0.418258 | 0.447617 |
| Vancomycin-resistant *E. faecalis* ATCC700802 | 0.238607 | 0.206435 | 0.361932 | 0.146958 | -0.4407 | 0 | 0 | 0.184652 |
| Vancomycin-resistant *E. faecium* ATCC700221 | 0.495449 | 0.436833 | 0.3834 | 0.427797 | 0.359027 | 0.333684 | 0.392771 | 0.397954 |
| **Average** | **0.60657** | **0.581659** | **0.564406** | **0.527405** | **0.474082** | **0.502231** | **0.475172** | **0.463174** |

**Table S15. Pearson correlation of various ML models on human encrypted peptides.** The human encrypted peptides from Torres et al.[1] constitute a subset of our in-house peptide dataset. We treated this subset as the test data and excluded it from ML model training. The table shows species-specific prediction performance in terms of Pearson correlation for various ML models.

| Strain | Ensemble APEX v2 | Ensemble APEX v1 | Single APEX | RF | GBDT | ExtraTree | ElasticNet | LinearSVR | Torres *et al.* |
|---|---|---|---|---|---|---|---|---|---|
| *E. coli* ATCC11775 | 0.482757 | 0.496734 | 0.494723 | 0.435897 | 0.365328 | 0.407247 | 0.428736 | 0.320893 | 0.50776 |
| *P. aeruginosa* PAO1 | 0.710869 | 0.686719 | 0.579761 | 0.731061 | 0.679557 | 0.699833 | 0.569554 | 0.143789 | 0.582418 |
| *P. aeruginosa* PA14 | 0.734694 | 0.73291 | 0.671569 | 0.749065 | 0.68225 | 0.749057 | 0.725144 | 0.254412 | 0.712036 |
| *S. aureus* ATCC12600 | 0.126253 | 0.186218 | 0.199264 | 0.076771 | 0.020007 | 0.07906 | -0.01304 | -0.26126 | 0.42482 |
| *E. coli* AIC221 | 0.403527 | 0.404244 | 0.389027 | 0.521815 | 0.413622 | 0.483885 | 0.398567 | 0.200215 | 0.311953 |
| *E. coli* AIC222 | 0.653576 | 0.655583 | 0.741654 | 0.680734 | 0.606786 | 0.762781 | 0.411687 | 0.316373 | 0.433937 |
| *K. pneumoniae* ATCC13883 | 0.460277 | 0.466471 | 0.507818 | 0.456984 | 0.295749 | -0.0989 | 0.22667 | 0.254919 | 0.282847 |
| *A. baumannii* ATCC19606 | 0.773425 | 0.773293 | 0.73518 | 0.72433 | 0.760228 | 0.753661 | 0.510162 | 0.601455 | 0.430196 |
| **Average** | **0.543172** | **0.550272** | **0.539874** | **0.547082** | **0.477941** | **0.479578** | **0.407185** | **0.228849** | **0.460746** |

**Table S16. Spearman correlation of various ML models on human encrypted peptides.** The human encrypted peptides from Torres et al.[1] constitute a subset of our in-house peptide dataset. We treated this subset as the test data and excluded it from ML model training. The table shows species-specific prediction performance in terms of Spearman correlation for various ML models.

| **Strain** | **Ensemble APEX v2** | **Ensemble APEX v1** | **Single APEX** | **RF** | **GBDT** | **ExtraTree** | **ElasticNet** | **LinearSVR** | **Torres et al.** |
|---|---|---|---|---|---|---|---|---|---|
| *E. coli* ATCC11775 | 0.523727 | 0.535223 | 0.506701 | 0.492004 | 0.370203 | 0.473523 | 0.440344 | 0.220609 | 0.497825 |
| *P. aeruginosa* PAO1 | 0.719366 | 0.718463 | 0.622332 | 0.673903 | 0.575407 | 0.648483 | 0.626982 | 0.03871 | 0.615923 |
| *P. aeruginosa* PA14 | 0.76658 | 0.760304 | 0.657554 | 0.657068 | 0.683707 | 0.662736 | 0.688632 | 0.225784 | 0.721803 |
| *S. aureus* ATCC12600 | 0.254791 | 0.233065 | 0.259042 | 0.244744 | 0.081066 | 0.177344 | 0.025934 | -0.20224 | 0.4654 |
| *E. coli* AIC221 | 0.403575 | 0.397753 | 0.346453 | 0.491014 | 0.378764 | 0.376929 | 0.409659 | 0.137742 | 0.31409 |
| *E. coli* AIC222 | 0.584407 | 0.582971 | 0.583851 | 0.589457 | 0.487067 | 0.636713 | 0.55179 | 0.17471 | 0.477801 |
| *K. pneumoniae* ATCC13883 | 0.420755 | 0.417876 | 0.315581 | 0.313113 | 0.332973 | -0.10474 | 0.440967 | 0.066277 | 0.293077 |
| *A. baumannii* ATCC19606 | 0.634678 | 0.64788 | 0.602808 | 0.6092 | 0.62634 | 0.611794 | 0.513264 | 0.458603 | 0.455221 |
| Average | 0.538485 | 0.536692 | 0.48679 | 0.508813 | 0.441941 | 0.435348 | 0.462197 | 0.140025 | 0.480142 |

**Table S17. Cytotoxic activity of AEPs and MEPs.** The cytotoxic activity was expressed in terms of $CC_{50}$ values (μmol $L^{-1}$), *i.e.*, cytotoxic concentration values needed to damage 50% of the HEK293T cells present in each condition. The values were estimated by non-linear regressions based on the screen of all active AEPs and MEPs at concentrations from 8 to 128 μmol $L^{-1}$, to ensure coverage of all tested antimicrobial activity concentrations. The experiments were done in three independent biological replicates with two technical replicates within each biological replicate. The therapeutic index (TI) was calculated to show the margin of safety obtained by comparing the lowest MIC values (μmol $L^{-1}$) obtained in the antimicrobial activity assays to the $CC_{50}$ values of each active AEP or MEP.

| Peptide | $CC_{50}$ (μmol $L^{-1}$) | MIC (μmol $L^{-1}$) | TI | Peptide | $CC_{50}$ (μmol $L^{-1}$) | MIC (μmol $L^{-1}$) | TI |
|---|---|---|---|---|---|---|---|
| Equusin-1 | >128 | 1 | >128 | Megalocerin-1 | >128 | 8 | >16 |
| Hesperelin-1 | >128 | 2 | >64 | Pinguinusin-1 | >128 | 4 | >32 |
| Elephasin-1 | >128 | 4 | >32 | Ursusin-1 | >128 | 64 | >2 |
| Arctodutin-1 | >128 | 2 | >64 | Elephasin-2 | >128 | 1 | >128 |
| Arctoterin-1 | >128 | 64 | >2 | Mammuthusin-4 | >128 | 2 | >64 |
| Lophiosin-1 | 68.02 | 16 | >8 | Psephotellin-1 | >128 | 16 | >8 |
| Mammutin-1 | >128 | 2 | >64 | Eudyptin-1 | >128 | 64 | >2 |
| Ararin-1 | >128 | 32 | >4 | Paleopropin-2 | >128 | 16 | >8 |
| Mylodonin-1 | >128 | 8 | >16 | Hydrodamin-2 | >128 | 8 | >16 |
| Mammuthusin-1 | >128 | 4 | >32 | Hydrodamin-3 | >128 | 16 | >8 |
| Paleopropin-1 | >128 | 32 | >4 | Hesperelin-4 | >128 | 64 | >2 |
| Bisonin-1 | >128 | 16 | >8 | Mylodonin-2 | >128 | 32 | >4 |
| Hesperelin-2 | >128 | 8 | >16 | Anomalopterin-2 | >128 | 32 | >4 |
| Equusin-2 | >128 | 8 | >16 | Equusin-3 | >128 | 4 | >32 |
| Mammuthusin-2 | >128 | 32 | >4 | Bisonin-2 | >128 | 8 | >16 |
| Mammuthusin-3 | >128 | 16 | >8 | Mammuthusin-5 | >128 | 32 | >4 |
| Hydrodamin-1 | >128 | 4 | >32 | Smilodin-1 | >128 | 64 | >2 |
| Xenothrixin-1 | 70.77 | 8 | 8.84 | Mylodonin-3 | >128 | 16 | >8 |
| Hesperelin-3 | >128 | 64 | >2 | Mylodonin-4 | >128 | 32 | >4 |
| Ararin-2 | >128 | 16 | >8 | Equusin-4 | >128 | 16 | >8 |
| Anomalopterin-1 | >128 | 64 | >2 | | | | |

**Caption for Data S1 to S3**

**Data S1**. List of EPs predicted by APEX to have median MIC ≤80 μmol $L^{-1}$.

**Data S2**. Information of EPs identified by APEX and validated experimentally. 69 EPs predicted by APEX and selected based on our criteria were synthesized and validated. Their sequences, antimicrobial activities and annotations are provided.

**Data S3**. Information of EPs identified by our earlier AMP scoring function and validated experimentally. 49 EPs predicted by our AMP scoring function[1] were synthesized and validated. Their sequences and antimicrobial activities are provided.
